# Light-guided actin polymerization drives directed motility in protocells

**DOI:** 10.1101/2024.10.14.617543

**Authors:** Hideaki T. Matsubayashi, Shiva Razavi, Yuhei O. Tahara, Oghosa H. Akenuwa, T. Willow Rock, Daichi Nakajima, Rina Otsuka-Yamaguchi, Hideki Nakamura, Daniel A. Kramer, Tomoaki Matsuura, Baoyu Chen, Christopher T. Lee, Makoto Miyata, Satoshi Murata, Shinichiro M. Nomura, Takanari Inoue

## Abstract

Motility is a hallmark of life’s dynamic processes, enabling cells to actively chase prey, repair wounds, and shape organs. Recreating these intricate behaviors using well-defined molecules remains a major challenge at the intersection of biology, physics, and molecular engineering. Although the polymerization force of the actin cytoskeleton is characterized as a primary driver of cell motility, building a minimal platform to test this process in protocellular systems has proven elusive. The difficulty lies in the daunting task of distilling key components from motile cells and integrating them into model membranes in a physiologically relevant manner. To address this, we developed a method to optically control actin polymerization with high spatiotemporal precision within cell-mimetic lipid vesicles known as giant unilamellar vesicles (GUVs). Within these active protocells, the reorganization of actin networks triggered outward membrane extensions as well as the unidirectional movement of GUVs at speeds of up to 0.43 µm/min, within the range of adherent mammalian cells. Notably, our findings reveal the requirements of both branched and linear actin networks for efficient membrane protrusions. This approach offers a powerful platform for unraveling the intricacies of cell migration, designing synthetic cells with active morphodynamics, and advancing bioengineering applications, such as self-propelled delivery systems and autonomous tissue-like materials.

## Introduction

Motility is a fundamental property of cells across biological systems, from bacteria to large animals (*1–6*). By transcending the randomness and viscosity of the microscopic world, cells move toward their destination by extending membrane protrusions and by clutching onto the extracellular environment. In amoeboid cell migration, these processes are universally galvanized by actin polymerization (*7–12*). While hundreds of biomolecules are identified to be required for cell motility (*4, 6, 13*), a minimal list of biomolecular components sufficient for cell motility remains elusive. This primarily reflects the inherent difficulty of eliminating all biomolecules beyond the minimal set in the context of living cells.

Cell migration is driven by the coordination of forward movement, rear contraction, and adhesion/deadhesion of the cell edges (*4, 6, 14–16*). Among these processes, actin polymerization at the leading edge is a strong candidate for driving overall forward displacement. For instance, *Dictyostelium*, fish keratocyte, and human neutrophil retain their motility even after myosin contractility is compromised genetically (*17, 18*) or pharmacologically (*19, 20*). Cells also manage to migrate even without strong adhesions especially in confined conditions (*21, 22*). Conversely, we and others have shown that synthetic activation of the Rho family GTPase Rac1, an upstream activator of the actin cytoskeleton, induces directional cell movements toward the site of stimulation (*23, 24*). However, it is still unclear if Rac1-mediated actin polymerization alone is sufficient for cell motility in cellular settings, as Rac1 crosstalks with other Rho small GTPases such as RhoA and triggers other cytoskeletal activities including actomyosin contraction (*25*).

In a different context, molecules necessary for force generation by actin polymerization have been identified based on the motility reconstitution assay (*26–36*). These reports demonstrated that a mixture of purified cytoskeletal proteins, such as actin, Arp2/3, profilin, cofilin, and capping protein, can reconstitute the motility of *Listeria monocytogenes* bacteria using microspheres coated with actin nucleation-promoting factors (NPFs). Furthermore, a number of studies were conducted to reconstitute actin (*37, 38*) and other cytoskeletons (*39–41*) inside cell-sized membrane-bound liposomes. These studies demonstrated that actin polymerization generates membrane deformations (*42–45*), revealed the roles of actin-binding proteins (*46–52*), and further showed that internal reactions can be regulated by light and external environmental factors (*52–55*). However, unidirectional motion by the cytoskeletal polymerization force has remained challenging. Moreover, non-invasive and non-constitutive—thus more physiological—induction of actin polymerization has yet to be achieved, particularly within cell-sized compartments that maintain proper membrane configuration. Building on these findings, we developed a minimal membrane-bound system in which actin polymerization can be locally controlled and tested whether such an artificial cytoskeletal activity is sufficient to induce cell-like motility.

## Results

### Reversible and asymmetric light control of protein localization inside GUVs

To achieve *in vitro* reconstitution of cell motility, we previously developed active protocells where chemically inducible dimerization (CID) tools were used to polymerize actin based on a bacterial NPF inside giant unilamellar vesicles (GUVs) (*56*). While the previous system efficiently produced actin polymerization, outward membrane extension necessary for driving movement was not observed. We reasoned that this was due to a lack of fast turnover of NPF recruitment to the membranes. To aim for a physiologically relevant generation of protrusive force exerted against membranes, we adopted a light-inducible protein dimerization tool for controlling the localization of proteins of interest. Among optogenetic dimerization tools (*57–59*), we chose iLID-SspB (*60*) as its second-scale reversible binding kinetics match the turnover of leading-edge polarization (Supplementary Text 1), its blue-light activation is compatible with standard microscope setups, and its relatively small molecular size facilitates expression and purification of recombinant proteins.

We first set out to test light-induced dimerization of purified iLID and SspB in GUVs. In our construct, iLID was membrane-tethered, and blue light exposure triggered the Jα helix to extend, revealing the binding site for SspB. In the dark, iLID returned to its closed state, releasing SspB (Fig. 1A). We visualized the localization and the light responsiveness of the proteins by fusing iLID with YFP and SspB with mCherry (fig. S1A). To anchor iLID to the GUV membrane, we tested four different protein-lipid interaction strategies to mediate light-induced translocation, including coordination bonding, electrostatic, strong non-covalent, and covalent interactions (Fig. 1A, fig. S2, Supplementary Text 2) (*61, 62*). Among these, the SNAP-tag/benzylguanine (BG)-conjugated lipid pair provided efficient iLID anchoring with minimal nonspecific membrane binding of SspB, and was therefore used for subsequent experiments.

**Fig. 1.**
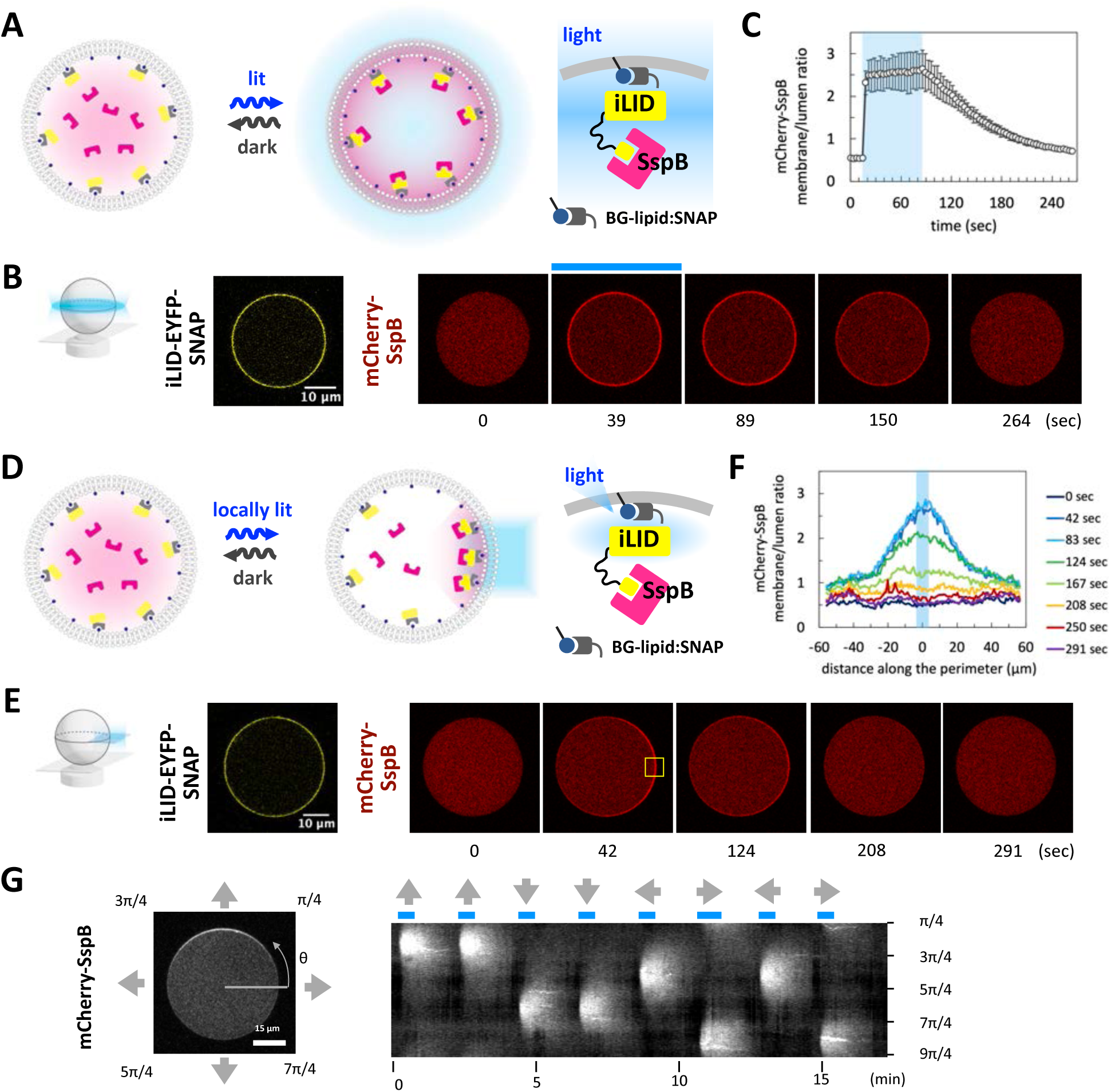
Reversible and asymmetric control of protein localization with iLID-SspB within GUVs. (A) Schematic representation of iLID-SspB reaction with global light stimulation. iLID was tethered on the membrane via the interaction between SNAP-tag and benzylguanine-conjugated lipids. Upon blue light illumination, Jα helix of iLID extends to expose the binding site for SspB, whereas in the dark, iLID reverses to its closed state and releases SspB. (B) Representative images of iLID-YFP-SNAP and mCherry-SspB. Scale bar, 10 µm. Blue bar indicates the period of blue light illumination. (C) Time course of membrane/lumen ratio of mCherry-SspB signal (n=5). Error bars indicate 95% CI. Blue area indicates the period of blue light stimulation. (D) Schematic representation of iLID-SspB reaction with local light stimulation. (E) Representative images of mCherry-SspB. Scale bar, 10 µm. Yellow box indicates the area of blue light illumination. (F) Membrane/lumen ratio of mCherry-SspB for the representative image shown. The distance is measured from the center of blue light illumination along the perimeter of the GUV. (G) Representative image and kymograph of repetitive and reversible SspB translocation. Kymograph shows membrane signal of mCherry-SspB.

When GUVs were made with 2% BG-conjugated lipid in the inner leaflet, iLID localized on the membrane, while SspB remained in the lumen in the dark (Fig. 1B). Upon blue light illumination of the entire GUV, SspB translocated to the membrane within 4 seconds (Fig. 1B, C). After switching back to the dark, SspB gradually dissociated from the membrane with a half-time of 61.2 ± 13.7 seconds. The time scale is consistent with AsLOV2-based optogenetic tools measured in cells (*58*). These data confirmed that iLID-SspB enables reversible control of protein localization in GUVs.

Next, we tested the potential for spatially asymmetric control of iLID-SspB interactions. When light was applied locally to one side of the GUV, SspB accumulated at the illuminated site (Fig. 1D, E). In the dark, dissociation proceeded with a half-time of 41.8 ± 7.1 seconds (fig. S3A). Despite faster diffusion on artificial lipid membranes compared to cell membranes (*63–65*), localized light stimulus created an asymmetric pattern of SspB in cell-sized vesicles (Fig. 1E, F, see Supplementary Text 1). Furthermore, SspB precisely and repeatedly responded to the directional change of blue light illumination (Figs. 1G, S3B, C, and Movie S1). Collectively, these results demonstrated that the iLID-SspB system can direct asymmetric protein distribution at regions of interest on the GUV membranes.

### Light-guided control of ActA

Next, we sought to manipulate the actin cytoskeleton with the iLID-SspB system. Previous studies have established that the rate of actin polymerization depends on the surface density of NPFs (*26, 30, 35*) and that clustering of NPFs accelerates actin polymerization (*66, 67*), as further demonstrated by a reconstituted optogenetic VCA system (*68*). We have recently shown that chemically-induced membrane recruitment of ActA, an N-WASP homolog NPF derived from *L. monocytogenes*, induces actin polymerization on the membrane of GUVs, leading to symmetry breaking (*56, 69*). Thus, we first tested ActA with the iLID-SspB system. To identify an optimal optogenetic configuration, we tested four ActA-SspB constructs with different SspB variants and fusion orders (Supplementary Text 3). These constructs were evaluated based on actin polymerization activity and light-dependent membrane recruitment, leading us to select ActA-SspB_nano_-mCherry (fig. S4). When supplemented with actin and Arp2/3, light-induced global membrane recruitment of ActA-SspB led to the emergence of actin patches on the membrane, consistent with our previous findings with a chemical input (*56*) (fig. S5). With local light, the ActA distribution became asymmetric, albeit the directionality of actin pattern could not be maintained. In most cases, actin patches diffused randomly out of the illuminated region and the GUV lost actin directionality over time (fig. S6 and Movie S2).

### Reversible control of pVCA and light-guided actin polymerization

For robust maintenance of actin asymmetry, we aimed to rapidly depolymerize actin filaments in the non-illuminated area while strengthening actin polymerization in the illuminated region. To promote depolymerization and turnover, we incorporated factors that facilitate the actin turnover process, namely profilin, cofilin, and capping protein (*13, 33, 70, 71*). In ultracentrifugation assays, the combined addition of profilin, cofilin, and capping protein increased the pool of G-actin, supporting the notion that these factors enhance actin turnover (fig. S8A, B). To enhance polymerization in the illuminated regions within GUVs, we optimized the polymerization module. Among several NPF constructs evaluated, GST-tagged N-WASP pVCA (proline-rich region + VCA domain) showed the fastest actin polymerization in a bulk pyrene assay, which monitors actin polymerization through the fluorescence increase of pyrene-labeled actin. Notably, GST-pVCA maintained its activity when fused to SspB-mCherry, making GST-pVCA-SspB-mCherry (pVCA-SspB-mCherry) a suitable light-inducible activator for our GUV system (Fig. 2A, S1, S8C, D).

**Fig. 2.**
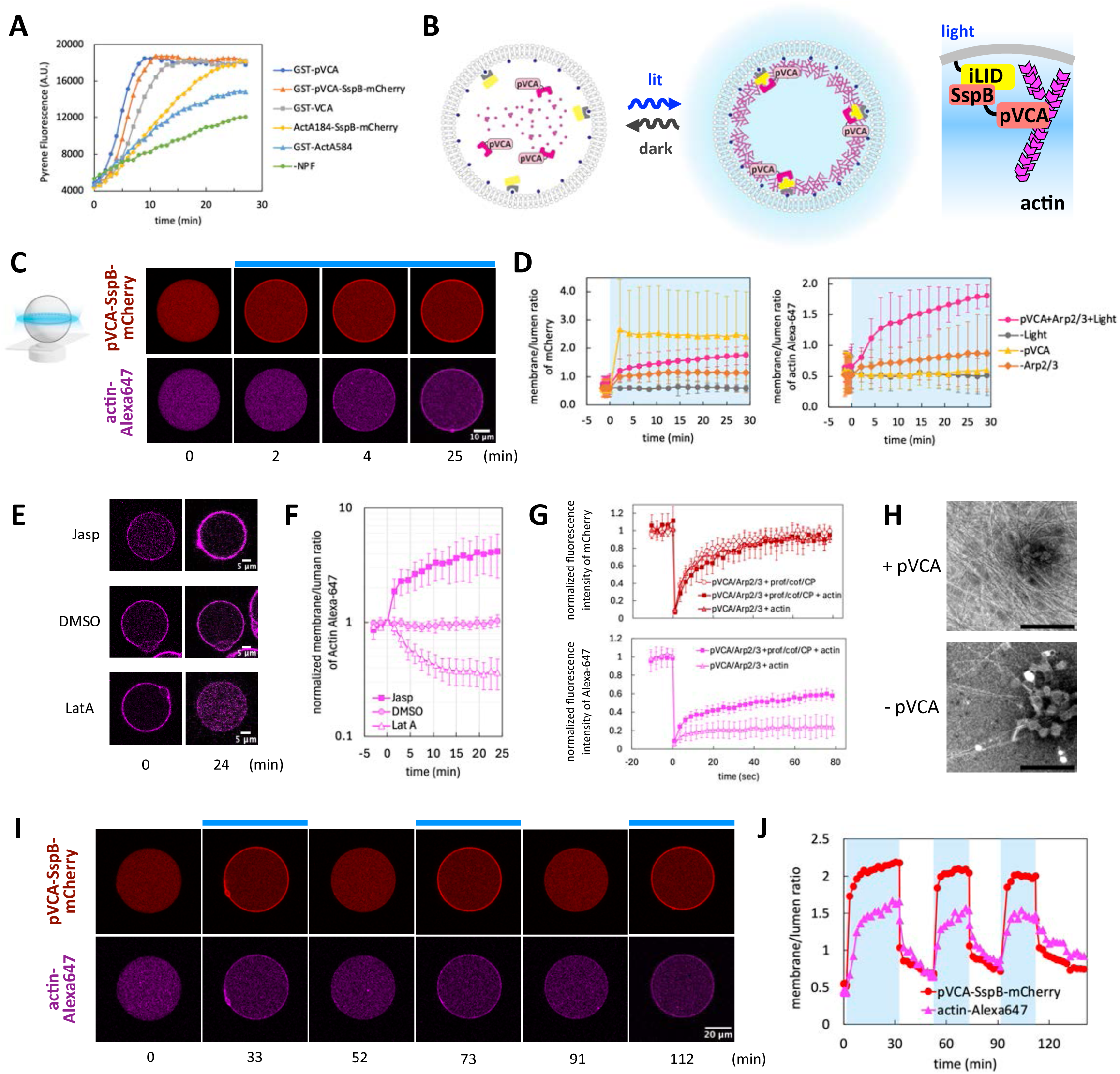
Reversible light control of actin cytoskeleton within GUVs. (A) Bulk pyrene actin polymerization assay with 1 µM actin (5% pyrene-labeled), 10 nM Arp2/3, and 100 nM NPFs. (-NPF) represents actin and Arp2/3 only condition. (B) Schematic representation of light-inducible actin polymerization. Translocation of pVCA-SspB-mCherry increases local concentration of pVCA on the membrane and enhances actin polymerization. (C) Representative images of pVCA-SspB-mCherry and Alexa 647 actin. Blue bar indicates the period of blue light illumination. (D) Time course of membrane/lumen ratio of mCherry and Alexa 647 actin in (C) and fig. S9. pVCA +Arp2/3+Light: full set of components (n=4). -Light: no blue light stimulation (n=3). -pVCA: pVCA-SspB-mCherry was replaced with mCherry-SspB (n=4). -Arp2/3: Arp2/3 was omitted (n=4). In (E)–(H), pVCA-SspB-mCherry was constitutively anchored to the membrane via SsrA-SNAP. (E) Pharmacological perturbation of membrane-associated actin. DMSO (n=8), Jasplakinolide (Jasp, n=10), Latrunculin A (LatA, n=9). (F) Time course of normalized membrane/lumen ratio of Alexa-647 actin in (E). The values were normalized by time=0. (G) FRAP analysis of membrane fluorescence of mCherry and Alexa 647 actin in fig. S11. The values were normalized by time=0. Sample sizes are listed in fig. S11A. (H) Electron microscopy (EM) images of actin filaments in the presence and absence of pVCA. Scale bar indicates 500 nm. (I) Representative images of reversible actin polymerization. Blue bars indicate the period of blue light illumination. (J) Time course of membrane/lumen ratio of pVCA-SspB-mCherry and Alexa 647-labeled actin in (F). (D, F, G) Error bars indicate 95% CI.

We then encapsulated this light-inducible pVCA-SspB-mCherry with actin (10% Alexa-647 labeled), Arp2/3, profilin, cofilin, and capping protein in GUVs (Fig. 2B). Upon illumination, pVCA-SspB-mCherry rapidly translocated to the membrane, followed by the increase in membrane actin signal. Notably, the actin signal increased gradually over time with a delay of several minutes after pVCA translocation, consistent with light-induced actin polymerization (Fig. 2C, D, Movie S3). In contrast, in the absence of either pVCA or light stimulation, no significant change in membrane actin intensity was observed (Fig. 2D, fig. S9). When Arp2/3 was omitted, the membrane actin signal showed a slight increase over time. This suggested that the G-actin binding property of pVCA (*72, 73*) increased the local concentration of actin at the periphery of the membrane, albeit this effect was not as intense as Arp2/3-dependent actin assembly (Fig. 2D, fig. S9). These data showed that light-dependent membrane accumulation of pVCA drives Arp2/3-dependent actin enrichment on the GUV membrane.

Next, we employed pharmacological perturbations to probe the dynamics of membrane-associated actin. Specifically, Latrunculin A sequesters G-actin to inhibit polymerization, whereas Jasplakinolide stabilizes F-actin and suppresses depolymerization. To decouple actin dynamics from optogenetic kinetics, iLID-SNAP was replaced with SsrA-SNAP, in which SsrA constitutively binds SspB-fused pVCA independently of light (fig. S10A). Under these conditions, actin was readily observed on the membrane (Fig. 2E, fig. S10B, C). Upon Latrunculin A treatment, the membrane actin signal markedly decreased, confirming that the membrane actin signal derives from F-actin. Conversely, Jasplakinolide increased membrane actin fluorescence, revealing additional capacity for membrane actin accumulation beyond the steady-state level observed under DMSO conditions (Fig. 2E, F, fig. S10D–F). Together, these results suggest that the membrane-associated actin is maintained by a balance between ongoing actin assembly and disassembly.

To further characterize the dynamics of pVCA and actin on the GUV membrane, we performed fluorescence recovery after photobleaching (FRAP) analysis. pVCA-SspB-mCherry exhibited rapid and near-complete fluorescence recovery, indicating that membrane-associated pVCA remains highly mobile even during actin enrichment (Fig. 2G, fig. S11). In contrast, FRAP analysis of Alexa Fluor 647–labeled actin showed slower and partial fluorescence recovery, suggesting the presence of a less mobile F-actin fraction at the membrane (Fig. 2G, fig. S11). Notably, actin fluorescence recovery was significantly reduced in the absence of regulatory factors, profilin, cofilin, and capping protein (fig. S11A, E), indicating that membrane-associated actin exists in a dynamic steady state regulated by these factors.

To examine the ultrastructure of the actin network, we performed negative staining electron microscopy (EM). For this analysis, small unilamellar vesicles (SUVs; 50–150 nm in diameter) containing 2% benzylguanine-modified lipids (fig. S12A) were used to recruit pVCA-SspB-mCherry to their outer surface via SsrA-SNAP. EM images revealed dense actin filament networks (Fig. 2H, fig. S12B–D) with frequent Y-junctions, characteristic of Arp2/3-mediated branching (fig. S12B), providing structural evidence for Arp2/3-dependent actin polymerization.

Next, we tested the reversible actin regulation by pVCA-SspB-mCherry. As shown in Fig. 2I and J, both pVCA-SspB-mCherry and actin displayed reversible and repetitive changes in membrane intensity in response to blue light switching (Movie S4). Notably, once switched to the dark, pVCA-SspB-mCherry quickly dissociated from the membrane, whereas actin decreased its membrane intensity at a relatively slower rate (fig. S13). Taken together, these results suggested that the polymerization state is reversible in our system and that the balance between polymerization and depolymerization can be controlled by light.

### Asymmetric and spatially controlled actin polymerization

The asymmetric distribution of NPFs and actin polymerization are fundamental features of directed cell migration (*9, 74, 75*). Thus, we next tested whether locally targeted pVCA could establish asymmetric actin polymerization (Fig. 3A). As shown in Fig. 3B, C, and fig. S14A, pVCA showed asymmetric patterning toward the localized light stimulus (Movie S5). About 5 minutes after pVCA translocation, the actin signal also increased from the illuminated region and created an asymmetric pattern. Line scans along the GUV perimeter indicated that pVCA responded rapidly with a narrow peak, whereas actin accumulated more gradually and exhibited a broader distribution. These results suggest that actin polymerization requires a finite time to initiate and propagate, and once filaments are formed they remain associated with the membrane for a longer duration (fig. S14B). With continuous local illumination, both pVCA and actin maintained stable polarity for more than 100 minutes (Fig. 3B, C, S14A, B, Movie S5). Notably, at later time points, the actin signal appeared as two lines in the latter time points in the kymograph. This likely reflects reduced lateral diffusion of membrane-associated actin filaments, combined with gradual photobleaching of Alexa Fluor 647-labeled actin under repeated blue-light excitation. We next tested whether the direction of actin polymerization could be reversibly reprogrammed. After establishing an asymmetric actin pattern with a local light on the right side of the vesicle, we reversed it back to the dark mode and subsequently applied new light on the left. Both pVCA and actin rapidly re-polarized toward the new stimulation site (Fig. 3D, E, and Movie S6). Together, these results demonstrate that the light-inducible pVCA system enables reversible and spatially reconfigurable polarization of actin networks.

**Fig. 3.**
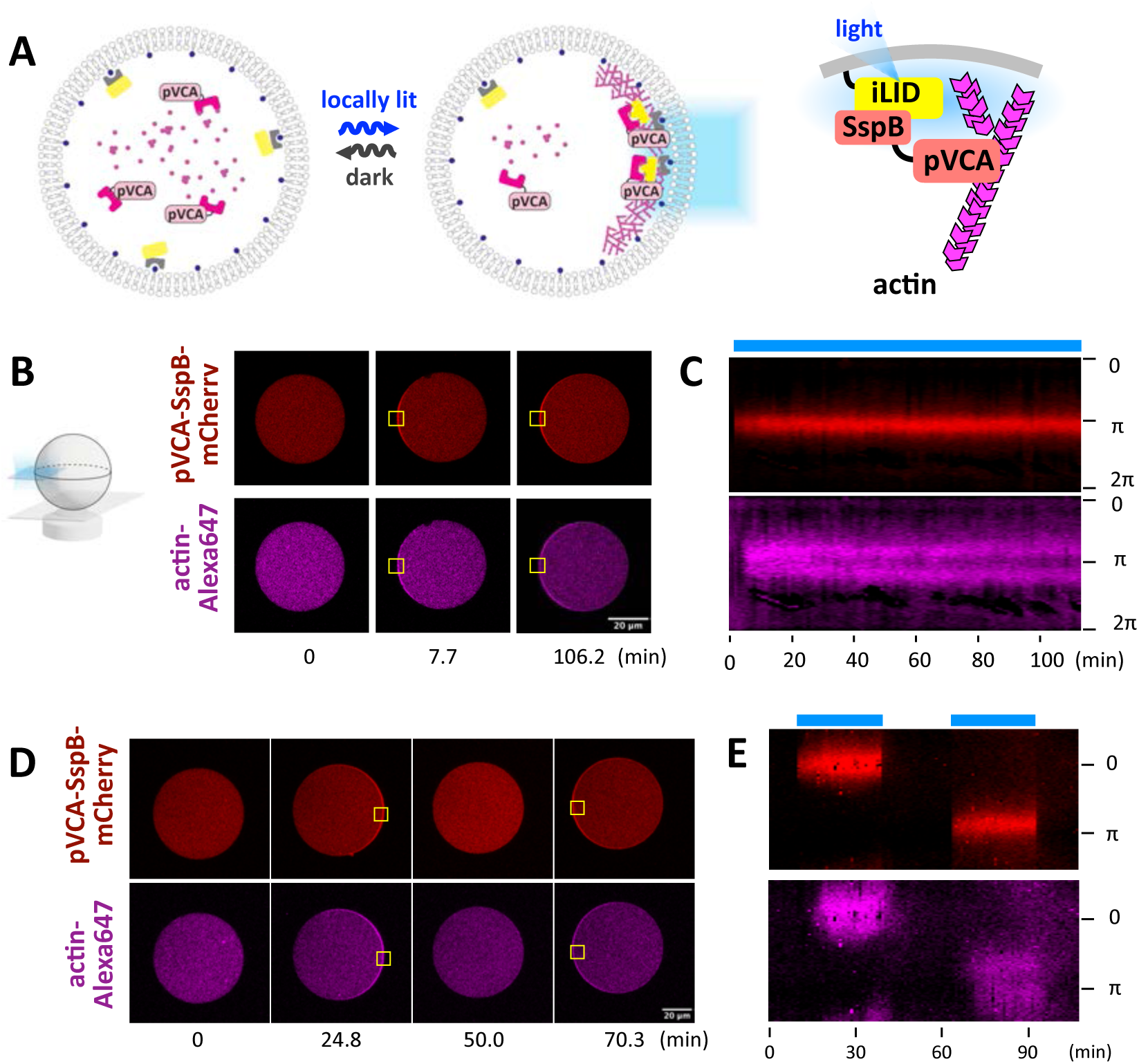
Asymmetric light control of actin cytoskeleton within GUVs. (A) Schematic representation of iLID-SspB reaction with local light stimulation. (B, D) Representative images of light-inducible asymmetric actin polymerization. Yellow boxes indicate the area of blue light illumination. (C, E) Kymograph of membrane pVCA-SspB and actin signals. Blue bars indicate the period of blue light illumination. Triangles indicate timepoints represented in (B, D).

Importantly, when locally illuminated, the front membrane of some GUVs moved toward the illuminated direction (fig. S14C), albeit occasionally (2 out of 7 instances). We speculate that characteristics inherent to the GUV system–such as surface-to-volume ratio of individual GUVs, surface density of pVCA, and the ratio of Arp2/3 to capping protein (*33, 76*), either individually or in combination, may determine whether productive protrusive behavior emerges.

### mDia1-driven actin polymerization drives membrane protrusions in cells

To enhance forward membrane protrusion, we next sought to reinforce actin polymerization. At the leading edge of migrating cells, two major actin nucleators, namely Arp2/3 and formin, coordinate the forward extension of the plasma membrane (*77–79*). While Arp2/3 stays at the branch points of the actin filaments and does not engage in the incorporation of monomeric actin into the growing ends, formin continuously binds to the growing ends and actively recruits profilin-actin complexes to the elongating tips. Consequently, one subtype of formin, mDia1, accelerates actin polymerization 4–5-fold (*80, 81*) and drives motility 6–23 times faster than Arp2/3 in *in vitro* motility assays using functionalized beads (*30, 28, 31*). Furthermore, a theoretical study suggests that actin polymerization perpendicular to the membrane, characteristic of formin-driven filopodia formation, more efficiently transduces the polymerization force to advance the membrane (*82*). Therefore, we reasoned that the integration of mDia1 could enhance protrusive activity.

To test whether mDia1 promotes membrane protrusion, we first used living cells and performed a plasma membrane recruitment assay (Fig. S15A) (*83–85*). More specifically, we took FH1-FH2-DAD domains from mDia1 as a constitutively active form (*86*) and recruited it to the plasma membrane by the CID system. As shown in fig. S15B–H, YF-mDia1 translocation induced filopodia-like protrusions (fig. S15B–H). These protrusions were positive for Lifeact signal and showed coiling motions with their elongating tips labeled with YF-mDia1 (fig. S15D), consistent with formin-dependent filopodia elongation (*87–90*). Together, these data suggest that membrane recruitment of mDia1 generates an actin-dependent protrusive force on the membrane.

### mDia1-driven actin polymerization drives forward movement of GUVs

We then constructed the light-inducible version of mDia1 (mCherry-SspB-mDia1) for *in vitro* reconstitution (Fig. S1). Pyrene actin assays suggested that mDia1 had an effect of increasing the initial rate of actin polymerization (Fig. S8E, F). We encapsulated mCherry-SspB-mDia1 with iLID-YFP-SNAP, actin (10% Alexa-647 labeled), profilin, and cofilin in GUVs. Upon local recruitment of mDia1, some vesicles showed gradual drift toward the stimulated direction, suggesting that mDia1-generated actin polymerization can produce protrusive forces in the reconstituted lipid vesicles (fig. S16A, B). However, the response was modest and difficult to detect.

Because mDia1 alone produced only limited protrusive activity, we added pVCA-SspB, Arp2/3, and capping protein to better mimic actin network organization of migrating cells (Fig. 4A). Strikingly, this full component system (pVCA + mDia1) produced robust directional movement toward light stimulation in 18 out of 22 GUVs (Fig. 4B–D, fig. S16C, Movie S7, S9, S10). Polarized actin accumulation at the illuminated side preceded membrane displacement, consistent with actin polymerization driving forward protrusion (Fig. 4B, Movie S7). The vesicle continued to move steadily for an hour until the light was turned off (Fig. 4B, D, Movie S7). Furthermore, when we shifted the position of the blue light, the signals for mDia1, pVCA, and actin reoriented, causing the vesicle to move in the new direction, resembling chemotactic reorientation in migrating neutrophils (Fig. 4B, D, Movie S7). The speed of pVCA/mDia1 dual system was approximately threefold higher than that of mDia1 GUVs (Fig. 4G, H, S16B–C). The distance the front side membrane moved in 30 minutes was also significantly longer in pVCA + mDia1 GUVs (Fig. 4I). Furthermore, membrane recruitment of mDia1 and pVCA occasionally showed local protrusions, suggesting the generation of protrusive force (fig. S17, Movie S8). Together, these results indicate that mDia1-driven actin polymerization generates protrusive force that is substantially enhanced by cooperative activity with pVCA and Arp2/3.

**Fig. 4.**
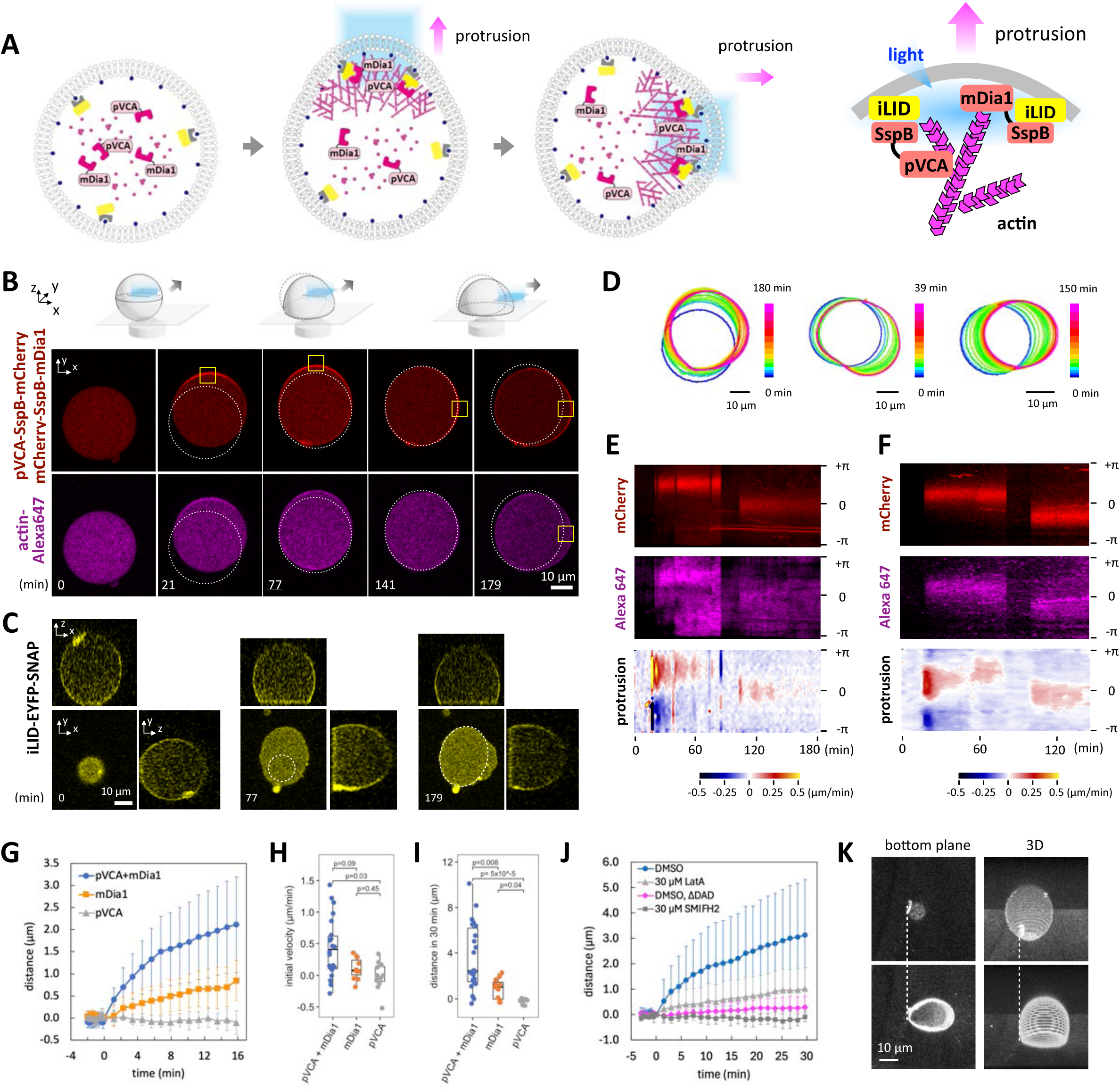
mDia1-mediated actin polymerization drives protrusive movement of GUVs. (A) Schematic representation of light-inducible membrane protrusion driven by pVCA and mDia1-mediated actin polymerization. (B) Representative images of pVCA-mDia1-mediated movement of GUVs. Yellow boxes indicate the area of blue light stimulation. Dotted line shows the initial position of the GUV. (C) Representative images of iLID-EYFP-SNAP showing the increase in adhesive area and vesicle deformation into dome-like shape. X-y images show the confocal plane right above the bottom substrate. Z-x and y-z images are reconstituted from z-stack. Dotted line shows the adhesive area of the previous images. (D) Color-coded boundaries of GUVs. The data correspond as follows. Left: Fig. 4B. Middle: fig. S16A. Right: fig. S16B. (E, F) Kymographs of membrane signal of mCherry and actin Alexa-647 and membrane protrusions. Triangles in (E) indicate the timepoints represented in (B, C). Data of (E) and (F) correspond to Fig. 4 (B) and fig. S16A, respectively. (G) Distance the front (illuminated) side of GUV membrane moved forward. pVCA+mDia1 (1 µM GST-pVCA-SspB-mCherry + 1 µM mCherry-SspB-mDia1, n=14 vesicles), mDia1 (1 µM mCherry-SspB-mDia1, n=12 vesicles), pVCA (1 µM GST-pVCA-SspB-mCherry, n=12 vesicles). Error bars indicate 95% CI (H, I) Quantification of the initial velocity and the distance the front side of GUV membrane moved forward. P values of Steel-Dwass test (two sided) are indicated. pVCA + mDia1: n=22 vesicles. mDia1: n=11 vesicles. pVCA: n=12 vesicles. Box whisker plots represent median, 1st, 3rd quartiles and 1.5×inter-quartile range. (J) Distance the front (illuminated) side of GUV membrane moved forward. DMSO (n=9 vesicles), 30 µM LatA (Latrunculin A) (n=10 vesicles). DMSO, ΔDAD: mCherry-SspB-mDia1 (FH1-FH2-DAD) was replaced with mCherry-SspB-mDia1(FH1-FH2) (n=13 vesicles). 30 µM SMIFH2 (n=6 vesicles). Error bars indicate 95% CI. (H, I) Quantification of the initial velocity and the distance the front side of GUV membrane moved forward. P values of Steel-Dwass test (two sided) are indicated. pVCA + mDia1: n=22 vesicles. mDia1: n=11 vesicles. pVCA: n=12 vesicles. Box whisker plots represent median, 1st, 3rd quartiles and 1.5×inter-quartile range. (K) A GUV deformed into dome-like shape after movement. Upper images: before light stimulation. Lower images: after light stimulation. The data corresponds to fig. S16C.

When mDia1 and pVCA were replaced with an equivalent amount of mCherry-SspB, vesicles did not move even with the local blue light illumination (fig. S18A). In addition, the kymograph analysis revealed that the timing and direction of membrane protrusions correlated with actin intensity and its polarity, consistent with actin-driven dynamics (Fig. 4E, F). Notably, the kymographs also revealed that protrusions at the front often coincide with retractions at the rear. Corroborating this, the actin sequestering compound Latrunculin A and the formin inhibitor SMIFH2 substantially compromised the motion of vesicles (Fig. 4J). Furthermore, deletion of the DAD domain from mDia1, which is responsible for efficient actin nucleation by mDia1 (*91*), abolished this motion (Fig. 4J). Together, these results indicate that membrane protrusion and motion of the vesicles are driven by actin polymerization.

### The pVCA–mDia1 combination forms a fast-turnover, actively polymerizing actin network

To elucidate the mechanistic basis of the distinct actin architectures generated by pVCA, mDia1, and the pVCA–mDia1 dual system and their contributions to vesicle motion, we combined ultrastructural, pharmacological, and FRAP analyses of reconstituted actin networks.

To examine ultrastructure, we performed EM on actin assemblies formed on SsrA-coated SUVs. Although assemblies deposited onto a solid support may not fully represent filament organization within GUVs, micrometer-scale filaments were observed under all conditions, with the highest filament density observed when pVCA was used as the sole NPF (fig. S12D, E). However, filament density alone did not explain the observed differences in vesicle motion, underscoring the importance of actin network dynamics.

To probe network dynamics, we examined the pharmacological responses of actin assemblies in a system where NPFs were constitutively tethered to the GUV membrane via SsrA. Consistent with EM observations, pVCA generated the highest basal level of membrane-associated actin but showed only modest additional F-actin accumulation upon Jasplakinolide treatment (fig. S10B–F). In contrast, mDia1-driven assemblies exhibited a progressive increase in membrane-associated actin in response to Jasplakinolide, with the most pronounced effect observed in the combined pVCA-mDia1 system (fig. S10E, F), consistent with the enhanced polymerization capacity of the dual condition observed in bulk pyrene–actin assays (fig. S8F). Notably, Jasplakinolide treatment of mDia1 and pVCA-mDia1 GUVs was occasionally accompanied by outward membrane deformation, consistent with polymerization-generated forces acting on the membrane (fig. S10G). Moreover, the pVCA-mDia1 dual system was least susceptible to Latrunculin A, indicating a shift in network behavior toward sustained polymerization (fig. S10E, F).

FRAP analysis further distinguished network dynamics. pVCA-driven actin networks displayed rapid fluorescence recovery, suggesting highly dynamic actin turnover (fig. S11C, E). In contrast, actin assemblies generated by mDia1 showed minimal fluorescence recovery of actin and mDia1 itself (fig. S11B–E). This slow recovery is consistent with processive filament elongation at the membrane, where mDia1 remains persistently associated with growing barbed ends. Importantly, the pVCA–mDia1 dual system—despite exhibiting strong Jasplakinolide-induced actin accumulation—retained rapid fluorescence recovery in FRAP (fig. S11C, E), indicating that the dual network maintains dynamic turnover while sustaining high polymerization activity.

Together, these observations indicate that the pVCA–mDia1 system exhibits high polymerization capacity and rapid turnover, properties potentially well suited for efficient force generation and directional vesicle movement.

### Deformation and adhesion of GUVs

Quantitative analysis revealed that the initial velocity of each GUV was strongly correlated with its net displacement after 30 minutes (*R* = 0.89; fig. S19A). In addition, displacement showed a moderate positive correlation with vesicle diameter, suggesting that larger vesicles can accommodate greater chemical and cytoskeletal resources and may also be more favorable for generating light-induced asymmetry. In contrast, vesicles showing membrane localization of pVCA and mDia1 prior to blue light illumination tended to exhibit smaller displacement. This suggests that nonspecific membrane binding limits the light-dependent establishment of asymmetry (fig. S19B, C).

We next analyzed changes in velocity and vesicle shape. The vesicles initially moved at an average velocity of 0.43 µm/min, but their speed gradually decreased over time (Fig. 4G, H, fig. S16C). Concomitant with this deceleration, the contact area of the bottom side of the vesicles progressively increased during the movement, stabilizing the vesicles in a dome-like shape (Fig. 4C, K). This increase in the contact area was not seen in the GUVs when pVCA and mDia1 were replaced with mCherry-SspB (fig. S18B). Thus, we speculate that the contact to the bottom substrate was induced by the protrusive reaction and that the adhesion and the concomitant deformation led to cumulative tension increase (*92*) which in turn counteracted the protrusive force. The force exerted by actin polymerization may not be enough to detach the GUV from the BSA-coated glass substrate. This is consistent with the observation that an initial contact site remained adherent even in a GUV that traveled farthest (Fig. 4K, fig. S20B, C, Movie S9, S10). Additionally, actin recycling in the system may be insufficient and the force gradually diminishes over time (*71*). Despite these potential resisting factors, the combination of mDia1 and pVCA induced movement consistently (Fig. 4, fig. S16C, S20). Together, these observations support the conclusion that asymmetric actin polymerization mediated by pVCA–mDia1 is sufficient to drive forward vesicle movement under the tested conditions.

### Computational modeling of actin network dynamics underlying membrane deformation

To dissect the mechanisms underlying actin-driven membrane deformation, we constructed a computational model of actin network dynamics in a minimal vesicle system. We hypothesized that membrane deformation could arise from intrinsic mechanical bracing within the actin network. Accordingly, we considered two candidate mechanisms: Arp2/3-dependent network organization and frictional interactions between actin filaments (*93, 94*). The model was implemented using the actin-network simulation framework Cytosim (*95*) and coupled to a deformable membrane (*96, 97*) (Supplementary Text 4 and Methods).

The simulations captured distinct NPF-dependent behaviors (fig. S21A, B, Movie S11, S12). pVCA produced short, branched filaments that frequently detached from the membrane, whereas mDia1 drove rapid filament elongation while remaining membrane-associated. Without inter-filament steric interactions, the pVCA + mDia1 condition yielded marginal but reproducible forward membrane motion (fig. S21A, C, Movie S11), indicating cooperativity between mDia1 and pVCA. Incorporating inter-filament interactions resulted in robust membrane deformation (fig. S21B, D, Movie S12), with extension increasing in the order pVCA < mDia1 < pVCA+mDia1, consistent with experiments (Fig. 4G–I). These results support a minimal internal bracing mechanism in which membrane-anchored polymerization, Arp2/3-dependent branching, and inter-filament interactions collectively generate protrusive force.

The simulations also revealed substantial retrograde flow of actin filaments (fig. S21A, B, Movie S11, S12). This provides a mechanistic explanation for why the observed vesicle motility (0.1–1.0 µm/min, Fig. 4G–J, fig. S16) is more than an order of magnitude slower than the actin polymerization rate (1–10 µm/min) (*80, 81*) and neutrophil migration (∼10 µm/min) (*98*). Incorporating additional backing mechanisms that provide mechanical resistance to the actin network, such as external anchoring or confinement (*21, 22*), is therefore predicted to enhance protrusive efficiency and bridge minimal synthetic systems toward cellular modes of amoeboid migration.

## Discussion

In the present study, we developed a fully reconstituted system to achieve light-guided directional movement in a protocell. We leveraged mDia1-mediated actin polymerization to establish the forces that drive movement. Our findings not only corroborate recent cell biological findings on formin’s significant roles in cell migration (*87, 98–103*) but also show that the mDia1-mediated actin polymerization can drive the forward movement of lipid vesicles in a minimal, reconstituted system, overcoming challenges thus far posed by genetic redundancy and cellular complexity. Formin-mediated actin polymerization exhibits several unique features compared to free barbed end elongation in Arp2/3-dependent polymerization, including processive elongation (*31, 80*), accelerated polymerization rate (*80, 81*), rotation, and coiling motion (*89, 90, 104*). These characteristics could contribute to the effective propulsion of the vesicle. Remarkably, our minimal system further indicated a functional interplay between pVCA and mDia1. Although further investigation is required, this cooperative effect likely arises from coordination between the two nucleators: mDia1 generates mother filaments required for Arp2/3 nucleation, and, in turn, Arp2/3-mediated branching increases the number of growing ends available for mDia1 elongation (*100, 101*). To further elucidate the mechanical coupling to force generation and membrane protrusion, unveiling the nanoscopic architecture of the actin network in the migrating protocell will be crucial. Our protocell approach, which bridges the gap between cellular and biochemical domains, opens new avenues for comprehensively interrogating modules of cellular motility.

We achieved a GUV displacement speed in the range of 0.1–1.0 µm/min, which is comparable to the migration rates of typical fibroblasts (*105*) and other adherent cells including those that competed in the World Cell Race (*106*). How could we make the movement even faster? *L. monocytogenes* and NPF-coated beads in purified cytoskeletal protein solution typically move in the range of 1–15 µm/min (*27, 28, 30–33*). Some adherent cells can migrate at 5 µm/min even when actomyosin contractility is inhibited (*19*). We propose three possible ways to make artificial cell migration faster. First, increasing actin concentration (*81, 107*) and integrating additional factors, such as VASP (*108*), Fascin (*109*), and NPFs (*67, 110*), could accelerate the rate of actin polymerization. Secondly, optimizing membrane tension could enhance the deformability of the lipid membrane, akin to the malleable structures observed in motile cells, thereby potentially promoting more efficient protrusion and motility (*50, 111*). Finally, fine-tuning adhesion could enhance migration efficiency as many cells exhibit a preference for an optimal adhesion range, where both insufficient and excessive adhesion can hinder motility (*112, 113*).

Interestingly, the phenotypes of mDia1 membrane recruitment differed between cells and GUVs. In cells, mDia1 produced filopodia-like thin protrusions (fig. S12), whereas in GUVs, it pushed the lipid membrane as a continuous plane structure on a much broader scale than filopodia (Fig. 4, fig. S13–16). Since living cells tend to have higher membrane tension than liposomes (*114*), it may appear counterintuitive that the cell membrane was more susceptible to mDia1-induced local deformation. One possible explanation is that, in cells, endogenous proteins might work in concert with mDia1 to amplify local membrane deformation and form long filopodia-like protrusions (*100, 115*). Alternatively, localized defects in membrane tension could predispose certain areas to form thin protrusions (*116*). Additionally, cytoskeleton-bound transmembrane proteins impede tension propagation in cells (*117*). Furthermore, the nuclear accumulation of mDia1 in cells (fig. S12) could reduce its effective concentration and cause a heterogeneous distribution on the cell membrane, resulting in focused protrusive force in regions with high mDia1 concentration, similar to how pVCA-driven finger-like protrusions are formed in liposomes, as demonstrated by Gat *et al.* (*118*). Since our light-inducible actuators are genetically encodable, the same molecular toolkit can be deployed in both GUVs and living cells—providing a unified platform to dissect cellular morphodynamics.

We have recently shown that chemically-induced membrane recruitment of ActA, an NPF derived from *L. monocytogenes*, and subsequent global actin polymerization lead to eccentricity in the shape of GUVs (*56*). We noted a difference in membrane deformation between the CID and LID systems, as the CID could induce a clear membrane deformation given a global, non-directional input. This difference could be attributed to the variations in cytoskeletal proteins and input signals between the two systems. In the present study, we have incorporated profilin, cofilin, and capping proteins. Since these proteins accelerate actin depolymerization (fig. S8B), they may render the actin network more plastic rather than exerting force to deform the membrane. Consistent with this, stabilization of F-actin by Jasplakinolide occasionally resulted in outward membrane deformation in GUVs containing mDia1 or the pVCA–mDia1 combination (fig. S10G). Moreover, the binding kinetics of iLID-SspB (k_on_ =1.2 × 10^3^ M^-1^s^-1^, k_off_ =1.1 × 10^-3^ s^-1^, K_d_ = 0.8 µM) substantially differ from those of FKBP-FRB (k_on_ =1.9 × 10^6^ M^-1^s^-1^, k_off_ =2.2 × 10^-2^ s^-1^, K_d_ = 12 nM) (*60, 119, 120*). This may result in different reaction dynamics between actin cytoskeleton and the lipid membrane.

Our finding that internal cytoskeletal force can drive the unidirectional motion of lipid vesicles offers avenues to reverse-engineer cell motility. Although we used light as an input, asymmetric actin polymerization may be achievable with chemical inputs with consideration of binding affinity (*83, 121*), transmembrane signal transduction mechanisms (*122*), and self-organized pattern formation (*123–127*). For sustained motility, it will be important to incorporate other migratory forces—reconstructing myosin contractility at the rear (*128*) and regulating adhesion/de-adhesion turnover (*119, 129*) in concert with actin polymerization. Furthermore, integrating synthetically engineered molecules (*81, 130, 131*) could enhance—motility even beyond the limits of natural systems. Achieving synthetic motility in GUVs could be a step toward future developments in active therapeutic agents, self-organizing artificial tissues, and synthetic neural circuits.

## Materials and Methods

### Reagents

ATP was purchased from Gold Biology (A-081) or Nacalai Tesque (01072-82), prepared as 100 mM solution in 100 mM Tris-HCl (pH 7.4 at RT), and stored at -20°C. Creatine phosphate was purchased from Gold Biotechnology (C-323-5) or Oriental Yeast (45180000), prepared as 1 M solution in PBS (Gibco 10010-023), stored at -20°C. Creatine kinase was purchased from Sigma-Aldrich (10127566001) or Oriental Yeast (46430003), prepared as 40 µM solution in buffer V (50 mM HEPES-NaOH (pH 7.5), 150 mM NaCl, 7 mM β-mercaptoethanol, 10% glycerol), snap-frozen, and stored at -20°C. Flavin mononucleotide (FMN) was purchased from TCI (R0023) and Wako (06500171), freshly prepared as 600 µM solution in Milli-Q water before iLID expression. Alexa Fluor 647 C2 Maleimide (A20347) and Pierce™ Glutathione Agarose (16100) were purchased from Thermo Fisher Scientific. Hexadecane (H6703), Silicone oil (378348), and SMIFH2 (S4826) were purchased from Sigma-Aldrich. Latrunculin A was purchased from Wako (125-04363). Jasplakinolide was purchased from Cayman Chemical Company (11705). Benzylguanine-PEG2000-DSPE was prepared and purified as previously described (*61*). POPC (1-palmitoyl-2-oleoyl-glycero-3-phosphocholine, 850457), POPS (1-palmitoyl-2-oleoyl-sn-glycero-3-phospho-L-serine, 840034), 18:1 DGS-NTA-Ni (1,2-dioleoyl-sn-glycero-3-[(N-(5-amino-1-carboxypentyl)iminodiacetic acid)succinyl] (nickel salt), 790404), 18:1 PE-PEG2000-benzylguanine (1,2-dioleoyl-sn-glycero-3-phosphoethanolamine-N-[benzylguanine(polyethylene glycol)-2000], 880137), and 18:1 Biotinyl Cap PE (1,2-dioleoyl-sn-glycero-3-phosphoethanolamine-N-(cap biotinyl) (sodium salt), 870273C) were purchased from Avanti Polar Lipids as chloroform solution. Actin (AKL99), pyrene labeled actin (AP05), Arp2/3 (RP01P), Profilin (PR02), Cofilin (CF01), were purchased from Cytoskeleton. Strep-Tactin XT (2-4010-010, discontinued) resin, Strep-Tactin XT 4Flow (2-5010-010) resin, and purified Strep-Tactin protein were purchased from IBA. Ni-NTA agarose (30210) was purchased from Qiagen. Amylose resin (E8021S) was purchased from New England BioLabs.

### Plasmid construction

#### pQE80L-6×His-MBP-TEVprorease(S219V)

Ser219Val mutation (*132*) was introduced by PCR with the template plasmid gifted from Dr. Yubin Zhou and the primer set of fwd: 5’-TGGTGAAACCTGAAGAACCTTTT-3’ and rev: 5’-TTCACCATGAAAACTTTATGGCC-3’. TEV prorease (S219V) was then PCR amplified with the primer set of fwd: 5’-AATTGAGCTCGATGAGCGGCCTGGTGC-3’ and rev: 5’-AATTGGATCCTTATTGCGAGTACACCAATTCATTCATG-3’ and inserted between SalI and BamHI sites of pQE-80L MBP-SspB Nano plasmid (Addgene #60409) by using restriction digestion and T4 ligase ligation process.

#### pQE80L-6×His-MBP-TEVsite-mCherry-SspB(micro)

EYFP-2×SAGG linker was PCR amplified from the template plasmid YFP-FKBP (*133*)with the primer set fwd: 5’-GTATTTTCAGGGATCGCCGCTAGCGCTACCGG-3’ and rev: 5’-TCGGGGAGCTGGATCCTCCGCCAGCGCTGC-3’ and inserted into BamHI site of pQE-80L MBP-SspB Nano plasmid (Addgene #60409) by using Gibson assembly. EYFP part was then replaced with mCherry from pmCherry-C1 (Clontech) by restriction digestion and T4 ligase ligation process with AgeI and BsrGI. Arg73 of SspB was mutated to Gln by PCR with the primer set fwd: 5’-AACGCCCagTTTAAGGGCGTGTCTCGT-3’ and rev: 5’-CTTAAActGGGCGTTGAACTGGATAA-3’.

#### pGEX-2T-GST-3Csite-TEVsite-6×His

Two sets of oligo DNA pairs were designed to introduce 3C protease cleavage site, TEV protease cleavage site, and 6×His tag (1st set, fwd: 5’-<u>GGTTCCGCGTGGATC</u>TGGTCTTGAGGTGCTCTTTCAGGGACCCGGCAGTCTCGAGGGTCTG<u>TACAAGCG</u> <u>AATTCAG</u>-3’, rev: 5’-<u>CTGAATTCGCTTGTA</u>CAGACCCTCGAGACTGCCGGGTCCCTGAAAGAGCACCTCAAGACCA<u>GATCCACG CGGAACC</u>-3’. 2nd set, fwd: 5’-<u>TACAAGCGAATTCAG</u>GAGAACCTCTACTTTCAAAGCGATCATCATCATCATCATCACTAAA<u>AATTCATCGT GACTG</u>-3’ and rev: 5’-<u>CAGTCACGATGAATT</u>TTTAGTGATGATGATGATGATGATCGCTTTGAAAGTAGAGGTTCTC<u>CTGAATTCG CTTGTA</u>-3’ (underlines indicate overhang sequences)). To anneal oligo DNAs individually, 0.5 µM each DNAs were mixed in 1× T4 Ligase buffer (NEB, B0202S) and heated at 85°C for 10 min and cooled 5°C per 2 min down to 40°C. pGEX-2T, which was kindly gifted from Dr. Miho Iijima, was digested by BamHI and EcoRI and assembled with the two annealed DNA fragments by Gibson assembly.

#### pGEX2T-GST-pVCA-3Csite-SspB(micro)-mCherry-TEVsite-6×His

To first construct pGEX2T-GST-pVCA-3Csite-TEVsite-6×His, pVCA (proline rich region and VCA domain) of human N-WASP was PCR amplified with the primer set fwd: 5’-CAGTCTCGAGGGAGGTGTTGAAGCTGTTAAAAATGA-3’ and rev: 5’-TCCTGAATTCCGTCTTCCCACTCATCATCATCCTC-3’ and inserted between XhoI and EcoRI site of pGEX-2T-GST-3Csite-TEVsite-6×His by using restriction digestion and T4 ligase ligation. Then, SspB(micro)-mCherry was PCR amplified with the primer set fwd: 5’-TGAGTGGGAAGACGGAATTGGGCAGTCGACGGTACCGCG-3’ and rev: 5’-GTAGAGGTTCTCCTGAATTCCCTTGTACAGCTCGTCCATGCC-3’ and inserted in the EcoRI site of pGEX2T-GST-pVCA-3Csite-TEVsite-6×His by Gibson assembly.

#### pET28-2×Strep-iLID-EYFP-MARCKS

First, pET28-2×Strep-iLID-6×His was constructed as follows. iLID sequence was cut out from the pLL7.0: Venus-iLID-Mito (Addgene #60413) by NheI and EcoRI and inserted between NheI and EcoRI sites of pET28-2×Strep-tag gifted from Dr. Kanemaki by using restriction digestion and T4 ligase ligation. Then, to construct pET28-2×Strep-iLID-EYFP-6×His, EYFP was PCR amplified from pEYFP-C1 (Clontech) with the primer set fwd: 5’-AATTGTCGACATGGTGAGCAAGGGCGAG-3’ and rev: 5’-AATTGCGGCCGCCTTGTACAGCTCGTCCATGC-3’ and inserted between SalI and NotI sites of pET28-2×Strep-iLID-6×His by restriction digestion and T4 ligation. MARCKS-ED fragment was constructed by oligo annealing (1st set fwd: 5’-GTACAAGGGAAGTGCTGGTGGTAAAAAGAAAAAGAAGCGCTTTTCCTTC-3’ and rev: 5’-TTCTTGAAGGAAAAGCGCTTCTTTTTCTTTTTACCACCAGCACTTCCCTT-3’, 2nd set fwd: 5’-AAGAAGTCTTTCAAGCTGAGCGGCTTCTCCTTCAAGAAGAACAAGAAGTA-3’ and rev: 5’-GTACTACTTCTTGTTCTTCTTGAAGGAGAAGCCGCTCAGCTTGAAAGAC-3’) and inserted into BsrGI site of pET28-2×Strep-iLID-EYFP-6×His.

#### pET28-2×Strep-iLID-EYFP-SNAP

SNAP tag sequence was PCR amplified from the template Phage-ubc-nls-ha-tdMCP-SNAP, a kind gift from Dr. Bin Wu, with the primer set fwd: 5’-AATTTGTACAAGTCTGCTGGCGGAAGCGCTGGAGGCAGCATGGACAAAGACTGCGAAATGAAGC-3’ and rev: 5’-AATTGCGGCCGCTTAACCCAGCCCAGGCTTG-3’ and inserted between BsrGI and NotI of pET28-2×Strep-iLID-EYFP-MARCKS by restriction digestion and T4 ligation.

#### pET28-2×Strep-SsrA-SNAP

The plasmid was constructed by replacing the iLID-EYFP part of pET28-2×Strep-iLID-EYFP-SNAP with the sequence containing SsrA peptide (AANDENY). A set of oligo DNA (5’-AGCGGGTGCCGCTAGC<u>GCAGCGAACGACGAAAATTAC</u>GGACCGGTCGGTCTGTACAAGTCTGCTGG-3’ and 5’-CCAGCAGACTTGTACAGACCGACCGGTCC<u>GTAATTTTCGTCGTTCGCTGC</u>GCTAGCGGCACCCGCT-3’, (underlines indicate SsrA peptide region)) was annealed and inserted between NheI and BsrGI site of pET28-2×Strep-iLID-EYFP-SNAP by Gibson assembly.

#### mCherry-SspB(micro) (Clontech C1)

SspB(micro) sequence was PCR amplified from pQE80L-6×His-MBP-TEVsite-mCherry-SspB(micro) with the primer set fwd: 5’-GAACAGTACGAACGCGCC-3’ and rev: 5’-AATTCTCGAGGACCACCAGCACTACCACCAGCACTACCACCAGCACTACCACCAGCACTACCACCAGCACTACCAATATTCAGCTCGTCATAGATT-3’ and inserted between BsrGI and XhoI sites by restriction digestion and T4 ligation. A silent mutation was introduced to BamHI site upstream of SspB by inverse PCR with the primer set fwd: 5’-GAGGATCtAGCTCCCCGAAACGCCCT-3’ and rev: 5’-GGGAGCTaGATCCTCCGCCAGCGCTG-3’

#### pQE80L-MBP-TEVsite-mCherry-MCS

mCherry sequence was PCR amplified from pmCherry-C1 (Clontech) with the primer set fwd: 5’-GTATTTTCAGGGATCGCTAGCGCTACCGGTC-3’ and rev: 5’-TCAGCTAATTAAGCTATCAGTTATCTAGATCCGGTGGATC-3’ and inserted between BamHI and HindIII sites of pQE-80L MBP-SspB Nano by Gibson assembly.

#### 2×Strep-mCherry-MCS

mCherry sequence was PCR amplified from pmCherry-C1 (Clontech) by the primer set fwd: 5’-AGCGGGTGCCGCTAGTCATATGGGTACGCTAGCGCTACCGGTCG-3’ and rev: 5’-GGTGGTGGTGCTCGATCAGTTATCTAGATCCGGTGGATCC-3’ and inserted between NheI and XhoI sites of pET28-2×Strep-tag by Gibson assembly.

#### 2×Strep-MBP-TEVsite-mCherry-MCS

MBP-TEVsite-mCherry-MCS was PCR amplified from the template pQE80L-MBP-TEVsite-mCherry-MCS with the primer set fwd 5’-CAAATGGGTCGGATCGACGGATCTAAAATCGAAGAAGG-3’ and rev: 5’-GGTGGTGGTGCTCGATCAGTTATCTAGATCCGGTGGA-3’ and inserted between BamHI and XhoI sites of pET28-2×Strep-tag by Gibson assembly.

#### pCold-6×His-2×Strep-MBP-TEVsite-mCherry-SspB(micro)-mDia1(FH1-FH2-DAD)

A DNA fragment 2×Strep-MBP-TEVsite-mCherry was PCR amplified from 2×Strep-MBP-mCherry-MCS with the primer set fwd: 5’-TCGAAGGTAGGCATATGGGCTGGTCTCACCC-3’ and rev: 5’-ACAGCTCGTCCATGCCG-3’. The other DNA fragment SspB(micro) was PCR amplified from mCherry-SspB(micro) (Clontech C1) with the primer sets fwd: 5’-GCATGGACGAGCTGTACAAG-3’ and rev: 5’-TCATTCTTGGCCATAGCTTGAGCTCGAGGACC-3’. pCold-mDia1, which was a kind gift from Dr. Roberto Dominguez, was digested by NdeI and assembled with the two DNA fragments by Gibson assembly.

#### pET28-2×Strep-mCherry-SspB(micro)-mDia1(FH1-FH2-DAD)

mCherry-SspB(micro)-mDia1(FH1-FH2-DAD) was cut out from pCold-6×His-2×Strep-MBP-TEVsite-mCherry-SspB(micro)-mDia1(FH1-FH2-DAD) by AgeI and BamHI and inserted between AgeI and BamHI sites of 2×Strep-mCherry-MCS by restriction digestion and T4 ligation.

#### pET28-2×Strep-mCherry-SspB(micro)-TEVsite-6×His

A DNA fragment mCherry-SspB(micro) was PCR amplified from mCherry-SspB(micro) (Clontech C1) with the primer set fwd: 5’-AGCGGGTGCCGCTAGCGGACCGGTCGCCACCATG-3’ and rev: 5’-AAAGTAGAGGTTCTCAGATCCGGTGGATCCCGG-3’. The other fragment containing TEV cleavage site was made by oligo annealing of a set of DNA 5’-GAGAACCTCTACTTTCAAagcgatgTCGAGCACCACCACC-3’ and 5’-GGTGGTGGTGCTCGAcatcgctTTGAAAGTAGAGGTTCTC-3’. These fragments were inserted between NheI and sites of pET100 by Gibson assembly.

#### pET28-2×Strep-mCherry-SspB(micro)-mDia1(FH1-FH2-DAD)-TEVsite-6×His

A DNA fragment mDia1(FH1-FH2-DAD) was PCR amplified from pCold-mDia1 with the primer set fwd: 5’-GGTCCTCGAGCCAAGAATGAAATGGCTTCTCTCTC-3’ and rev: 5’-CGGTGGATCCGCTTGCACGGCCAACCAG-3’ and inserted between XhoI and BamHI sites of pET28-2×Strep-mCherry-SspB(micro)-TEVsite-6×His by Gibson assembly.

#### pET28-2×Strep-mCherry-SspB(micro)-mDia1ΔDAD(FH1-FH2)-TEVsite-6×His

A DNA fragment mDia1(FH1-FH2) was PCR amplified from pCold-mDia1 with the primer set fwd: 5’-GGTCCTCGAGCCAAGAATGAAATGGCTTCTCTCTC-3’ and rev: 5’-CGGTGGATCCCTTCTCCTTGGCTAATTTTGCTCTCCG-3’ and inserted between XhoI and BamHI sites of pET28-2×Strep-mCherry-SspB(micro)-TEVsite-6×His by Gibson assembly.

#### EYFP-FKBP-mDia1(FH1-FH2-DAD)

mDia1(FH1-FH2-DAD) was cut out from pCold-6×His-2×Strep-MBP-TEVsite-mCherry-SspB(micro)-mDia1(FH1-FH2-DAD) by XhoI and BamHI and inserted between XhoI and BamHI of EYFP-FKBP by restriction digestion and T4 ligation.

#### pET28-6×His-2×Strep-TEVsite-Capβ2(mouse)-Capα1(mouse)

First, pET28-6×His-2×Strep was constructed as follows. 6×His sequence was synthesized by oligo annealing of fwd: 5’-AGGAGATATACCATGGGCCATCACCATCACCATCACATGGGCTGGTCTCA-3’ and rev: 5’-TGAGACCAGCCCATGTGATGGTGATGGTGATGGCCCATGGTATATCTCCT-3’ and inserted into NcoI site of pET28-2×Strep-tag. Capβ2(mouse)-Capα1(mouse) was PCR amplified from the template of pET3d-Capβ2(mouse)-Capα1(mouse), gifted from Dr. Julie Plastino and Dr. Cecile Sykes, with the primer set fwd: 5’-CTGGAGCGGGTGCCGCTAGCGAGAACCTCTACTTTCAAAGCGATATGAGCGATCAGCAGCTGG-3’ and rev: 5’-TGTCGACGGAGCTCGAATTCTTAAGCATTCTGCATTTCTTTGCCAATC-3’ and inserted between NheI and EcoRI of pET28-6×His-2×Strep by Gibson assembly. TEV cleavage site was designed in the forward primer and introduced between 2×Strep tag and Capβ2 ORF.

### Actin preparation

Actin for GUV experiments was reconstituted from lyophilized powder as described in manufacturer’s instructions, aliquoted, snap-frozen in liquid nitrogen, and stored at -80°C. For actin polymerization assays, actin was dissolved in G-buffer [2 mM Tris-HCl (pH 7.5 at RT), 0.1 mM CaCl_2_, 0.2 mM ATP, 0.5 mM DTT, 1 mM NaN_3_] at concentration less than 65 µM and dialyzed against G-buffer for 3 days with daily buffer exchange. Dialyzed actin was then purified by a Superdex 200 increase 10/300 GL column (Cytiva) to separate monomer fraction from occasional larger molecular size (smaller elution volume) fraction. The purified actin was stored at 4°C with dialysis against G-buffer. The dialysis buffer for actin storage was exchanged twice a week. Alexa 647 labeling of actin was performed following the previously reported method (*134*). First, rabbit skeletal muscle actin (Cytoskeleton Inc., #AKL99) was dissolved in G*-buffer [2 mM Tris-HCl (pH 7.5 at RT), 0.1 mM CaCl_2_, 0.2 mM ATP] at concentration less than 65 µM. The solution was dialyzed against G*-buffer overnight, followed by an additional 3 hour dialysis with fresh buffer. Dialyzed actin was collected and mixed with 4 molar excess Alexa Fluor 647 C2 maleimide (ThermoFisher, A20347, solubilized to 10 mM in DMSO) and incubated at 4°C overnight with rotation (10 min labeling as in the original protocol did not yield efficient labeling in the case of Alexa Fluor 647). The reaction was quenched by adding DTT to 10 mM. Aggregated proteins and insoluble dyes were removed by ultracentrifugation at 350,000×g (90,000 rpm, TLA-100.1 rotor) for 12 min. The supernatant was collected and mixed 1/10 volume of 10×KMEI buffer (500 mM KCl, 10 mM MgCl_2_, 10 mM EGTA, 100 mM Imidazole, pH 7.0) and final 1 mM ATP to polymerize actin at RT for 2–3 hours. Labeled filamentous actin was collected by ultracentrifugation at 195,000×g (67,000 rpm, TLA-100.1 rotor) for 30 min and resuspended with G-buffer. The labeled actin was dialyzed against G-buffer for 3 days with daily buffer exchange and purified through size exclusion chromatography with a Superdex 200 increase 10/300. The protein was snap-frozen in liquid nitrogen and stored at -80°C.

### Protein purification

Protein expression were carried out in BL21-CodonPlus (DE3)-RIL (Agilent Technologies) and cell lysis was performed by microfluidizer (Microfluidics, Model M-110Y), unless otherwise specified.

#### 6×His-MBP-TEV protease (S219V)

The bacteria transformed with pQE80L-6×His-MBP-TEVprotease (S219V) were cultured overnight in LB broth supplemented with 100 µg/mL Ampicillin and 25 µg/mL Chloramphenicol (LB/Amp/Cam). Pre-culture was inoculated to LB/Amp/Cam and cultured at 37°C until OD_600_ reached around 0.4. Protein expression was induced by adding 300 μM IPTG and allowed to proceed for 17 hours at 18°C. The cells were re-suspended with 50 mM Tris-HCl (pH 7.5 at RT), 250 mM NaCl, 5 mM MgCl_2_, 7 mM β-mercaptoethanol and lysed by microfluidizer. The lysate was centrifuged at 20,000×g at 4°C for 30 min to remove cell debris. The protein was purified by Ni-NTA, eluted around 60–100 mM imidazole, and then dialyzed against 25 mM Tris-HCl (pH 7.5 at RT), 200 mM NaCl, 7 mM β-mercaptoethanol. The elution was concentrated by Amicon Ultra (10 kDa cutoff) and further dialyzed against 25 mM Tris-HCl (pH 7.5 at RT), 200 mM NaCl, 7 mM β-mercaptoethanol, 50% glycerol. The protein was stored at -20 °C.

#### mCherry-SspB

The bacteria transformed with pQE80L-6×His-MBP-TEVsite-mCherry-SspB were cultured overnight in LB broth supplemented with 100 µg/mL Ampicillin and 25 µg/mL Chloramphenicol (LB/Amp/Cam). Pre-culture was inoculated to LB/Amp/Cam and cultured at 37°C until OD_600_ reached 0.4–0.6. Protein expression was induced by adding 300 μM IPTG and allowed to proceed for 18 hours at 22°C. The cells were re-suspended with 50 mM Tris-HCl (pH 7.5 at RT), 250 mM NaCl, 7 mM β-mercaptoethanol, 20 mM Imidazole supplemented with cOmplete EDTA-free protease Inhibitor Cocktail (MilliporeSigma) and lysed by microfluidizer. The lysate was centrifuged at 20,000×g at 4°C for 30 min to remove cell debris. The protein was first purified by Ni-NTA, eluted around 50 mM imidazole, and then dialyzed against 25 mM Tris-HCl (pH 7.5 at RT), 250 mM NaCl, 7 mM β-mercaptoethanol. 6×His-MBP tag was cleaved by 6×His-MBP-TEV protease (S219V) (purified in house) during the dialysis. The cleaved 6×His-MBP tag and TEV protease were removed through the second round of Ni-NTA column. Flow through and wash fractions were dialyzed against buffer S [25 mM Tris-HCl (pH 7.5 at RT), 250 mM NaCl, 7 mM β-mercaptoethanol] and concentrated by Amicon Ultra (10 kDa cutoff). The protein was snap-frozen in liquid nitrogen in storage buffer [25 mM Tris-HCl (pH 7.5 at RT), 250 mM NaCl, 7 mM β-mercaptoethanol, 20% glycerol] and stored at -80 °C.

#### iLID-YFP-MARCKS

The bacteria transformed with pET28-2×Strep-iLID-EYFP-MARCKS were cultured overnight in LB broth supplemented with 50 µg/mL Kanamycin and 25 µg/mL Chloramphenicol (LB/Kan/Cam). Pre-culture was inoculated to LB/Kan/Cam and cultured at 37°C until OD600 reached around 0.6. Protein expression was induced by adding 300 μM IPTG and allowed to proceed for 17 hours at 23°C in the presence of 6 µM FMN. The cells were resuspended with buffer L1 [50 mM Tris-HCl (pH 7.5 at RT), 200 mM NaCl, 1 mM EDTA, 7 mM β-mercaptoethanol] and lysed by microfluidizer. The lysate was centrifuged at 20,000×g at 4°C for 30 min to remove cell debris. The supernatant was applied to Strep-Tactin XT (IBA Lifesciences) equilibrated with buffer L1 and the protein was eluted with 50 mM biotin in buffer L1. The elution was dialyzed against buffer L2 [25 mM Tris-HCl (pH 7.5 at RT), 200 mM NaCl, 7 mM β-mercaptoethanol] and concentrated by Amicon Ultra (10 kDa cutoff). The protein was snap-frozen in liquid nitrogen in storage buffer [25 mM Tris-HCl (pH 7.5 at RT), 200 mM NaCl, 7 mM β-mercaptoethanol, 30% glycerol] and stored at -80 °C.

#### iLID-YFP-SNAP

The bacteria transformed with pET28-2×Strep-iLID-EYFP-SNAP were cultured overnight in LB broth supplemented with 50 µg/mL Kanamycin and 25 µg/mL Chloramphenicol (LB/Kan/Cam). Pre-culture was inoculated to LB/Kan/Cam and cultured at 37°C until OD_600_ reached around 0.5. Protein expression was induced by adding 300 μM IPTG and allowed to proceed for 20 hours at 23°C in the presence of 6 µM FMN. The cells were resuspended with buffer L1 (50 mM Tris-HCl (pH 7.5 at RT), 200 mM NaCl, 1 mM EDTA, 7 mM β-mercaptoethanol) and lysed by microfluidizer. The lysate was centrifuged at 20,000×g at 4°C for 30 min to remove cell debris. The supernatant was applied to Strep-Tactin XT (IBA Lifesciences) equilibrated with buffer L and the protein was eluted with 50 mM biotin in buffer L1. The elution was further purified by Superdex 200 increase 10/300 GL column in the buffer L2 [25 mM Tris-HCl (pH 7.5 at RT), 200 mM NaCl, 7 mM β-mercaptoethanol] and concentrated by Amicon Ultra (10 kDa cutoff). The protein was snap-frozen in liquid nitrogen in storage buffer [25 mM Tris-HCl (pH 7.5 at RT), 200 mM NaCl, 7 mM β-mercaptoethanol, 30% glycerol] and stored at -80 °C.

#### SsrA-SNAP

BL21(DE3) RP transformed with pET28-2×Strep-SsrA-SNAP were cultured overnight in LB broth supplemented with 50 µg/mL Kanamycin and 25 µg/mL Chloramphenicol (LB/Kan/Cam). Pre-culture was inoculated to LB/Kan/Cam and cultured at 37°C until OD_600_ reached around 0.5. Protein expression was induced by adding 300 μM IPTG and allowed to proceed for 20 hours at 23°C. The cells were resuspended with lysis buffer [50 mM Tris-HCl (pH 7.6 at RT), 200 mM NaCl, 1 mM MgCl_2_, 7 mM β-mercaptoethanol, 1 mM PMSF, 1× protease inhibitor cocktail V (Wako 168-26033)] supplemented with house-purified benzonase and sonicated by Misonix Ultrasonic Liquid Processor Sonicator S4000 (amplitude 30, 2 seconds ON, 10 seconds OFF, total 3 minutes ON). The lysate was centrifuged at 13,420×g at 4°C for 45 min to remove cell debris. The supernatant was applied to Strep-Tactin XT High Capacity (IBA Lifesciences) equilibrated with buffer L3 (25 mM Tris-HCl (pH 7.6 at RT), 200 mM NaCl, 1 mM MgCl_2_, 7 mM β-mercaptoethanol, 1 mM PMSF) and the protein was eluted with 100 mM biotin in wash buffer. The elution was dialyzed against buffer L3 and further purified by Superdex 200 increase 10/300 GL column in buffer L3. Peak fractions were collected and concentrated by Viva Spin Turbo (10 kDa cutoff). The protein was snap-frozen in liquid nitrogen in storage buffer (25 mM Tris-HCl (pH 7.6 at RT), 200 mM NaCl, 7 mM β-mercaptoethanol, 40% glycerol) and stored at -80 °C.

#### GST-pVCA-SspB-mCherry

The bacteria transformed with pGEX2T-GST-pVCA-3Csite-SspB(micro)-mCherry-TEVsite-6×His or pGEX2T-GST-pVCA-3Csite-SspB(nano)-mCherry-TEVsite-6×His were cultured overnight in LB broth supplemented with 100 µg/mL Ampicillin and 25 µg/mL Chloramphenicol (LB/Amp/Cam). Pre-culture was inoculated to LB/Amp/Cam and cultured at 37°C until OD_600_ reached around 0.5–0.6. After cooling the culture on ice, protein expression was induced by adding 300 μM IPTG and allowed to proceed for 24–26 hours at 18°C. The cells were resuspended with buffer V (50 mM HEPES-NaOH (pH 7.5), 150 mM NaCl, 10% glycerol, 14 mM β-mercaptoethanol, 1 mM EDTA, 1 mM PMSF, 100 µg/mL DNase, supplemented with cOmplete EDTA-free protease inhibitor cocktail) and lysed by microfluidizer. The lysate was centrifuged at 20,000×g at 4°C for 30 min to remove cell debris. To enhance resin binding, final 5 mM DTT was added to the lysate. The lysate was applied to glutathione sepharose equilibrated with buffer V [50 mM HEPES-NaOH (pH 7.5) 150 mM NaCl, 10% glycerol, 7 mM β-mercaptoethanol, and 1 mM PMSF], and the protein was eluted with 20 mM glutathione in buffer V (pH was adjusted between 7.0–8.0 by NaOH). The elution was then purified by Ni-NTA and eluted with 50–500 mM imidazole. After concentrated by Amicon Ultra (10 kDa cutoff), the protein was further purified by Superdex 200 increase 10/300 GL column in buffer V and concentrated again by Amicon Ultra (10 kDa cutoff). The protein was snap-frozen in liquid nitrogen and stored at -80 °C.

#### mCherry-SspB-mDia1

The bacteria transformed with pET28-2×Strep-mCherry-SspB(micro)-mDia1(FH1-FH2-DAD) were cultured overnight in LB broth supplemented with 50 µg/mL Kanamycin and 25 µg/mL Chloramphenicol (LB/Kan/Cam). Pre-culture was inoculated to LB/Kan/Cam and cultured at 37°C until OD_600_ reached around 0.5–0.6. After cooling the culture on ice, protein expression was induced by adding 300 μM IPTG and allowed to proceed for 24 hours at 16°C. The cells were resuspended with buffer D [50 mM Tris-HCl (pH 7.5), 500 mM NaCl, 10% glycerol, 1 mM DTT, 1 mM EDTA, 1 mM PMSF, 100 µg/mL DNase, supplemented with cOmplete EDTA-free protease inhibitor cocktail] and lysed by microfluidizer. The lysate was centrifuged at 20,000×g at 4°C for 30 min to remove cell debris. The supernatant was applied to Strep-Tactin XT (IBA Lifesciences) equilibrated with buffer D1 (50 mM Tris-HCl (pH 7.5), 500 mM NaCl, 10% glycerol, 1 mM DTT, 1 mM EDTA, 1 mM PMSF) and the protein was eluted with 50 mM biotin in buffer D1. The elution was concentrated by Amicon Ultra (10 kDa cutoff) and dialyzed against buffer D2 [25 mM HEPES-NaOH (pH 7.5), 150 mM NaCl, 7 mM β-mercaptoethanol, 10% glycerol]. The protein was further purified by Superdex 200 increase 10/300 GL column in buffer D2 and concentrated again by Amicon Ultra (10 kDa cutoff). The protein was snap-frozen in liquid nitrogen and stored at -80 °C. This protein was used as mDia1 in most figures unless otherwise indicated.

#### mCherry-SspB-mDia1-6×His and mCherry-SspB-mDia1ΔDAD-6×His

The bacteria transformed with pET28-2×Strep-mCherry-SspB(micro)-mDia1(FH1-FH2-DAD)-TEVsite-6×His or pET28-2×Strep-mCherry-SspB(micro)-mDia1ΔDAD(FH1-FH2)-TEVsite-6×His were cultured overnight in LB broth supplemented with 50 µg/mL Kanamycin and 25 µg/mL Chloramphenicol (LB/Kan/Cam). Pre-culture was inoculated to LB/Kan/Cam and cultured at 37°C until OD_600_ reached around 0.5–0.6. After cooling the culture on ice, protein expression was induced by adding 300 μM IPTG and allowed to proceed for 24 hours at 16°C. The cells were resuspended with buffer D3 [50 mM Tris-HCl (pH 8.0), 500 mM NaCl, 10% glycerol, 1 mM DTT, 1 mM PMSF] supplemented with 1× protease inhibitor cocktail V (Wako 168-26033)] and house-purified benzonase, and sonicated by Misonix Ultrasonic Liquid Processor Sonicator S4000 (amplitude 30, 2 seconds ON, 10 seconds OFF, total 3 minutes ON). Unbroken cells were pelleted by centrifugation at 13,420×g for 5 min at 4°C, resuspended in buffer D1, and subjected to sonication under the same conditions. This procedure was repeated twice. The lysate was then clarified by centrifugation at 13,420×g for 45 min at 4°C. The supernatant was applied to Strep-Tactin XT (IBA Lifesciences) equilibrated with buffer D3 and the protein was eluted with 100 mM biotin in buffer D3. The elution was then purified by Ni-NTA equilibrated with 50 mM Tris-HCl (pH 8.0), 150 mM NaCl, 10% glycerol, 7 mM β-mercaptoethanol, 1 mM PMSF and eluted stepwise with 20–500 mM imidazole. Peak fractions were collected and diluted 3-fold with 50 mM Tris-HCl (pH 8.0), 10% glycerol, 7 mM β-mercaptoethanol, 1 mM PMSF. The protein was further purified by HiTrapQ column equilibrated with 50 mM HEPES-NaOH (pH 7.5), 50 mM NaCl, 10% glycerol, 7 mM β-mercaptoethanol, 1 mM PMSF, and eluted with linear NaCl gradients of 50–500 mM NaCl. The protein was concentrated by Viva Spin Turbo (10 kDa cutoff), snap-frozen in liquid nitrogen and stored at -80 °C. mCherry-SspB-mDia1-6×His was used in Figure 2E, F, H, S10, S12. mCherry-SspB-mDia1ΔDAD-6×His was used in Figure 4J.

#### Capping protein

The bacteria transformed with pET28-6×His-2×Strep-TEVsite-Capβ2(mouse)-Capα1(mouse) were cultured overnight in LB/Kan/Cam. Pre-culture was inoculated to LB/Kan/Cam and cultured at 37°C until OD_600_ reached around 0.7. Protein expression was induced by adding 300 μM IPTG and allowed to proceed for 24 hours at 18°C. The cells were re-suspended with 50 mM Tris-HCl (pH 7.5 at RT), 150 mM NaCl, 14 mM β-mercaptoethanol,1 mM EDTA, 1 mM PMSF, 100 µg/mL DNase and lysed by microfluidizer. The lysate was centrifuged at 20,000×g at 4°C for 30 min to remove cell debris. The supernatant was applied to Strep-Tactin XT 4Flow (IBA Lifesciences) equilibrated with buffer C1 [50 mM Tris-HCl (pH 7.5 at RT), 150 mM NaCl, 7 mM β-mercaptoethanol] and the protein was eluted with 50 mM biotin in buffer C1. The elution was dialyzed against buffer C2 [25 mM Tris-HCl (pH 7.5 at RT), 150 mM NaCl, 7 mM β-mercaptoethanol]. 6×His-2×Strep tag was cleaved by 6×His-MBP-TEV protease (S219V) (purified in house) during the dialysis. The cleaved 6×His-2×Strep tag and TEV protease were removed through Ni-NTA column. Flow through and wash fractions were dialyzed against buffer and concentrated by Amicon Ultra (10 kDa cutoff). The protein was snap-frozen in liquid nitrogen in buffer C2 and stored at -80 °C.

### Pyrene-actin polymerization assay

For pyrene actin assay, actin was purified by size exclusion column. One milligram of actin (Cytoskeleton Inc. AKL99-B) was dissolved in 400 μL G-buffer [2 mM Tris-HCl (pH 7.5 at RT), 0.1 mM CaCl_2_, 0.2 mM ATP, 0.5 mM DTT, 1 mM NaN_3_] and dialyzed against the same buffer. After 3 days dialysis with daily buffer exchange, the actin was purified by Superdex 200 Increase 10/300 GL column (GE Healthcare) to separate monomer fraction from a fraction occasionally appear at larger size molecular size (smaller elution volume). The purified actin was stored at 4°C in a dialyzed manner against G-Buffer with twice a week buffer exchange.

Pyrene-actin polymerization assay was performed with FluoroMax 3 and DataMax software. Pyrene fluorescence and its kinetics were measured by 365 nm excitation (1 nm bandwidth) and 407 nm emission (5 nm bandwidth). The reaction was prepared by following the previously reported method (*135*), except that the reaction was 50 µL in quartz cuvettes (Hellma, 105-251-15-40).

### Ultracentrifugation assay

Reaction solutions were prepared by mixing 4.5 µL of actin in G-buffer and 4.5 µL of other proteins solution. At the final concentration, reaction solution contained 7.5 µM actin (10% Alexa 488 labeled), 1 µM GST-pVCA-SspB(micro)-mCherry, 1 µM mCherry-SspB(micro)-mDia1, 150 nM Arp2/3, 50 nM capping protein, 3 µM profilin, 2 µM cofilin, 1 µM creatin kinase, 1 mM ATP, 25 mM creatin phosphate, 240 mM sucrose, and 1×KMEI buffer [10 mM imidazole pH 7.0, 50 mM KCl, 1 mM MgCl_2_, 1 mM EGTA]. Polymerization reaction for 30 min at RT. The solutions were ultracentrifuged at 100,000×g (90,000 rpm, S55A2 rotor) for 1 hour at RT and analyzed by SDS-PAGE.

### GUV preparation by emulsion transfer

Lipid-in-oil solution was prepared as follows. Chloroform solutions of lipids were mixed in 5 mL glass vial, dried at 70°C with N_2_ gas flow, and left in vacuum 3 hours to overnight. The amounts of lipids were typically 6.5 µmol (1.3 mM, 1 mg/mL at final concentration in oil) for 40 mol% POPS/60 mol% POPC mixture (MARCKS peptide anchoring) and 1.5 µmol (0.3 mM, 0.23 mg/mL at final concentration in oil) for 2 mol% Benzylguanine-conjugated lipid/98 mol% POPC and 100% POPC (SNAP-tag anchoring). Lipid was dissolved in 5 mL of oil mixture (90 volume% hexadecane, 10 volume% silicone oil) by heating at 80°C for 1–2 hours, then brought to RT.

For iLID-SspB translocation experiments, inner solution contained 4 µM iLID-YFP-SNAP (or iLID-EYFP-MARCKS) and 1 µM mCherry-SspB(micro) in buffer (22.5 mM Tris-HCl (pH 7.5 at RT), 122.5 mM NaCl, 2 mM MgCl_2_, 240 mM sucrose, 0.7 mM β-mercaptoethanol, and 3% glycerol). Outer solution was prepared by replacing protein fractions of inner solution with their buffers and sucrose with glucose.

For actin polymerization experiments, inner solution was prepared by mixing 10 µL of actin (10% Alexa 647 labeled) in G-buffer and 10 µL of other proteins solution similarly to pyrene actin polymerization assay. At the final concentration, inner solution typically contained 11.75 µM iLID-EYFP-SNAP, 1 µM GST-pVCA-SspB(micro)-mCherry, 1 µM mCherry-SspB(micro)-mDia1, 7.5 µM actin, 150 nM Arp2/3, 50 nM capping protein, 3 µM profilin, 2 µM cofilin, 1 µM creatine kinase in the buffer (5.9 mM Tris-HCl (pH 7.5 at RT), 1.3 mM HEPES-NaOH (pH 7.5 at RT), 7.7 mM Imidazole (pH 8.0 at RT), 27 µM KH_2_PO_4_, 74 µM Na_2_HPO_4_, 39 mM KCl, 33 mM NaCl, 0.8 mM MgCl_2_, 43 µM CaCl_2_, 1.1 mM EGTA, 1.2 mM β-mercaptoethanol, 0.21 mM DTT, 0.43 mM NaN_3_, 1 mM ATP, 25 mM creatine phosphate, 240 mM sucrose and 2.8% glycerol). Outer solution was prepared by replacing protein fractions of inner solution with their buffers and sucrose with glucose. Protein and buffer conditions are summarized in Supplementary table 1.

In experiments using pVCA alone, SspB(nano) was employed to ensure sufficiently robust light-induced membrane recruitment and actin polymerization, and pVCA was used at 4–4.2 µM. When mDia1 was included, SspB(micro) was used to avoid excessive basal membrane association of mDia1 prior to illumination. To maintain comparable recruitment behavior within the combined system, pVCA was also fused to SspB(micro) and used at 1 µM, matching the concentration of mDia1.

GUVs were prepared as previously reported (*56, 136*). Briefly, oil-buffer interface was created by layering 250 µL lipid-in-oil solution on top of 250 µL outer buffer in 1.5 mL tube. Water-in-oil emulsion was made by vigorously pipetting 20 µL of inner solution in 250 µL of lipid-in-oil solution and then gently added to the top of the oil-buffer layers. The emulsions were transferred through the oil-buffer interface by centrifugation at 2500×g for 2 min at RT. GUVs were collected by pipetting.

### Image acquisition and analysis

For imaging GUVs, 8-well glass chamber slides (ThermoFisher Scientific, 154534) were pretreated with 10 mg/mL BSA in PBS for 10 minutes and washed with MilliQ water twice. LSM780 confocal microscope (Zeiss) equipped with Plan-Apochromat 63X/1.40na oil immersion DIC objective lens (Zeiss 420782-9900) was used for imaging and light stimulation of GUVs. YFP, mCherry, and Alexa Fluor 647 were imaged with 514, 561, and 633 nm excitation lasers, respectively. Blue light illumination for iLID stimulation was performed by using bleaching function with 458 nm laser at intensity 1.0–5.0%, scan speed 6 (pixel dwell time = 6.3 µsec). Main beam splitters MBS458/561, MBS 488/561/633, and MBS 458/514 were used for 458 and 561, 633, and 514 nm excitation, respectively. The imaging was performed at room temperature. Data were analyzed by Fiji software (*137*).

### FRAP analysis on GUV membranes

A small region of interest (ROI) on the GUV membrane was selectively photobleached for 1 s using high-intensity laser illumination. Fluorescence recovery within the bleached ROI was measured. To correct for photobleaching caused by repeated imaging, fluorescence intensity in the bleached ROI was normalized to that of a non-bleached reference region on the same GUV membrane. To capture rapid fluorescence recovery with short imaging intervals, mCherry- and Alexa Fluor 647–labeled signals were analyzed on different individual GUVs.

### Small unilamellar vesicles (SUVs) preparation

SUVs used for EM experiments were prepared as follows. Chloroform solutions of lipids were mixed in 5 mL glass vial, dried at 70°C under a flow of N_2_ gas, and left under vacuum overnight. The total amount of lipid was 0.5 µmol (corresponding to 5 mM, or 4 mg/mL, at the final concentration in buffer). Lipid composition was 2 mol% benzylguanine-conjugated lipid/98 mol% POPC. The lipid film was hydrated with 100 µL of SUV buffer (25 mM HEPES-KOH (pH 7.6), 50 mM KCl, 1 mM MgCl_2_, 1 mM TCEP), incubated at 37°C for 10 min, and sonicated using Misonix Ultrasonic Liquid Processor Sonicator S4000.

### Electron microscopy (EM)

The reaction for EM imaging was prepared by mixing 2.75 µL of a solution containing all components except actin (solution I) with 2.25 µL of actin in G-Mg buffer [2 mM Tris-HCl (pH 7.5), 0.2 mM ATP, 0.1 mM MgCl_2_, 0.5 mM DTT, 1 mM NaN_3_] (solution II). Solution I was first placed to carbon-coated collodion film on a Cu grid (Nisshin EM, Tokyo, Japan) and incubated for several seconds to allow SUVs to settle onto the grid surface. Solution II was then added, and actin polymerization was allowed to proceed for 3 min at room temperature (24–26°C). At final concentrations, the reaction mixture contained 0.3 mg/mL lipid SUVs, 7.5 µM actin (10% Alexa Fluor 647–labeled), 1 µM GST–pVCA–SspB(micro)–mCherry, 1 µM mCherry–SspB(micro)–mDia1(FH1–FH2–DAD)–6×His, 150 nM Arp2/3 complex, 50 nM capping protein, 3 µM profilin, 2 µM cofilin, 1 µM creatine kinase, 1 mM ATP, 25 mM creatine phosphate, and 1× KMEI buffer (10 mM imidazole, pH 7.0, 50 mM KCl, 1 mM MgCl₂, 1 mM EGTA). After the reaction, excess liquid on the grid was carefully removed, followed by negative staining with 2% phosphotungstic acid (pH 7.0). The samples were observed with a JEM1010 EM (JEOL, Akishima, Japan) at 80 kV equipped with a FastScan-F214(T) charge-coupled device (CCD) camera (TVIPS, Gauting, Germany).

### Live-cell CID experiments

DNA constructs: Four plasmids were used which encoded Lyn-CFP-FRB (*138*), YFP-FKBP (*138*), YFP-FKBP-Tiam1 (*133*), YFP-FKBP-pVCA, and YFP-FKBP-mDia1.

Cell culture: HEK293T cells (ATCC) were cultured in DMEM (Corning, 10-013-CV) supplemented with 10% fetal bovine serum (Corning, 35-010-CV) and 1% penicillin-streptomycin (ThermoFisher, 15140163).

Transient transfection: HEK293T cells were seeded at a density of 9,000-12,000 cells in an 8-well chamber slide. Per well, a solution of 0.3 μg of DNA, 1 μL of FuGENE HD, and 30 μL of Opti-MEM was added for transfection. FRB:FKBP pairs were cotransfected in a 1:1 or 1:2 ratio. Cells were incubated at 37°C with 5% CO_2_ and 95% humidity for 24–48 h before imaging.

Microscopes and imaging: Live cell imaging of actin NPF recruitment to the cell membrane was performed using an Eclipse Ti inverted fluorescence microscope (Nikon) with a 100x Oil objective lens and Andor Zyla 4.2 plus sCMOS camera, or with a 60x Oil, TIRF 60x Oil, or TIRF 100x Oil objective lens and ORCA-Fusion Digital CMOS camera. Images were captured every 1 min for 30-120 min, where within 5 min of starting, 0.1 μM rapamycin was added. Nikon microscopes were driven by NIS-Elements software (Nikon). All live-cell imaging was performed at 37 °C with 5% CO_2_ and humidity control by a stage top incubator. Image processing and analysis were performed using Fiji software.

Data analysis: Per transfection sample, a count was made based on the phenotype of cells falling into three categories: no change, lamellipodia, and filopodia. Cells with lamellipodia or filopodia pre-rapamycin were not counted unless new membrane protrusions formed after rapamycin addition. If the cell under observation did not produce any new lamellipodia or filopodia after addition of rapamycin, then it was counted as no change. Only healthy cells fully in the field of view with clear expression of both FRB and FKBP constructs were counted. If there was any doubt in the phenotype of the cell, then it was not included in the count.

#### Statistics

1. The cell counts for each category were summed across all experiments, and this sum was divided by the total number of cells to get a percentage of cells belonging to each category when a certain actin NPF was recruited to the membrane.
2. For each image set collected from a transfection sample, the categorized cell counts were divided by the total cell count to get percentages. For each actin NPF, the percentages from all transfection samples were averaged. These averages were analyzed by two-tailed Student’s t-test assuming equal variance (determined by F-test) in order to establish the significance of lamellipodia or filopodia detection between transfection samples.

### Statistics and reproducibility

Quantitative data with error bars were obtained from three or more independent experiments, except for fig. S10 and S11, which were obtained from two independent experiments. Where indicated, 95% confidence intervals of the mean were calculated in Microsoft Excel using the t-distribution (T.INV.2T) and the standard error (SD/√n). For statistical comparison of multiple groups, the Steel–Dwass test was performed using the pSDCFlig function (asymptotic option) in the NSM3 package in R. Pearson’s R and p-values in fig. S19 was calculated by cor.test function (pearson option).

### Theoretical model of actin filament-membrane interactions

We employed Cytosim (*95*), a stochastic agent-based cytoskeletal modeling framework that incorporates Brownian dynamics, actin filament dynamics, and the action of actin-binding proteins. To Cytosim, we added a deformable membrane model (*96*) along with terms for the reciprocal actin-membrane interactions (*97*). In the following text we elaborate on the details and implementation for each model component. We further refer the reader to Table 1, for a complete list of parameters and their corresponding references from where they were obtained. The use of these physically relevant parameters constraint our simulations against reality.

**Table 1.**
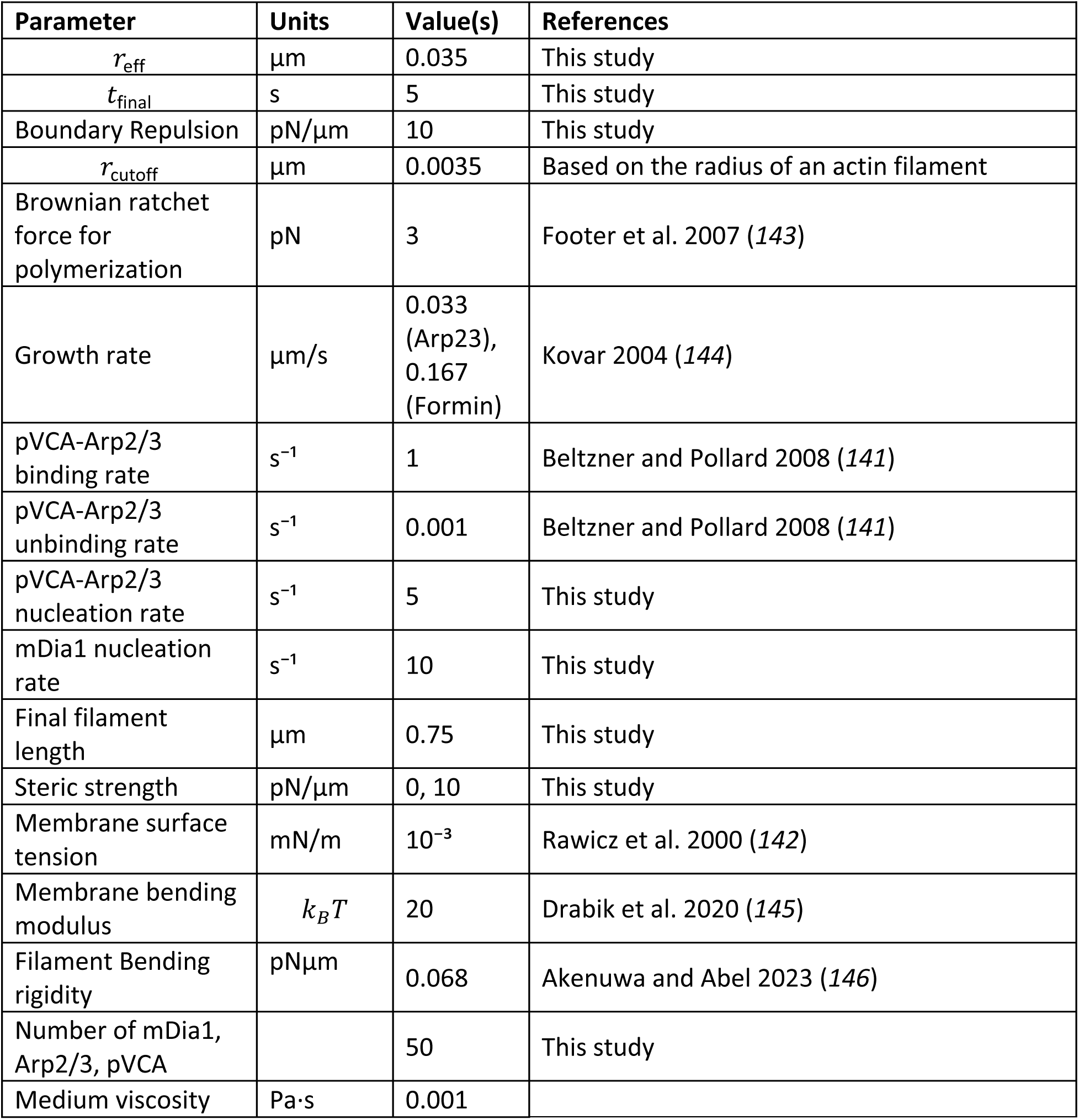
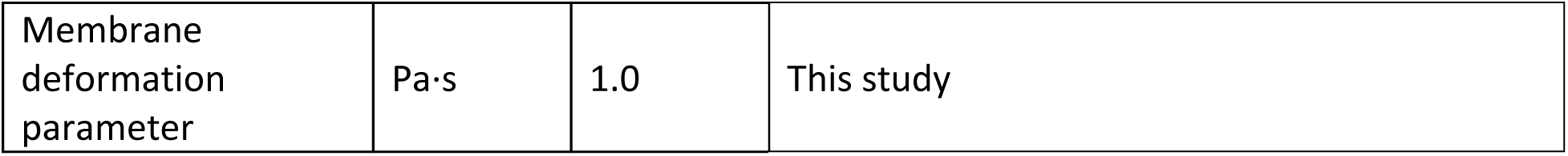
List of Parameters used in the simulations.

To model the membrane, we utilize a 2-D enclosed polygon with resistance to bending and stretching. The energy of the membrane is given by discrete 2-D Helfrich Hamiltonian,

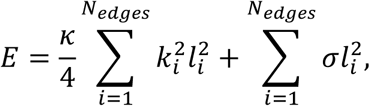

where *κ* is the bending rigidity of the membrane and *σ* is the membrane surface tension. *k*ᵢ is the local curvature at vertex *i* and *l*ᵢ is the length of the edge connecting vertices *i* and *i*+1. The force on a vertex *i* is given by

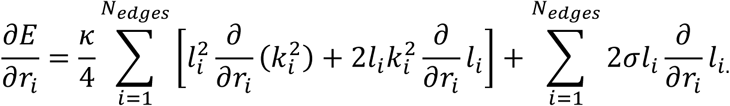

There are several established ways to define curvature in discrete settings (*139, 140*). For this model, we choose a definition of curvature based on the turning angle at each vertex.

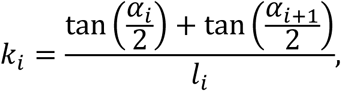

where the turning angle *α*ᵢ is given by the sum of the internal angles,

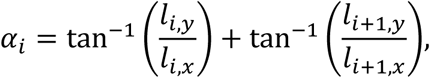

and we can express ∂*l*ᵢ/∂*r*ₐ and ∂*α*ᵢ/∂*r*ₐ as

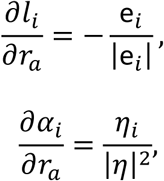

respectively, as the unit vector along the edge *i* and the unit normal to the edge *i*.

### Filament description

The actin filaments are modeled as described in the original Cytosim reference (*95*). In brief, actin filaments are modeled as semiflexible, inextensible fibers composed of linear segments with hinge points that allow for bending. The bending rigidity of the filament, 𝑘*_bend_*, is chosen so that the persistence length of an isolated filament is 17 𝜇m.

### Filament-membrane interactions

We model filament-membrane interactions using a purely repulsive harmonic potential between points on the filament and the edges of the enclosed polygon. Given a position **p** on a filament *i* and an edge *j* on the enclosed membrane, the force is given by

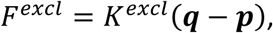

where 𝒒 is the point closest to the point 𝒑 on the edge *j* and 𝐾*^excl^*is the stiffness of the steric potential.

𝐹*^excl^* confines the actin filament within the enclosed membrane and applies a force from the actin filament to the membrane. The steric force to points on the membrane edge 𝑐_𝑗_ and 𝑐_𝑗+1_ is calculated using the lever rule,

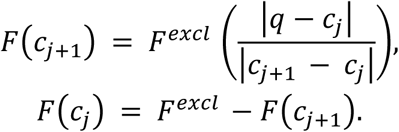

### Inter-filament interactions

They are implemented as a repulsive harmonic potential between filaments, mimicking the physical constraints and forces arising from their proximity and entanglement.

### Dynamics of filament growth

We assume that filaments polymerize at their barbed ends and do not depolymerize at the pointed ends. We assume a constant polymerization rate dependent on which actin-binding protein is bound to the filament.

### Actin-binding proteins and nucleation promoting factors

We considered three proteins: mDia1 (formin) and Arp2/3. To model the spatial control of mDia1 and pVCA, we assume that iLID-SspB is already attached to pVCA and mDia1 at the membrane surface. To capture the actin nucleation and polymerization dynamics of the experiment, we modeled mDia1 and pVCA as beads that can nucleate and elongate actin filaments at prescribed rates. For reasons of computational cost, we assume that pVCA recruitment of Arp2/3 is fast. It has been shown experimentally that pVCA unbinds Arp2/3 after nucleation (*141*). In our model, pVCA unbinds the mother filament after nucleation. After initial nucleation by pVCA-Arp2/3 complex, free Arp2/3 can bind to mother filaments and nucleate daughter filaments. Arp2/3 stochastically bind mother filaments with a rate 𝑘*_bind_* and unbind with a rate 𝑘*_unbind_* in a force-dependent manner given by

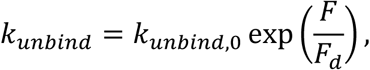

where 𝑘*_unbind,0_* is the unbinding rate in the absence of force, **F** is the force experienced by the actin-binding protein and 𝐹*_d_* is a characteristic detachment force.

### Dynamics of network evolution

Unbound actin-binding proteins, filament motion, and membrane deformations are governed by Langevin dynamics. In each time interval Δ*t*, we update the positions of all particles in the system using

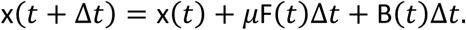

Here, *μ* is the viscosity of the medium. **F**(*t*) is the sum total of all forces acting on the particle at time *t*. **B**(*t*) is the Brownian noise term, which is a random variable drawn from a normal distribution with a mean of 0 and a variance of 2𝜇𝑘_𝐵_𝑇Δ𝑡. 𝑘_𝐵_is the Boltzmann constant and *T* is the temperature.

The forces on the membrane are composed of forces from membrane surface tension, bending rigidity, and point forces from the actin filaments on the membrane. In our formulation, the speed at which membrane deformation occurs is governed by a membrane deformation parameter, 𝜇*_mobility_* (not to be confused with the viscosity of the medium), which acts as a dampener to slow down the deformation of the membrane over time.

### Model Assumptions

To capture the length control effect of capping protein in the experiments, we cap filament growth to a particular filament length. Our current model does not account for filament turnover, a feature we plan to incorporate in future simulations. Because we are simulating the 2-D projection of the membrane, we assume that the membrane is in a low-tension regime and any stretching of the membrane during deformation results from overcoming entropic and enthalpic cost of unfolding of wrinkles. Membrane surface tension is set to 0.001 mN/m in line with experimentally reported values (*142*).

For computational cost and technical challenges of performing simulations on the timescales of the experiments, we assumed faster nucleation rates for mDia1 and pVCA-bound Arp2/3 and simulated for shorter timescales.

### Simulation set-up

Mimicking the instance of asymmetric surface accumulation after illumination, we randomly placed surface-bound nucleators at the right edge of the model GUV and simulated the system for a period of 5 s. We examined three cases: one with pVCA and Arp2/3, one with mDia1 only and one with both NPFs and Arp2/3. For each case, we simulated conditions with and without inter-filament interactions, giving a total of 6 simulation conditions. We performed 20 independent trajectories per condition.

### Quantification of membrane deformation in simulations

To quantify the extent of actin-induced membrane deformation, we calculated the average difference between the radius of the GUV and the distance from each node in the leading edge of the membrane to the center of the membrane at the final timepoint.

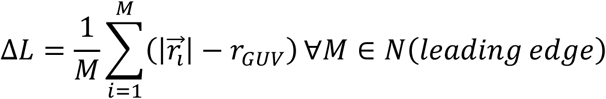

## Acknowledgement

We thank Yubin Zhou for TEV protease plasmid; Masato Kanemaki for pET28-2×Strep-tag plasmid; Miho Iijima for pGEX-2T plasmid; Bin Wu for Phage-ubc-nls-ha-tdMCP-SNAP plasmid; Roberto Dominguez for pCold-mDia1 plasmid, Saki Takayanagi for constructing GST-pVCA-FKBP-EYFP plasmid. We also thank Yusuke Sato, Taro Toyota, Pushpendra Singh, Tetsuya Kotani, Takeya Masubuchi for technical advice on the experiments. We appreciate Erin D. Goley for technical support of pyrene assay and insightful comments on actin preparation. We also appreciate Julie Plastino and Cecile Sykes for pET3d-Capβ2(mouse)-Capα1(mouse) plasmid and insightful advice on protein purification. Our appreciation extends to Pablo A. Iglesias for fruitful discussion. We thank our lab members, including Allen Kim, Abhijit Deb Roy, Helen D. Wu, Yuta Nihongaki for constructive discussion. We thank Kyoko Chiba for technical support on protein purification. We also thank Robert DeRose for proofreading the manuscript and experimental support. FRIS CoRE (a shared research environment in Tohoku University) is also acknowledged. This study was supported by the PRESTO program (JPMJPR12A5 to TI, JPMJPR20KA to HTM), CREST program (JPMJCR19S5 to MM), and FOREST program (JPMJFR2416 to HTM) of the Japan Science and Technology Agency, MEXT/JSPS KAKENHI Grants (23K14150, 23H04397, 25K02234, 25H02347 to HTM), National Institutes of Health (5R01GM123130, R01GM136858, R35GM149329 to TI, and R35GM128786 to BC), ExCELLS project research (25EXC604-6 to HTM) and the National Science Foundation (000819255 to TI, JL, BC). TI was also supported by Nitto Denko Corp. HTM was supported by Postdoctoral Fellowships from the Japan Society for the Promotion of Science. SR was supported by the National Science Foundation and the Japan Society for the Promotion of Science (JSPS) East Asia and Pacific Summer Institutes and Johns Hopkins Lucille Elizabeth Hay graduate fellowships. Computer simulations were run on hardware hosted by the Triton Shared Computing Cluster (*147*). OHA and CTL further acknowledge high performance computing support from the UCSD Physics Computing Facility.

## Author contributions

HTM, SR, and TI conceived the project. HTM performed most of the experiments and analyzed the data with contributions from SR, HN, BC, SM, SMN and TI. SR established GUV preparation method. WR performed in-cell CID experiment and analyzed the data. RY performed biochemical experiments. YOT performed EM under supervision by MM. OHA and CTL built and analyzed computational models. DN, DAK, and BC provided cytoskeleton proteins. TM provided benzylguanine-conjugated lipid. HTM wrote the manuscript in consultation with TI. HTM, SR, and TI edited the manuscript. All the authors contributed to the final version of the manuscript.

## Data Availability

The datasets generated during this study are available from the corresponding author upon request.

## Figure Legends

**Fig. S1.**
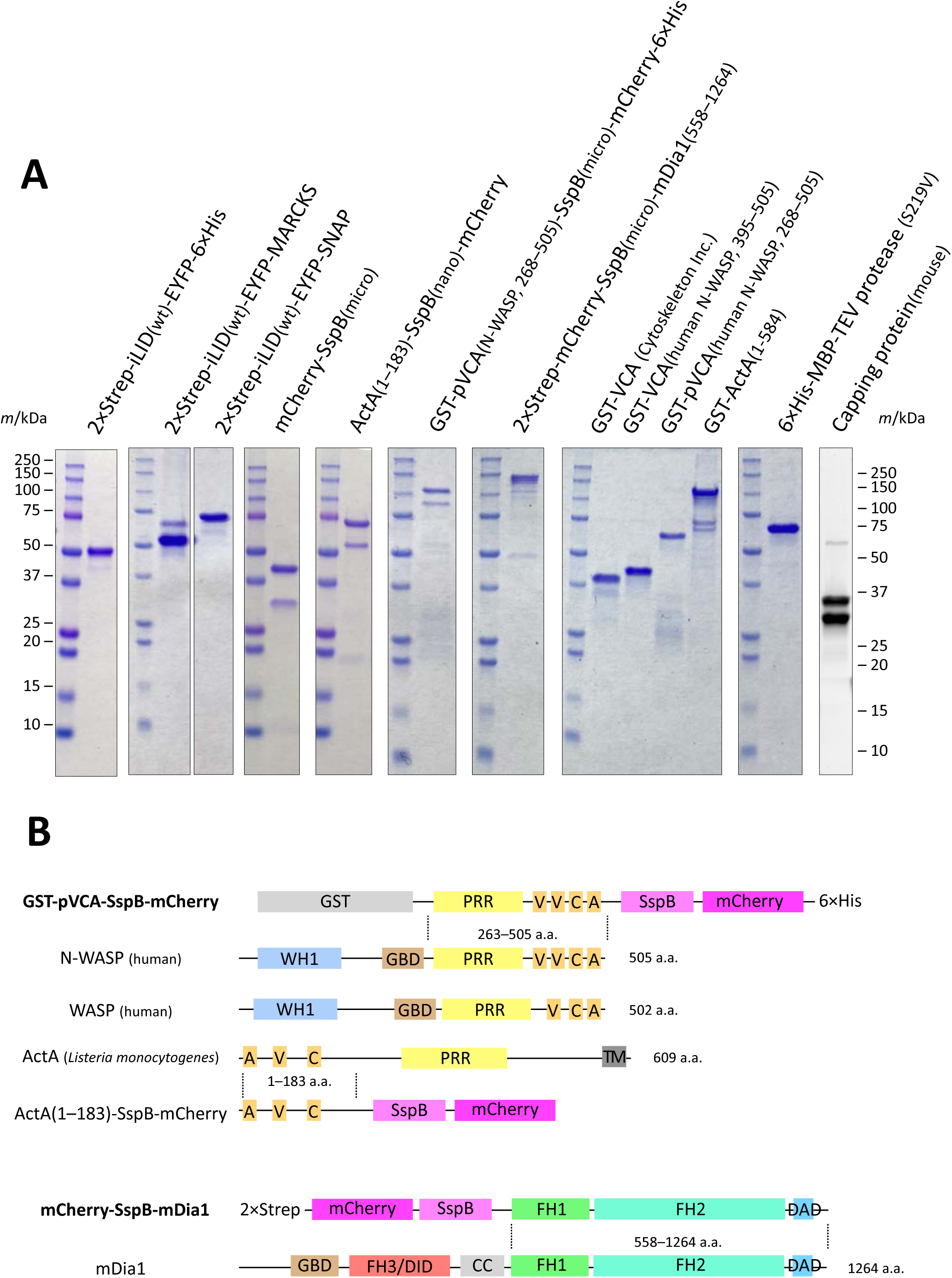
Proteins purified in this study. (A) SDS-PAGE results of purified proteins. Blue: Coomassie brilliant blue staining. Gray: SYPRO Ruby staining. Most mCherry-tagged proteins show two extra bands due to a known chromophore cleavage reaction during the boiling process (*148, 149*). (B) Domain structures of NPFs used in this study.

**Fig. S2.**
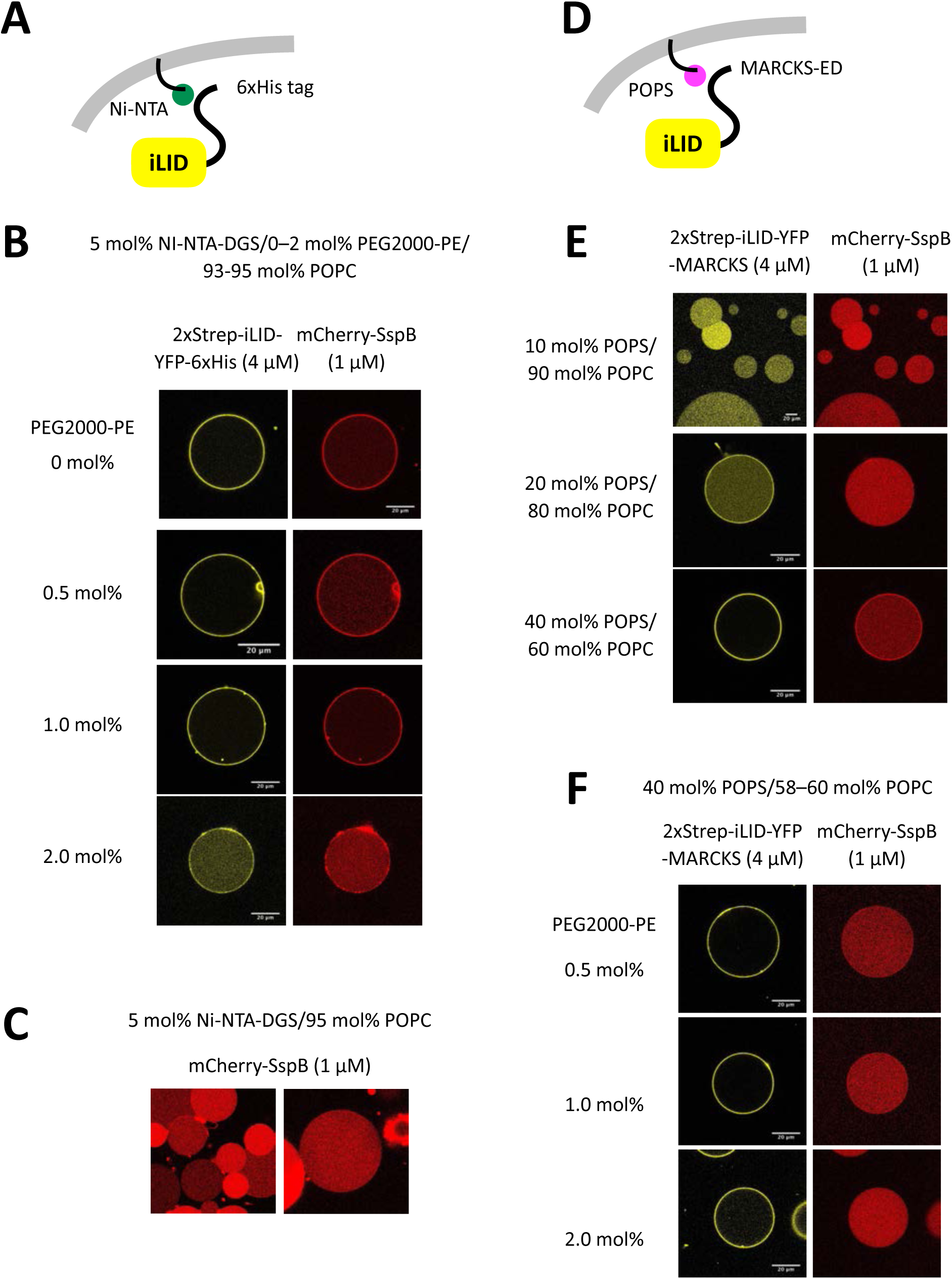
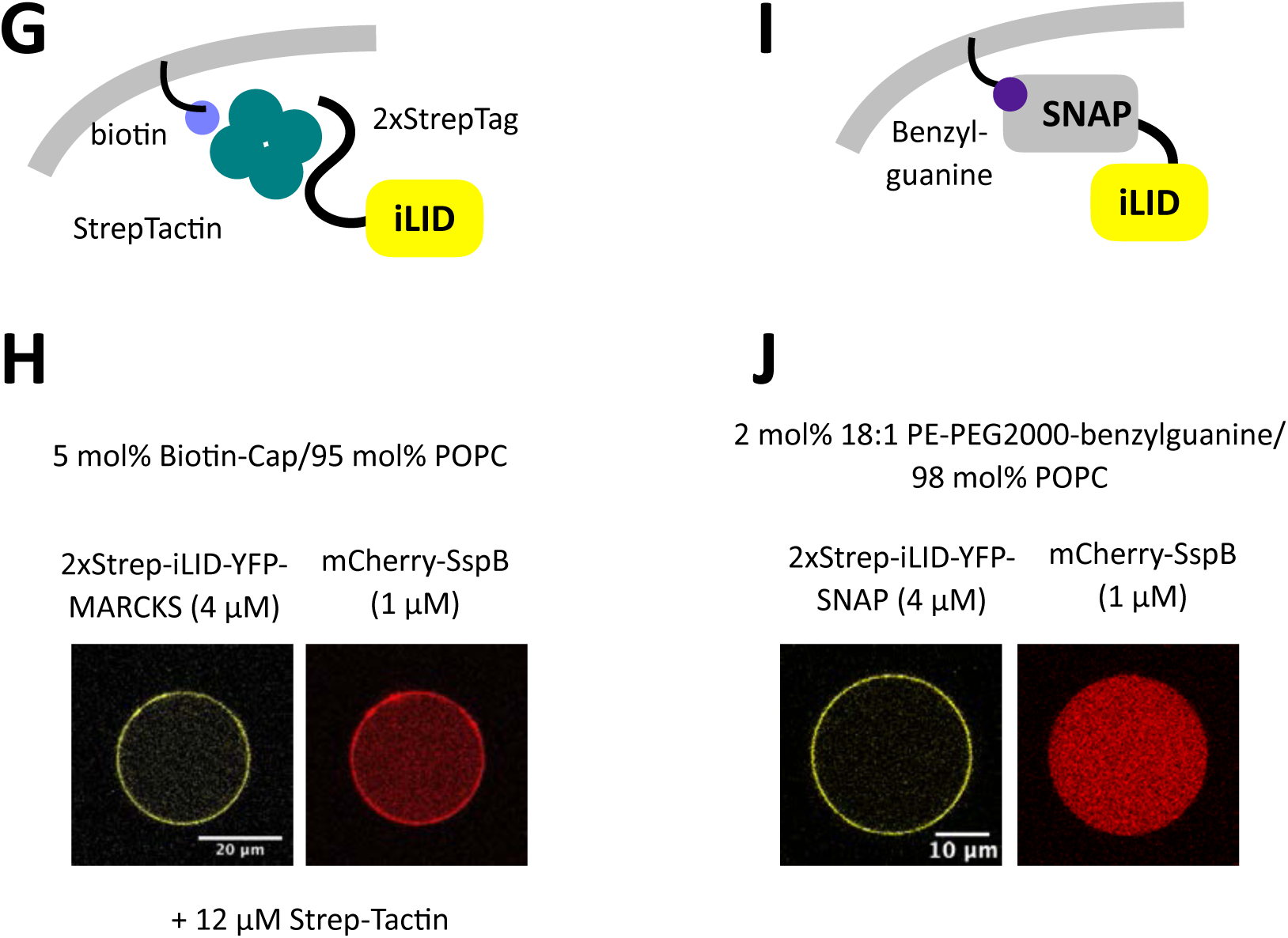
Optimization of membrane anchoring of iLID. (A),(D),(G),(I): Schematic representations of membrane anchoring of iLID. (B),(C),(E),(F),(H),(J): Confocal images of iLID variants and mCherry-SspB. (B) His-tag Ni-NTA pair caused non-specific membrane recruitment of SspB. This is possibly caused by the presence of 2×Strep-iLID-YFP-6×His because mCherry-SspB itself did not show non-specific binding to the membrane of 5%Ni-NTA DGS/95%POPC (C). (E),(F) MARCKS-ED peptide requires high percentage of PS and PEGylated-lipids to reduce non-specific binding of SspB. (H) Biotin-StrepTactin-Streptag also caused non-specific membrane binding of SspB. (J) The data correspond with Fig. 1B.

**Fig. S3.**
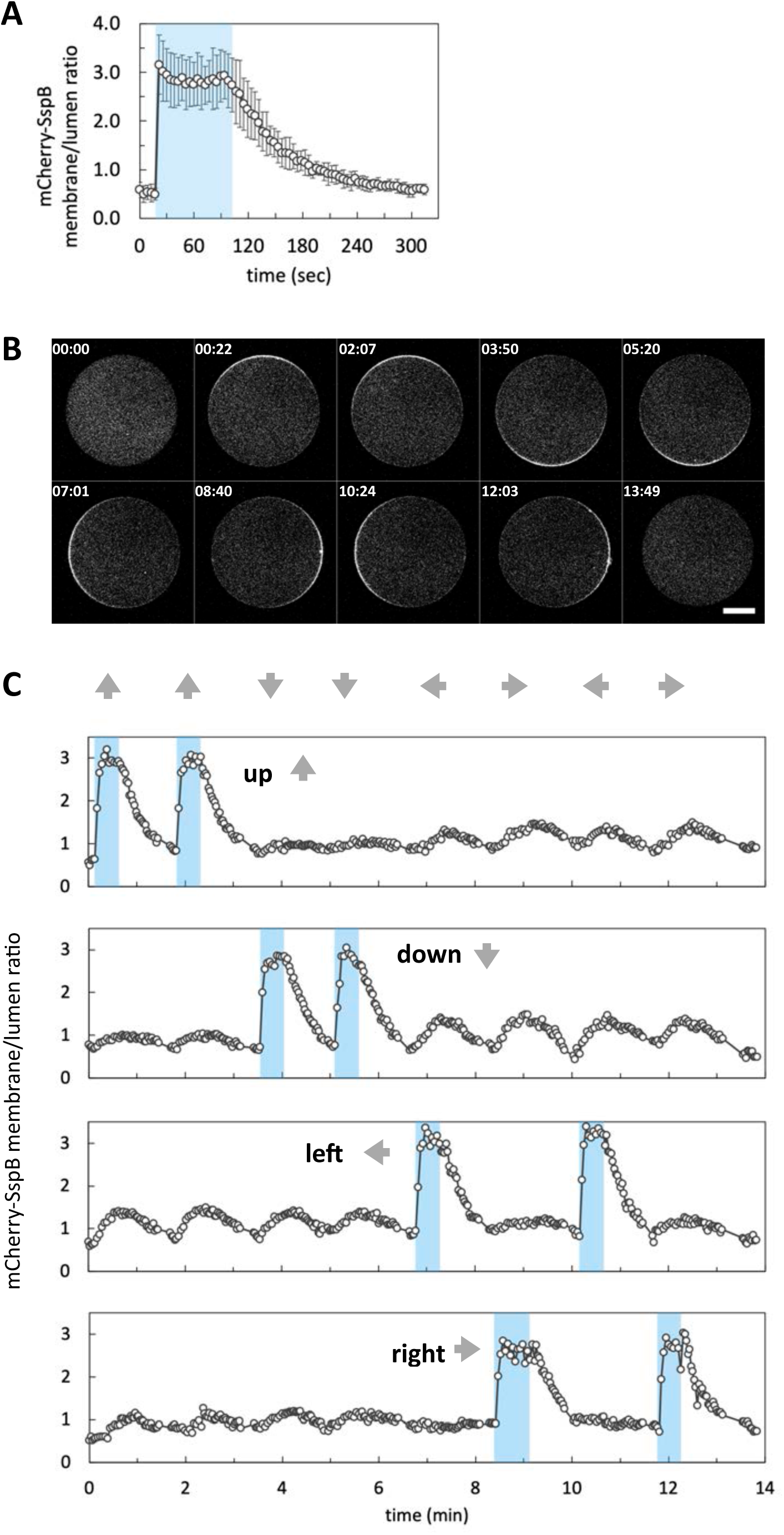
Asymmetric control of protein localization with iLID-SspB within GUVs. (A) Time course of membrane/lumen ratio of mCherry-SspB signal in the illuminated area (n=6). Error bars indicate 95% CI. Blue area indicates the time window of blue light stimulation. The data corresponds to Fig. 1E and F. (B, C) Confocal images and time course of membrane/lumen ratio of mCherry-SspB signal responding to repetitive local light illuminations. The data corresponds to Fig. 1G.

**Fig. S4.**
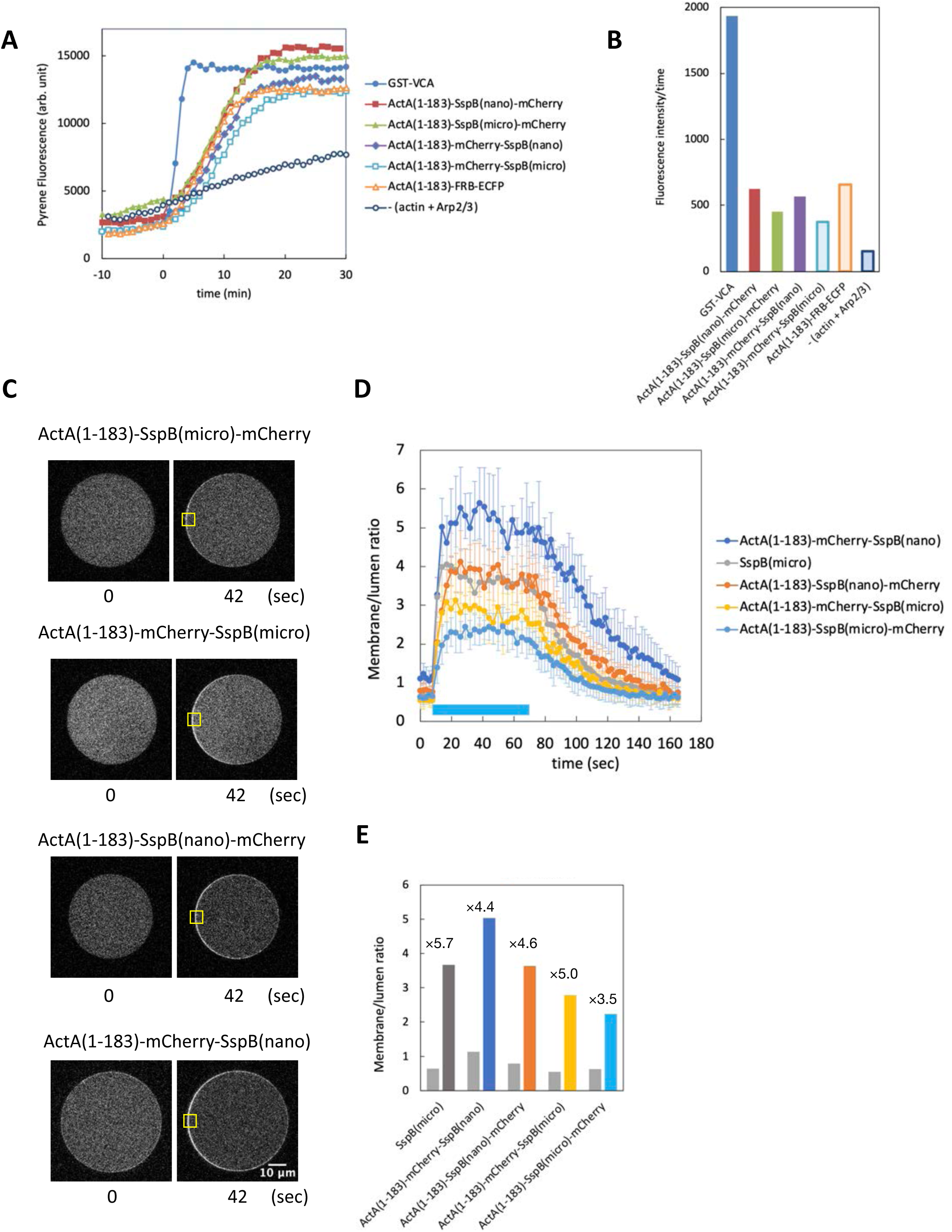
Optimization of ActA-SspB. (A) Pyrene actin polymerization assay. The reaction was performed with 1 µM actin (5% pyrene labeled), 10 nM Arp2/3, and 100 nM ActA variants or GST-VCA. (B) Fluorescence intensity/time in 0–6 min of (A) was plotted as bar chart. (C–E) Light-inducible local membrane recruitment of ActA (C) Confocal images of ActA translocation. Yellow boxes indicate the area of blue light illumination. (D) Time course of membrane/lumen ratio of ActA-SspB variants (n≥5 for each variant). Error bars indicate 95% CI. Blue bar indicates the time window of blue light stimulation. (E) Average membrane/lumen ratio of mCherry signal. In each condition, left and right bars indicate before and after light stimulation, respectively. The numbers on the bar chart indicate the fold change after stimulation.

**Fig. S5.**
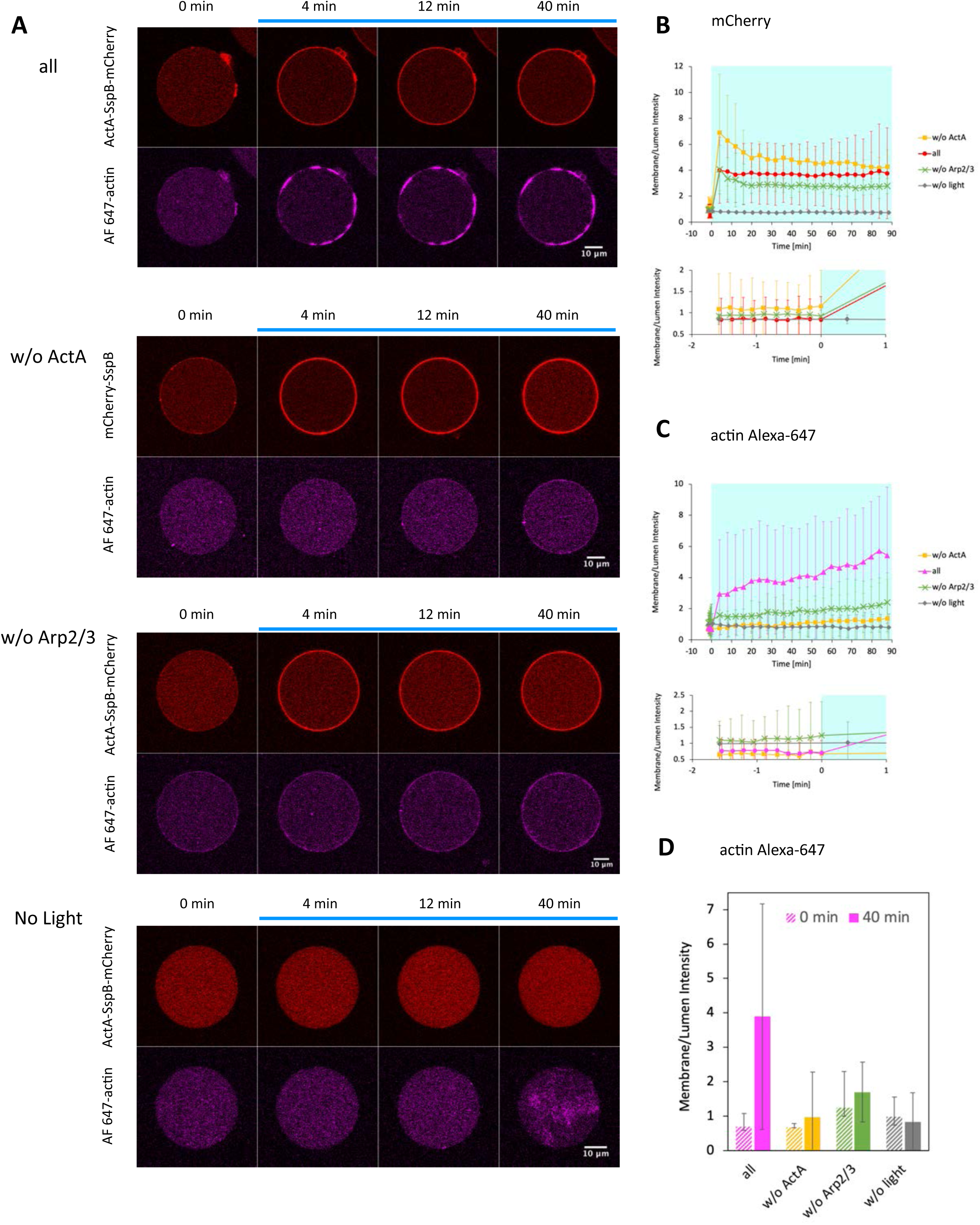
Light-inducible actin polymerization with global ActA membrane recruitment. iLID-EYFP-MARCKS (5 µM), ActA-SspBnano-mCherry (2 µM), actin (1.5 µM, 10% Alexa-647 labeled), and Arp2/3 (150 nM) were encapsulated in GUVs containing 60 mol% POPC/40 mol% POPS. Blue light was applied to the entire GUV. (A) Representative images of ActA(1–183)-SspB(nano)-mCherry and actin Alexa-647. All components (n=4): w/o ActA: ActA(1–183)-SspB(nano)-mCherry was replaced with mCherry-SspB(micro) (n=3), w/o Arp2/3: Arp2/3 was omited (n=3). No Light: no blue light stimulation (n=3). (B, C) Time course of membrane/lumen ratio of mCherry and actin Alexa 647. (D) Bar chart of membrane/lumen ratio of actin Alexa 647 before (0 min) and after (40 min) light stimulation. Error bars indicate 95% CI.

**Fig. S6.**
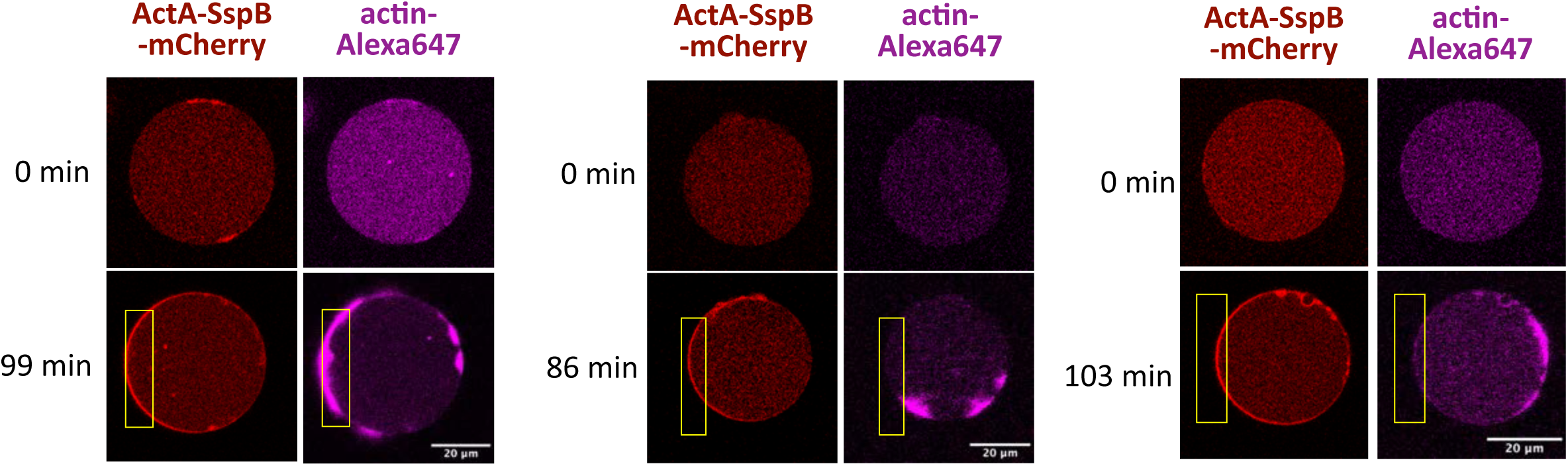
Asymmetric ActA membrane recruitment fails to induce directed actin polarization toward local light illumination. iLID-EYFP-MARCKS (5 µM), ActA-SspBnano-mCherry (2 µM), actin (1.5 µM, 10% Alexa-647 labeled), and Arp2/3 (150 nM), 1 mM ATP were encapsulated in GUVs containing 60 mol% POPC/40 mol% POPS. Yellow boxes indicate the area of blue light illumination. Although ActA creates asymmetric distribution, actin patches randomly distributed due to the diffusion on the membrane.

**Fig. S7.**
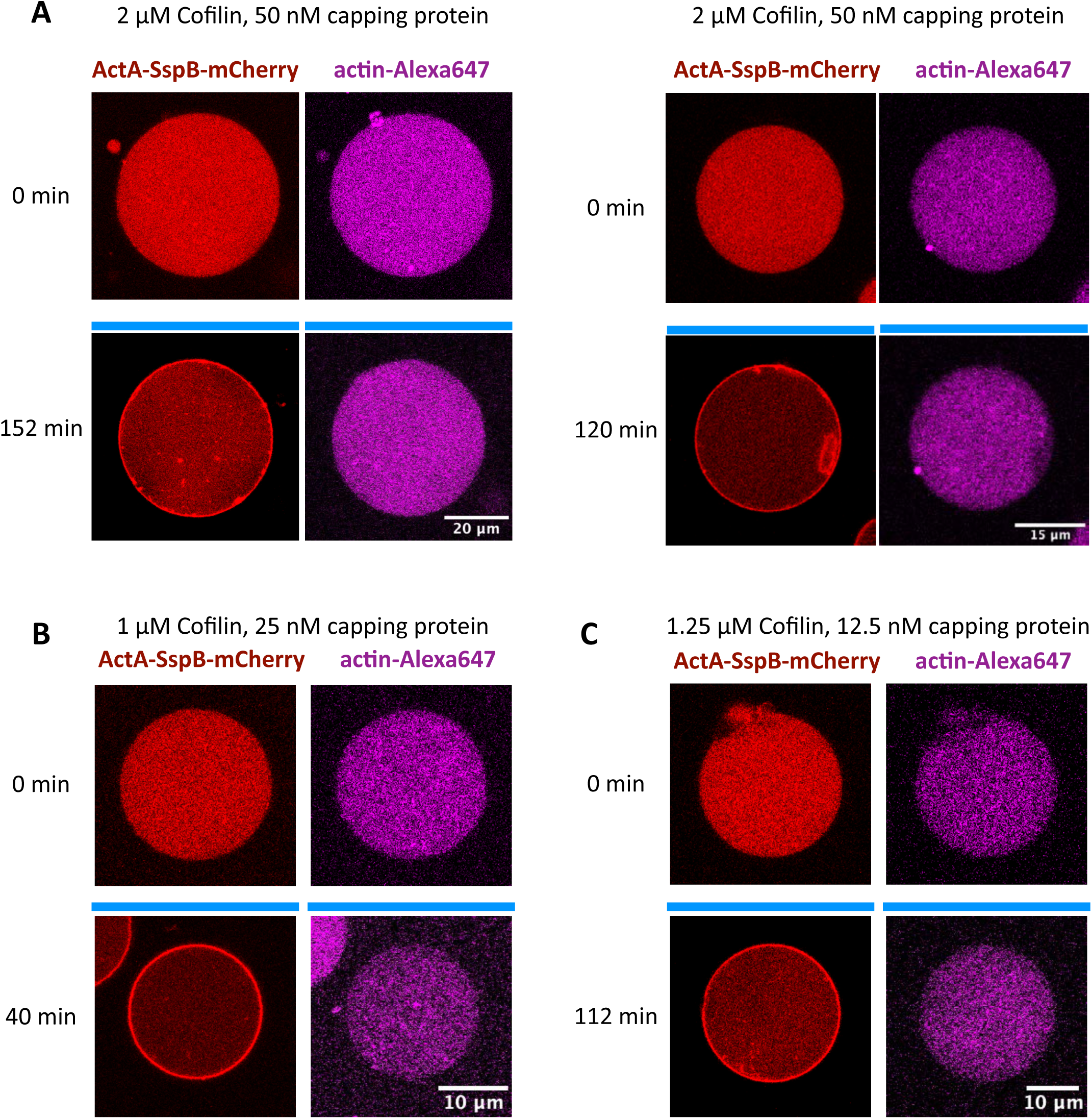
Global ActA membrane recruitment fails to induce actin polymerization on the membrane in the presence of profilin, cofilin, and capping protein. 15 μM iLID-YFP-MARCKS, 5 μM ActA(1-183)-SspB(nano)-mCherry, 7.5 μM actin (10% Alexa 647 labeled), 150 nM Arp2/3, 3 μM profilin, cofilin, and capping protein were encapsulated in GUVs containing 60 mol% POPC/40 mol% POPS. Blue bars indicate the period of blue light illumination.

**Fig. S8.**
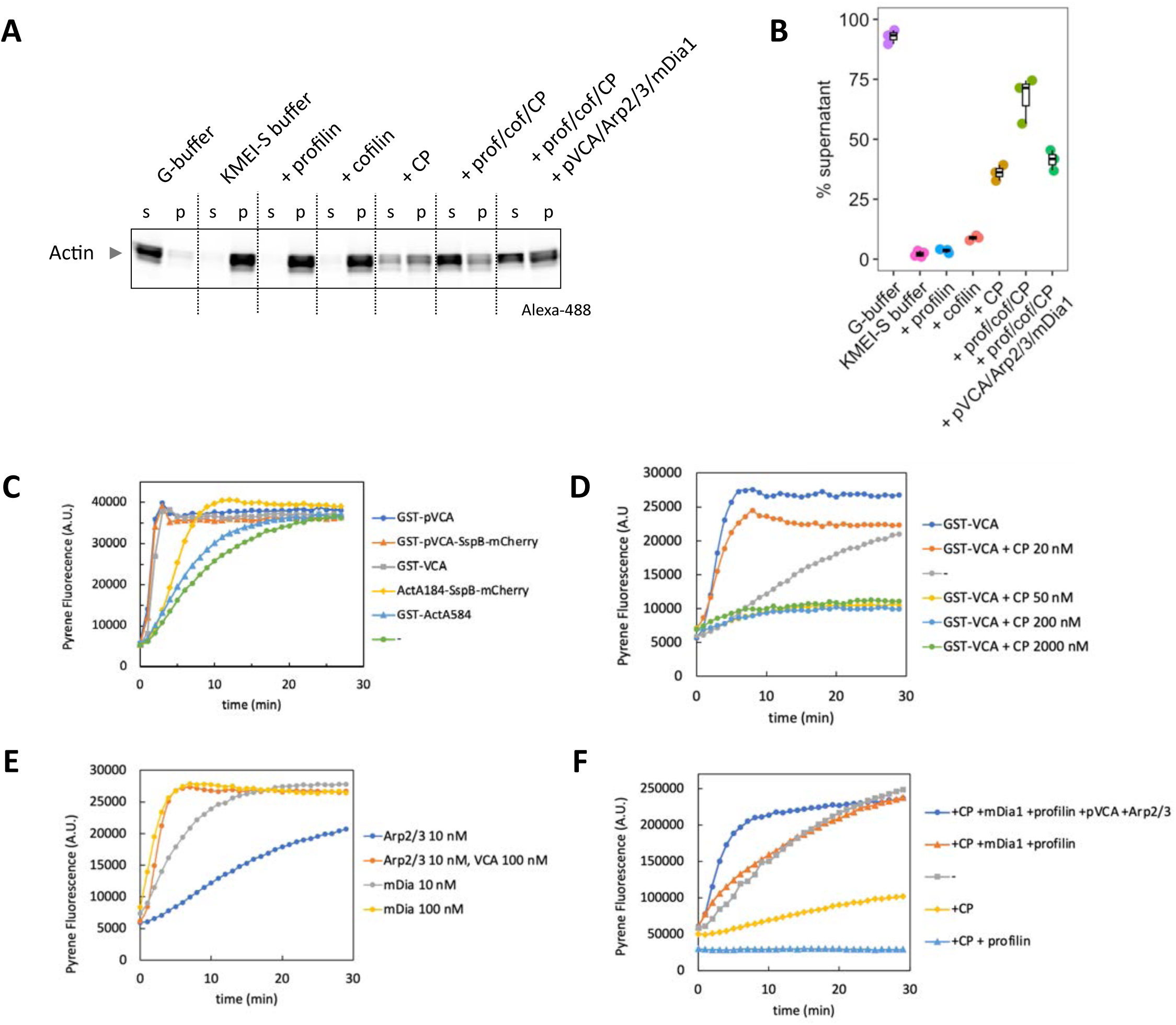
Biochemical characterization of actin and actin-binding proteins used in this study. (A, B) Ultracentrifuge assay. After 30 min actin polymerization reaction at RT, the solutions were ultracentrifuged 100,000×g, 60 min, at RT. Proteins and concentrations used: 7.5 µM actin (10% Alexa-488 labeled), 3 µM profilin, 4 µM cofilin, 50 nM capping protein, 150 nM Arp2/3, 1 µM GST-pVCA-SspB-mCherry, 1 µM mCherry-SspB-mDia1. G-buffer: 2 mM Tris-HCl (pH 7.5), 0.2 mM ATP, 0.1 mM CaCl_2_, 0.5 mM DTT, 1 mM NaN_3_. KMEI-S buffer: .10mM imidazole (pH 7.0), 50mM KCl, 1mM MgCl_2_, 1mM EGTA, 240 mM sucrose (sucrose was added to mimic the buffer conditions in GUV experiments). (C–F) Pyrene actin polymerization assay. The reactions were performed with 2 µM actin (5% pyrene labeled) at RT. (C, D) 10 nM Arp2/3, and 100 nM NPFs. (-) represents actin and Arp2/3 only condition. (E) The reactions were performed with the indicated factors. (F) 50 nM capping protein, 150 nM Arp2/3, 3 µM profilin, 1 µM mCherry-SspB-mDia1, 1 µM GST-pVCA-SspB-mCherry. (-) represents actin only condition.

**Fig. S9.**
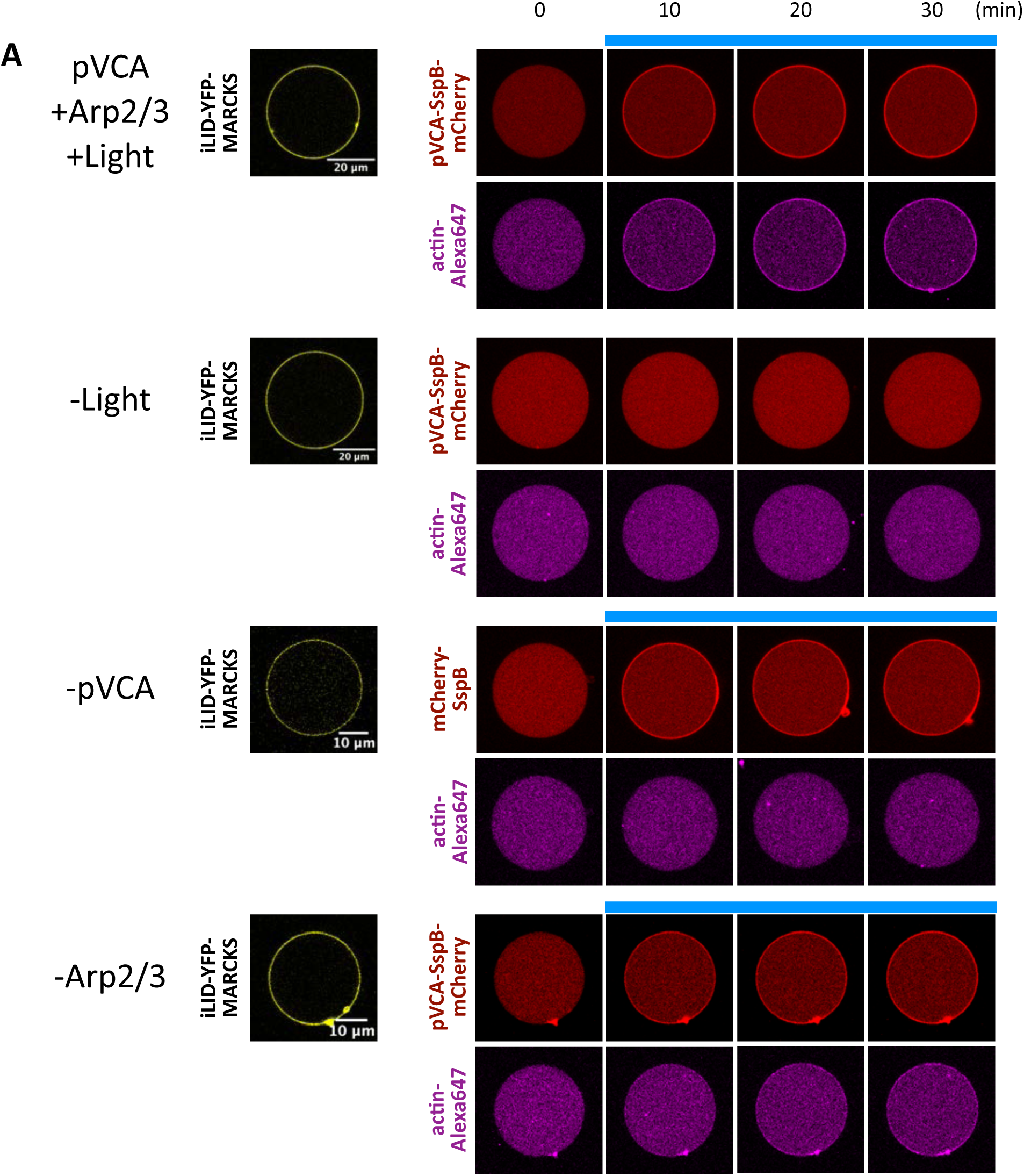
Light control of actin cytoskeleton with pVCA-SspB-mCherry. The data correspond to Fig. 2C–E. (+) condition contains 15 µM iLID-YFP-MARCKS, 4.2 µM pVCA-SspB(nano)-mCherry, 7.5 µM actin (10% Alexa 647 labeled), 150 nM Arp2/3, 100 nM capping protein, 3 µM profilin, 4 µM cofilin, and 1 mM ATP. In (-light) and (-Arp2/3) conditions, blue light illumination or Arp2/3 was omitted from (+pVCA +Arp2/3 +Light) condition, respectively. In (-pVCA) condition, pVCA-SspB(nano)-mCherry was replaced with mCherry-SspB(micro).

**Fig. S10.**
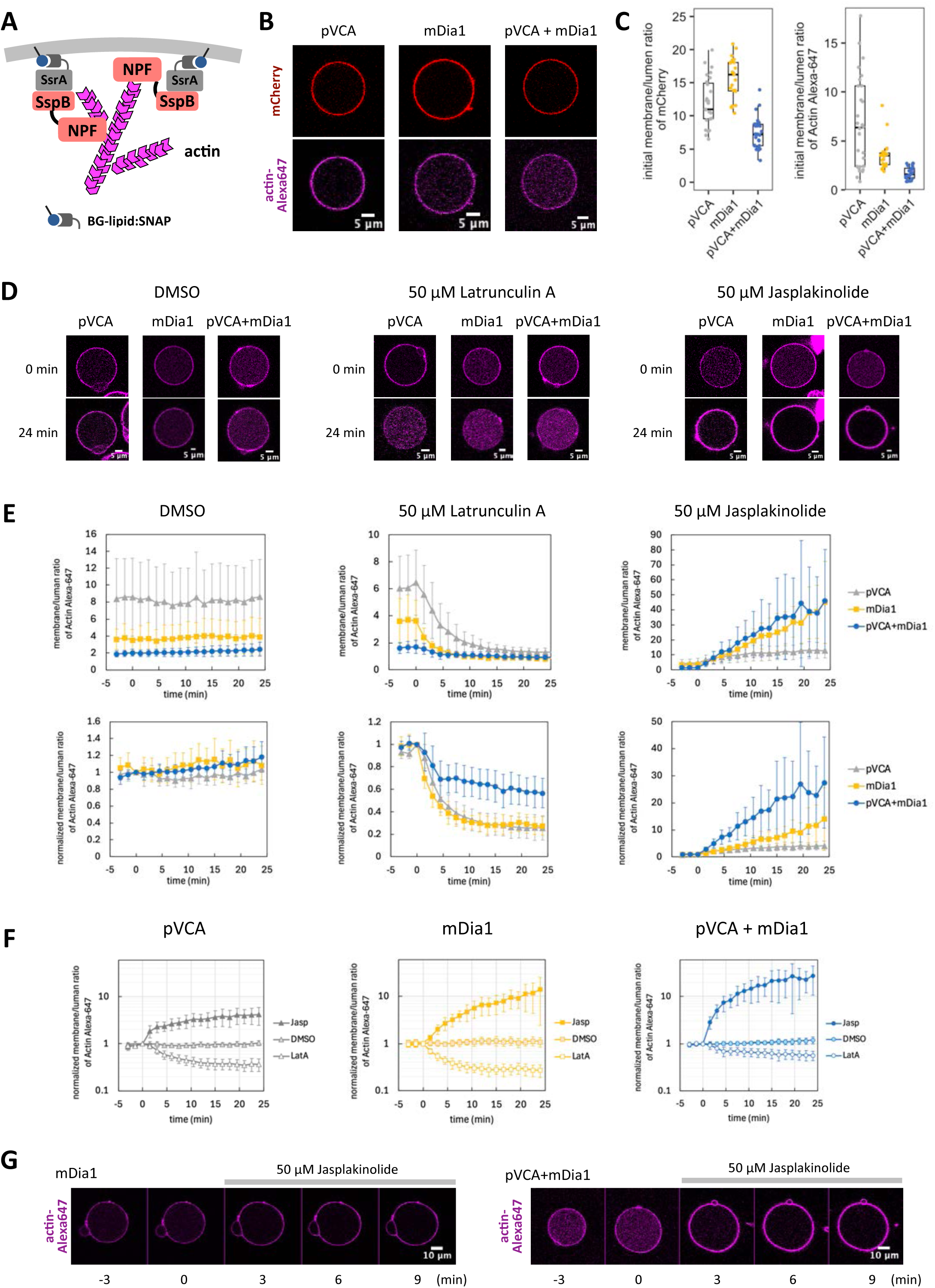
Pharmacological perturbation of membrane-associated actin assemblies. (A) Schematic representation of constitutive tethering of NPFs. SspB-fused NPFs (pVCA and/or mDia1) were tethered to the GUV membrane via the SsrA–SNAP/BG-lipid system. (B) Representative confocal images of membrane-associated actin assemblies formed by pVCA, mDia1, or their combination. Scale bars, 5 μm. (C) Quantification of the initial membrane/lumen fluorescence intensity ratio for mCherry and actin Alexa-647. pVCA: n=27 vesicles. mDia1: n=23 vesicles. pVCA + mDia1: n=27 vesicles. Box whisker plots represent median, 1st, 3rd quartiles and 1.5×inter-quartile range. (D) Representative images before (0 min) and after (24 min) treatment with DMSO, 50 μM Latrunculin A, or 50 μM Jasplakinolide. Scale bars, 5 μm. (E) Time courses of membrane/lumen ratio and normalized membrane/lumen ratio of actin Alexa-647. (F) Time courses of normalized membrane/lumen ratio of actin Alexa-647 for each NPF condition. The y-axis is shown on a logarithmic scale. (E, F) Error bars indicate 95% CI. (G) Representative images showing membrane deformation after Jasplakinolide treatment. Scale bars, 10 μm. mCherry-SspB-mDia1-6×His was used as mDia1 in this figure.

**Fig. S11.**
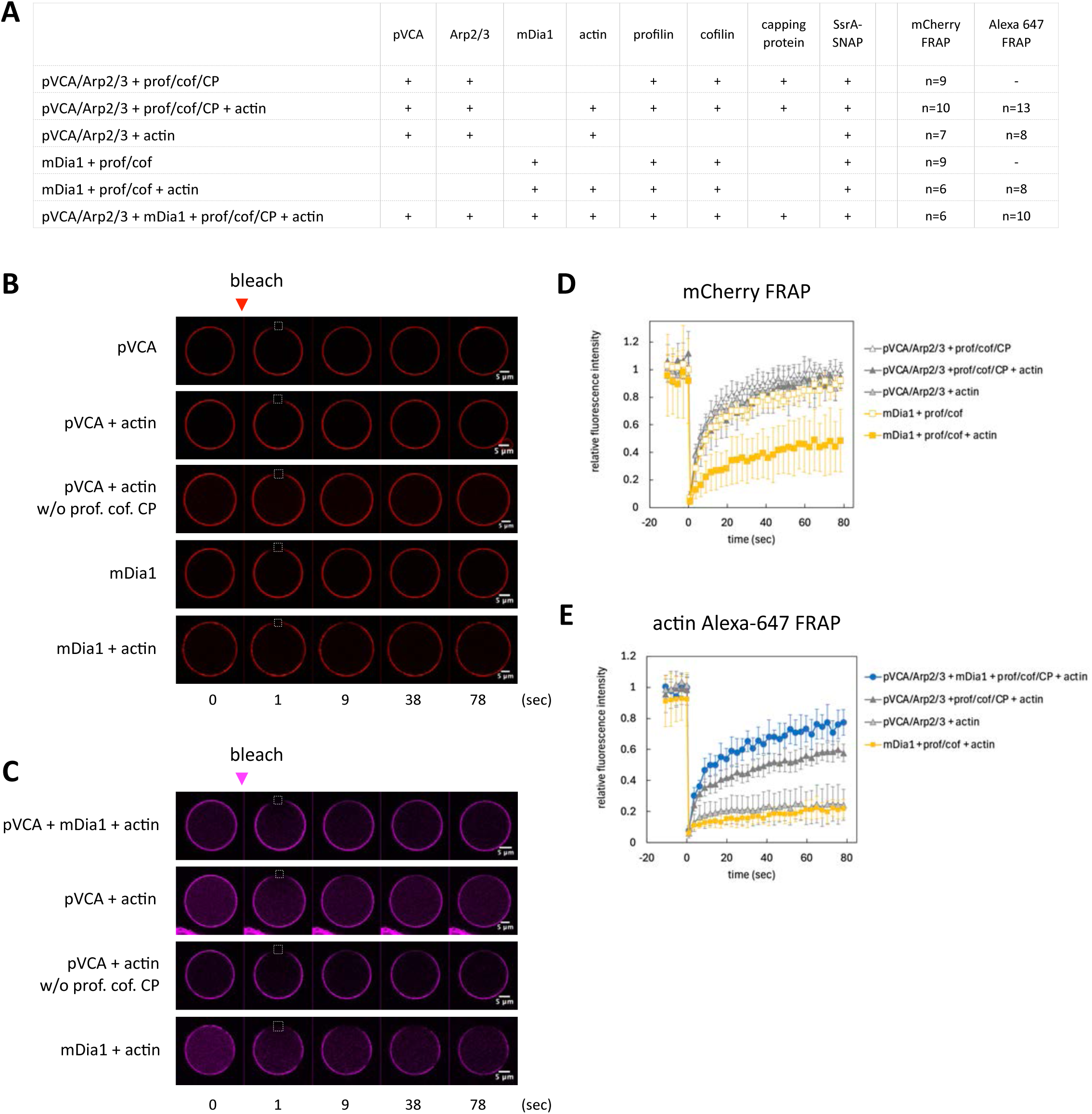
Fluorescence recovery after photobleaching (FRAP) analysis of membrane-associated NPFs and actin. (A) Summary table of proteins included in each conditions. In FRAP analysis, SspB-fused NPFs (pVCA and/or mDia1) were tethered to the GUV membrane via the SsrA–SNAP/BG-lipid system. (B, C) Representative confocal images of mCherry-tagged NPFs and actin Alexa-647. The white dashed box indicates the photobleached region. Scale bars, 5 μm. (D, E) Time course of mCherry and Alexa-647 fluorescence recovery at the bleached region. Fluorescence intensities were normalized to those of a non-bleached region to compensate for photobleaching during image acquisition. The data from Fig. 1G are reproduced to facilitate direct comparison with the other conditions presented in this figure.

**Fig. S12.**
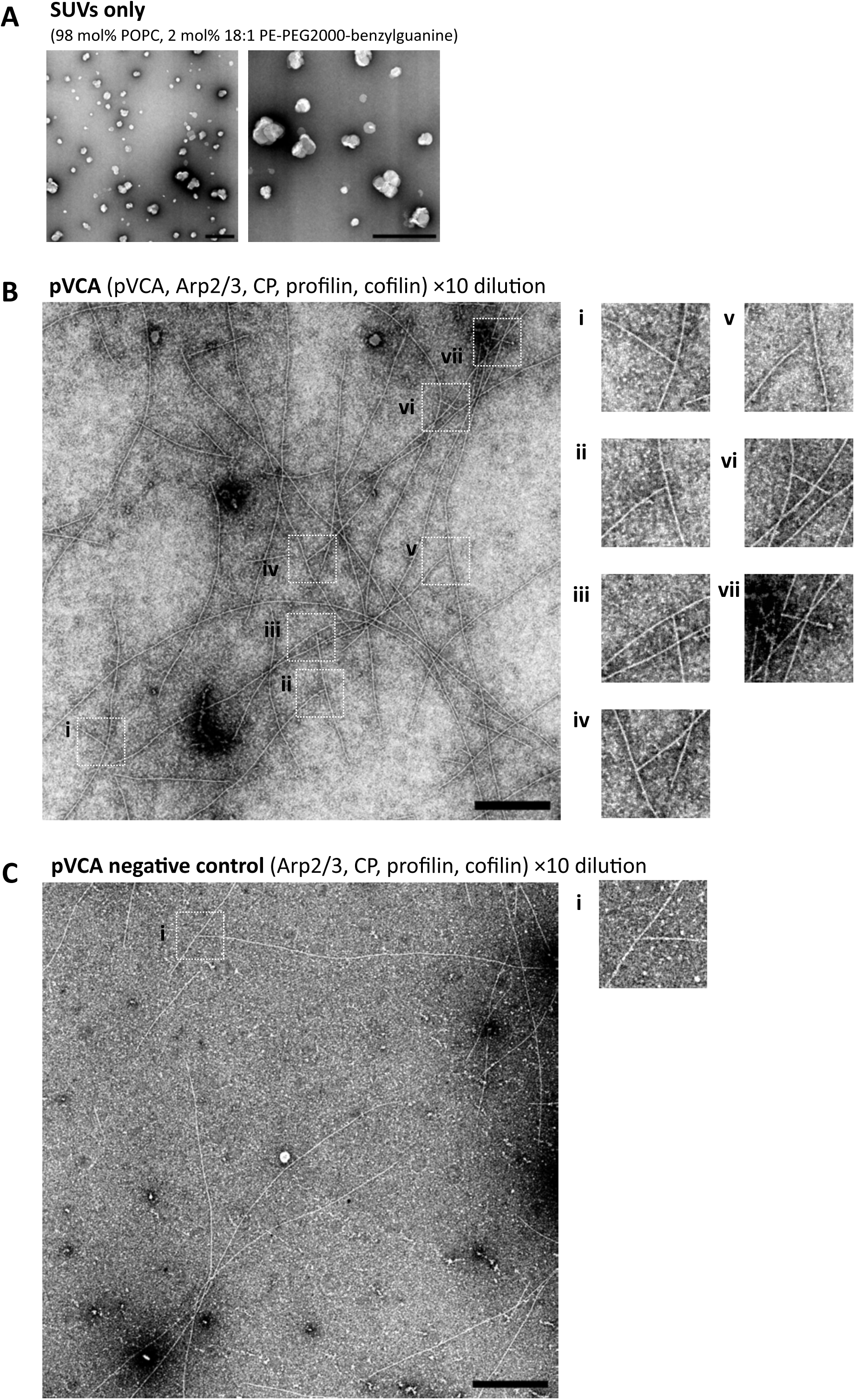
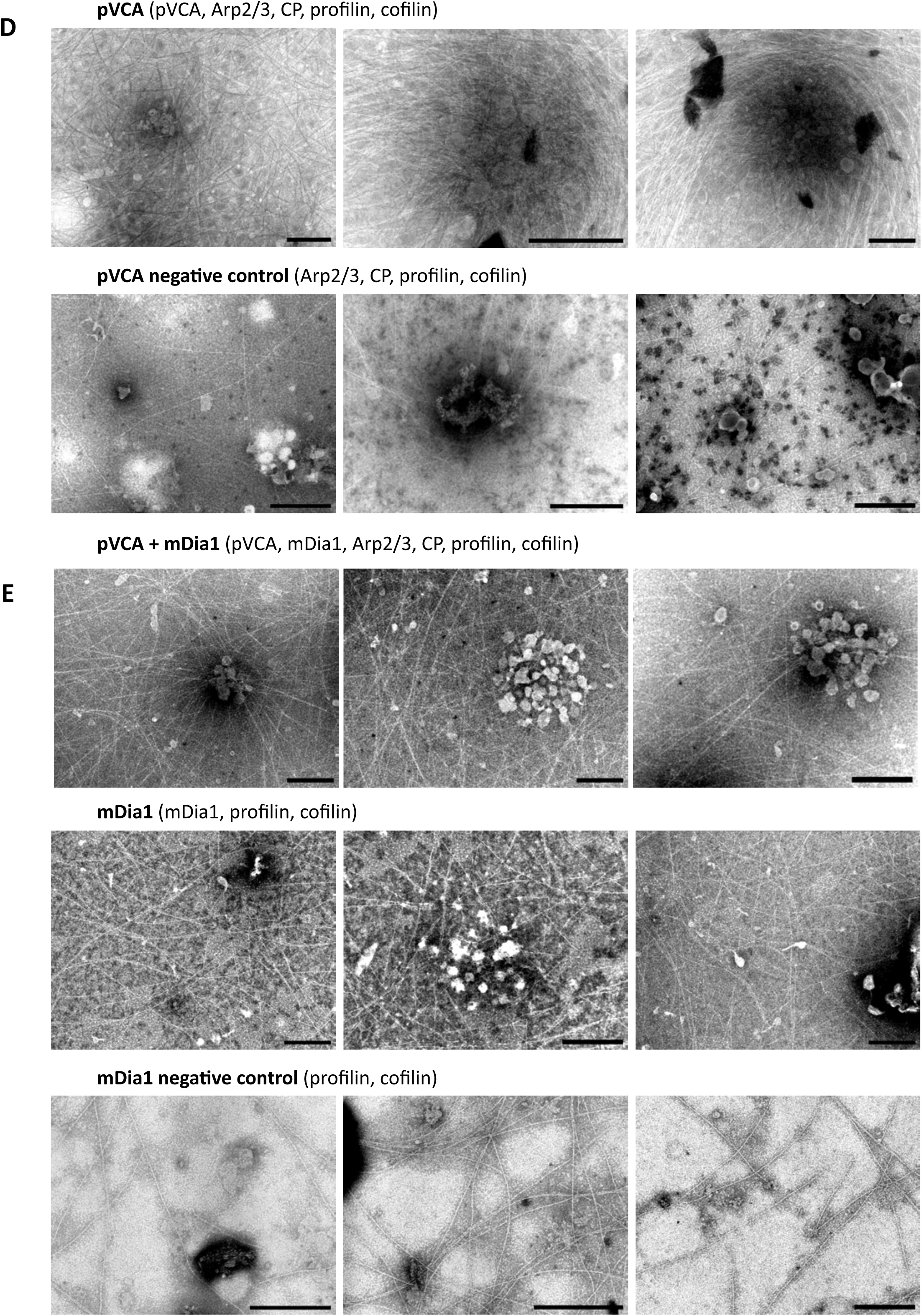
Electron microscopy (EM) analysis of actin filaments. Ultrastructure of small unilamellar vesicles (SUVs) and actin filaments were analyzed by negative staining electron microscopy. (A) Representative EM images of SUVs alone. (B) Representative images of Y-junctions formed in the presence of pVCA, Arp2/3, capping protein (CP), profilin, and cofilin. (C) Negative control sample lacking pVCA (Arp2/3, CP, profilin, and cofilin only). (B, C) To reduce filament overlap, the polymerization reactions were diluted 10-fold prior to deposition onto EM grids. (D) Representative EM images of actin assemblies formed with pVCA or the corresponding negative control lacking pVCA. (E) Representative EM images of actin assemblies formed with pVCA + mDia1, mDia1 alone, or the corresponding negative control lacking mDia1. All samples were deposited onto EM grids and negatively stained prior to imaging. Scale bars are shown in each panel. mCherry-SspB-mDia1-6×His was used as mDia1 in this figure.

**Fig. S13.**
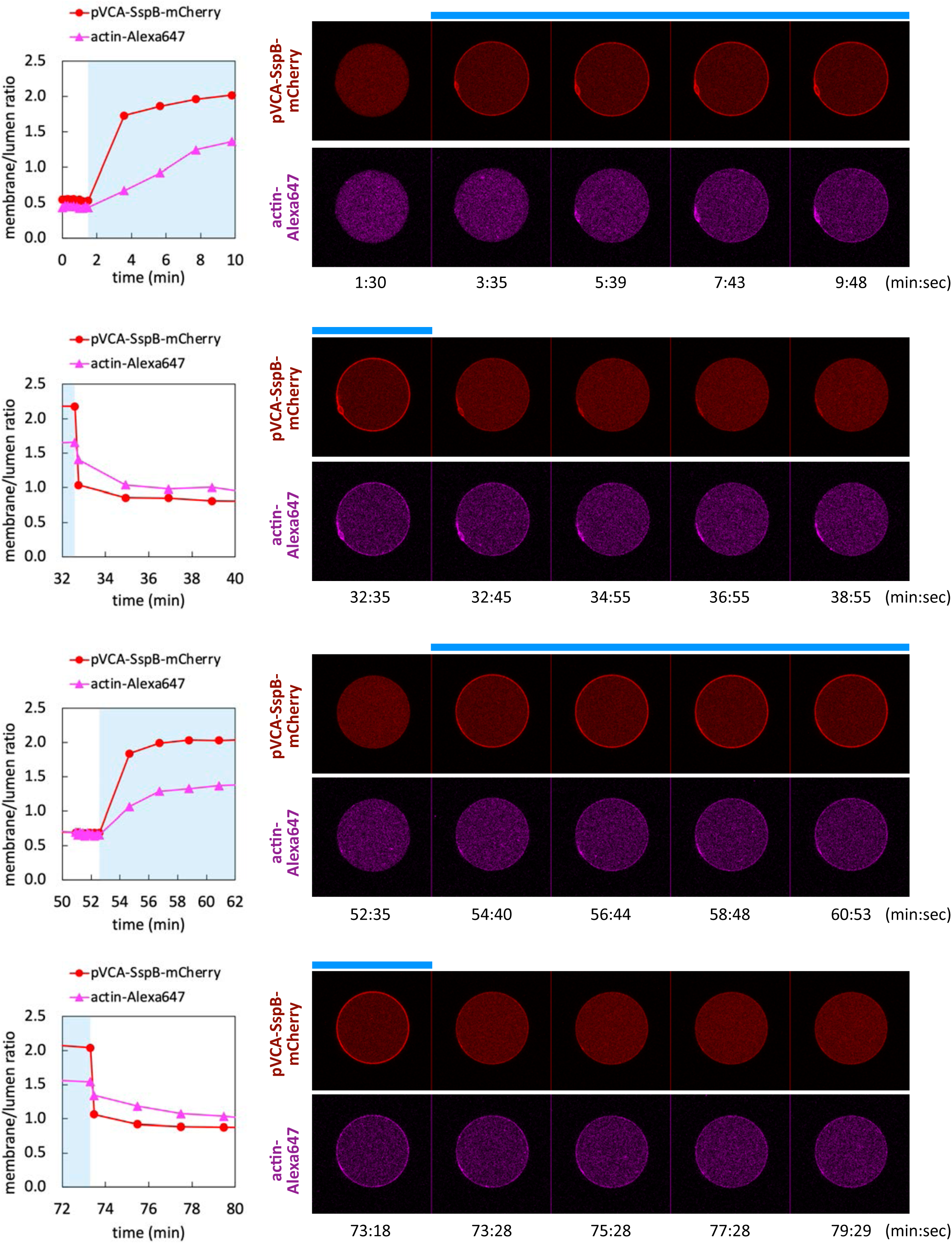
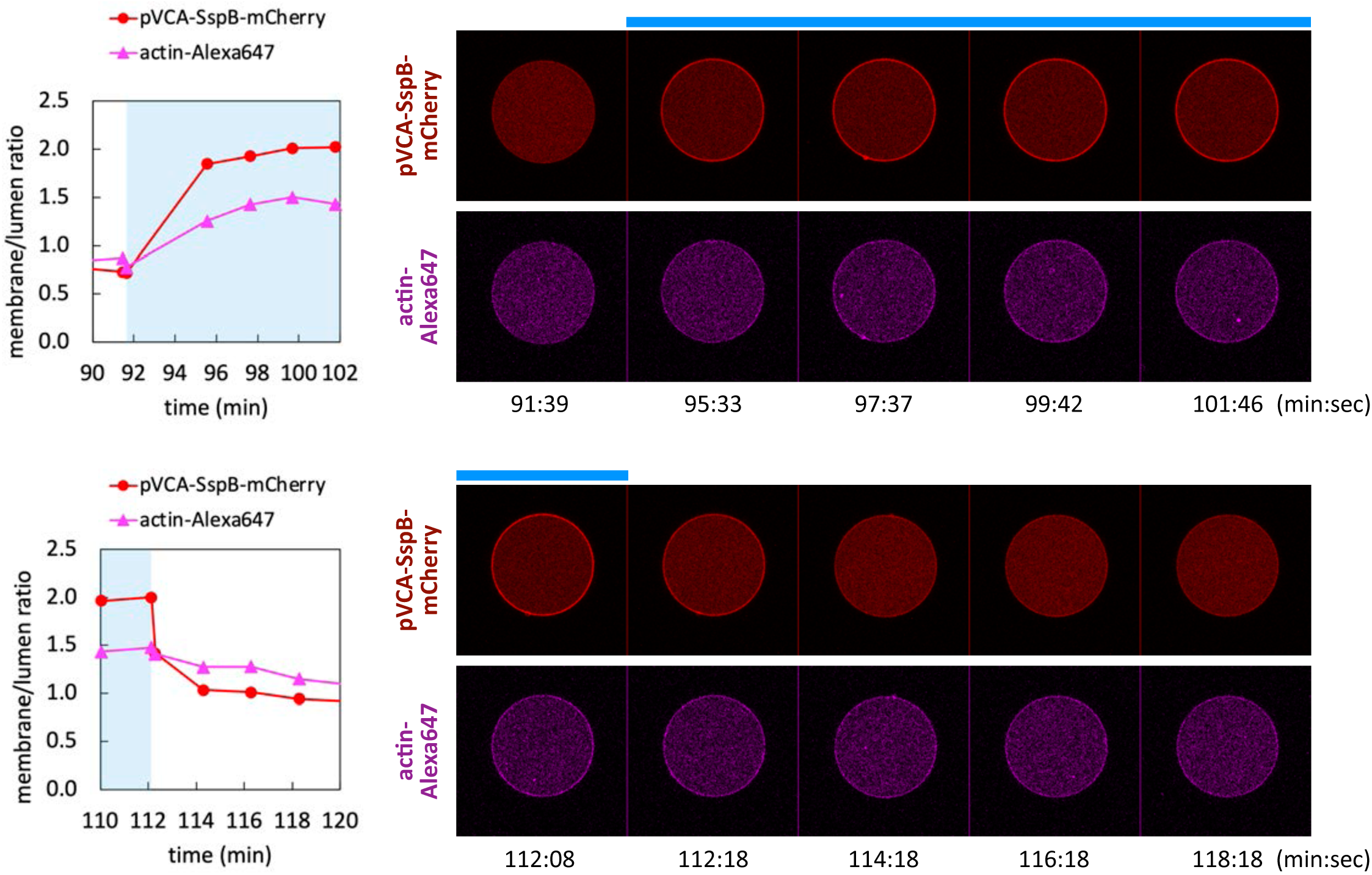
Representative images of reversible actin polymerization. Confocal images and time course of membrane/lumen ratio of pVCA-SspB-mCherry and Alexa 647-labeld actin. Blue bars on images and blue regions in graphs indicate timepoints with blue light illumination. This figure corresponds with Fig. 2F, G.

**Fig. S14.**
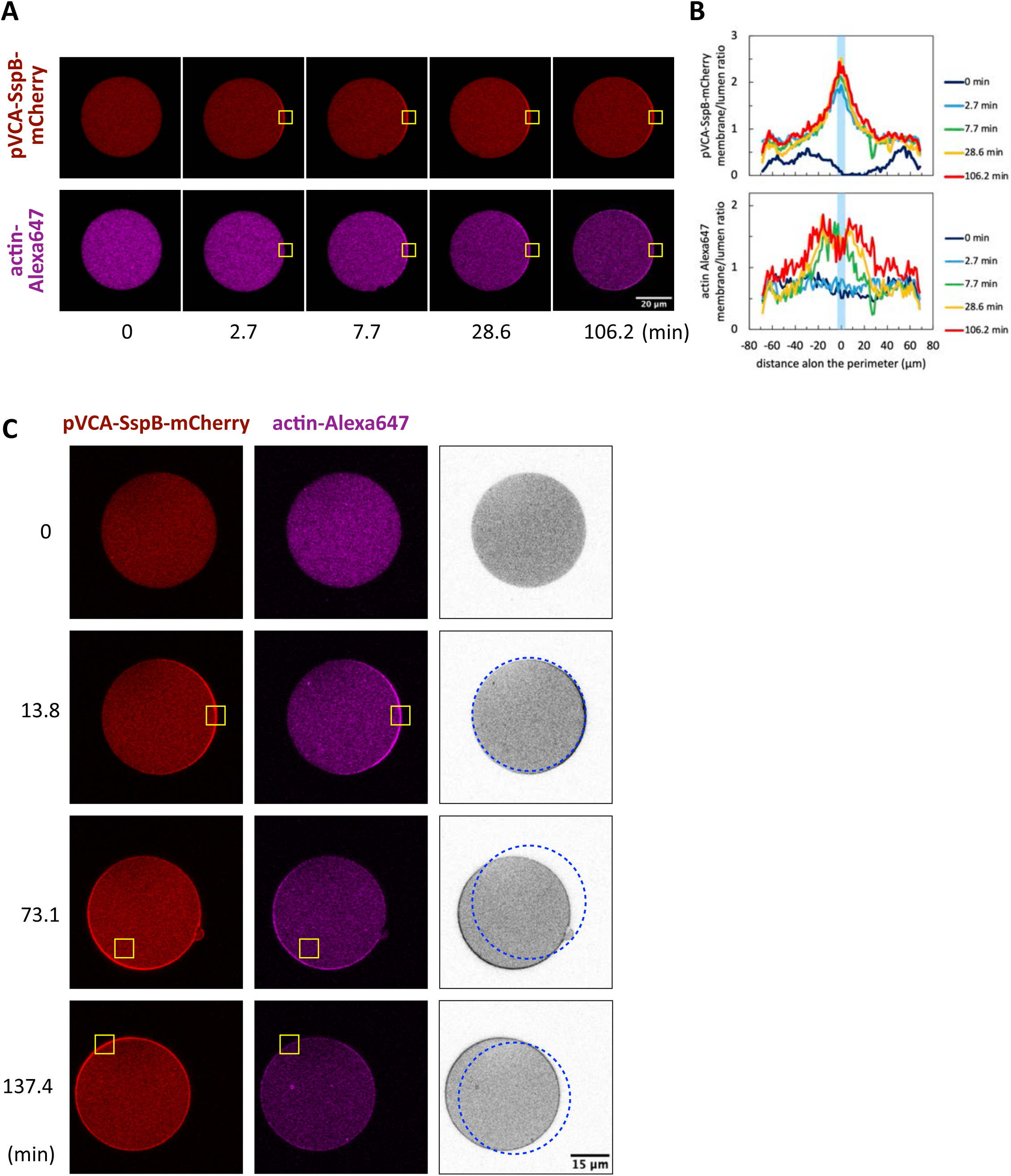
Asymmetric actin polymerization induced by pVCA. (A) Confocal images of local pVCA recruitment and actin polymerization. Yellow boxes indicate areas of blue light illumination. (B) Membrane/lumen ratio of mCherry-tagged pVCA and actin Alexa-647 for the images in (A). The distance is measured from the center of blue light illumination along the perimeter of the GUV. (A, B) The data corresponds to Fig. 3B and C. (C) Confocal images of local pVCA recruitment accompanied by vesicle movement toward light stimulation. Gray scaled images were created by averaging pVCA and actin images. Dotted circles indicate positions of vesicles in the images of one above.

**Fig. S15.**
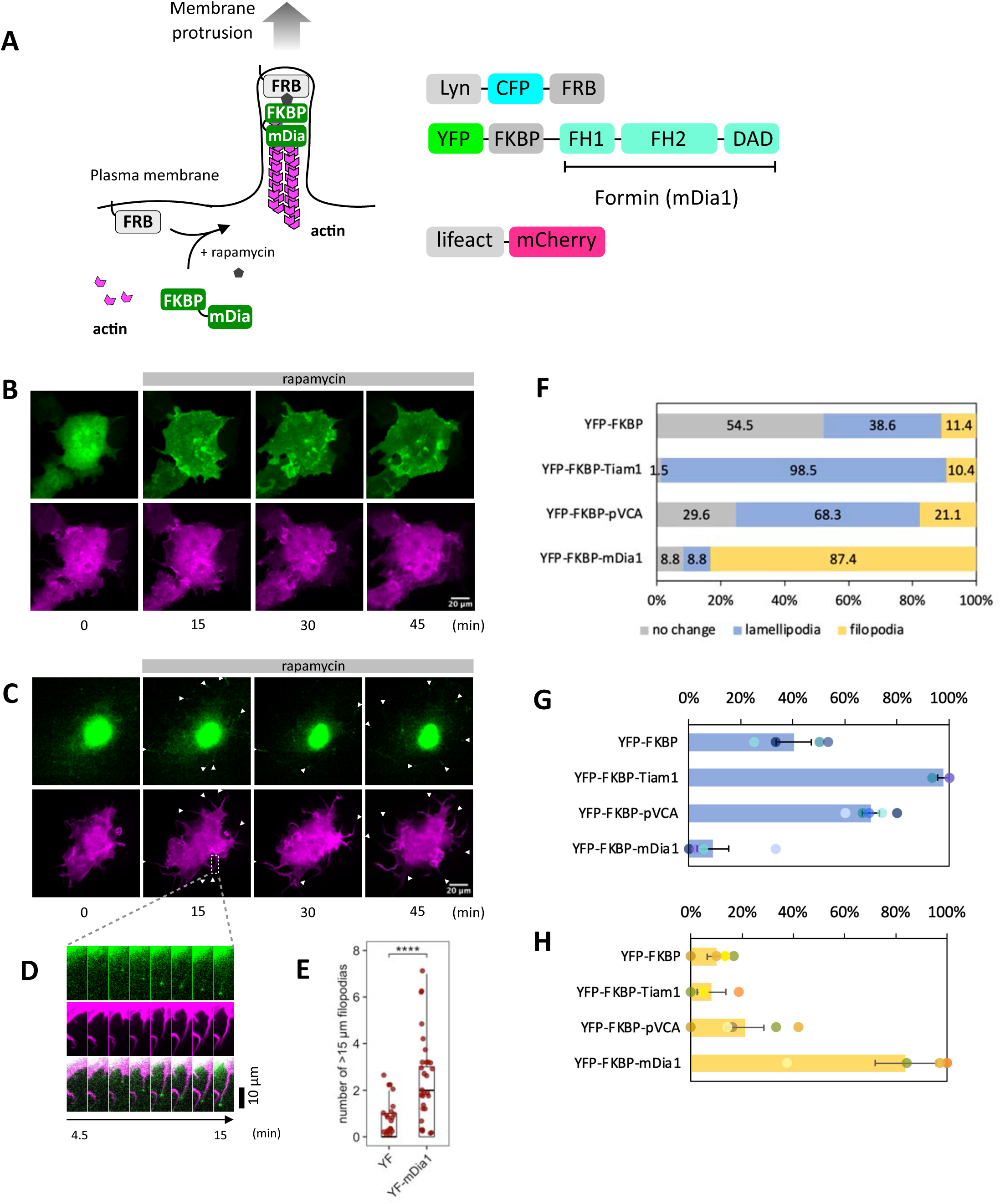
Quantitative analysis of mDia1-induced filopodia formation in Cos7 cells. (A) Schematic representation of chemically-inducible mDia1 translocation and membrane protrusions. (B, C) Confocal images of plasma membrane recruitment of mDia1 and subsequent filopodia formation. Cos7 cells were transfected with Lyn-ECFP-FRB, Lifeact-mCherry, and either of YFP-FKBP or YFP-FKBP-mDia1. At time 0, 100 nM rapamycin was added. Green: YFP-FKBP or YFP-FKBP-mDia1. Magenta: Lifeact-mCherry. (B) YFP-FKBP control. (C) YFP-FKBP-mDia1. (D) Zoomed-up view of filopodia extension. Images were acquired every 1.5 minutes. The bottom is merged image. (E) Quantification of mDia1-induced filopodia. Number of filopodia whose length is longer than 15 µm was counted. Box whisker plots represent median, 1st, 3rd quartiles and 1.5×inter-quartile range. P-values: ****: < 0.0001. Wilcoxon rank sum test. YF (YFP-FKBP): n=36 cells. YF-mDia1 (YFP-FKBP-mDia1): n=33 cells. (F–H) Quantification of the phenotypes observed after plasma membrane translocation of mDia1 (n=159 cells), pVCA (n=142 cells), Tiam1 (Rac1 GEF) (n=67 cells), and control protein YFP-FKBP (n=44 cells).

**Fig. S16.**
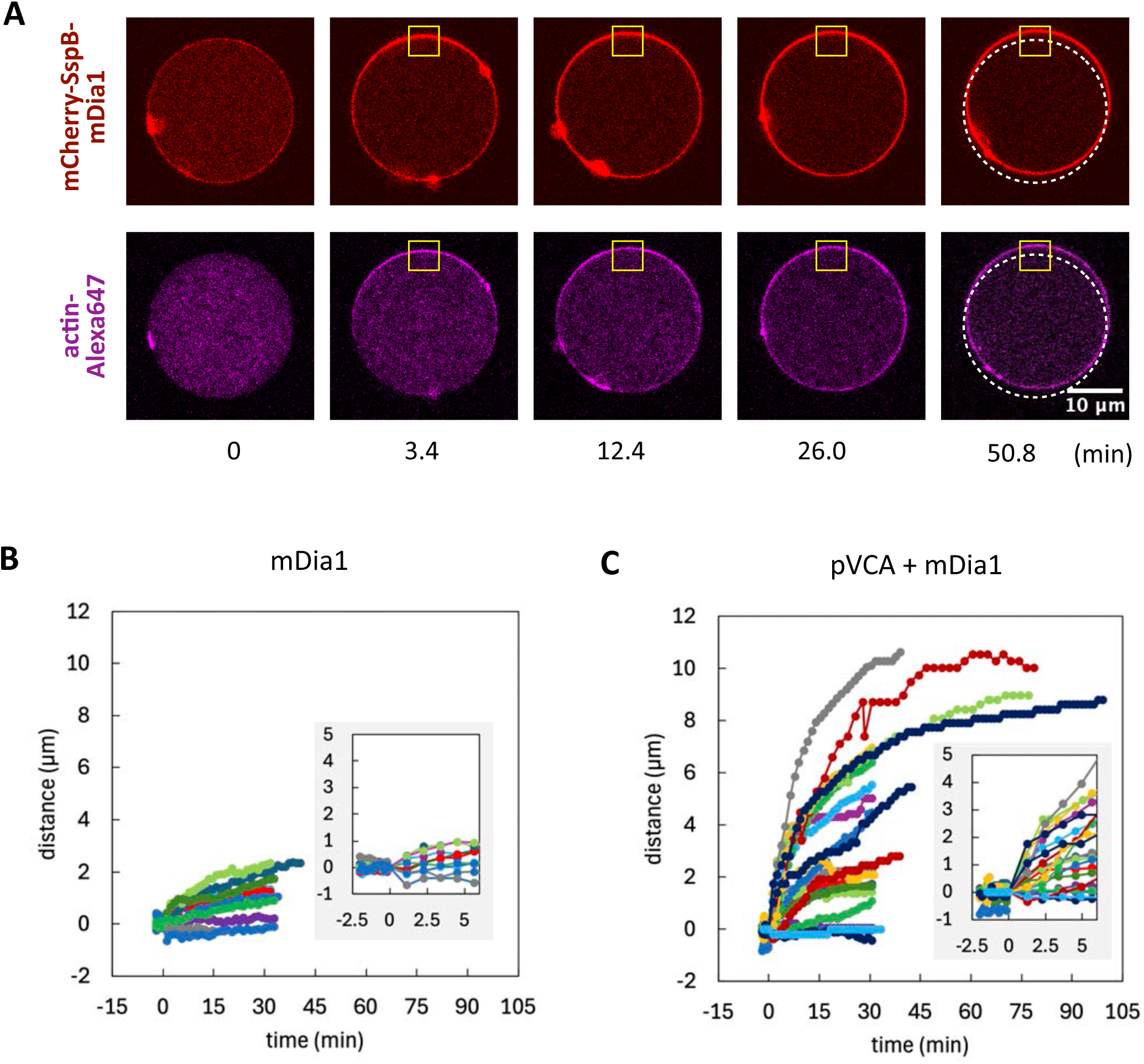
mDia1-mediated GUV movement and the comparison between mDia1 and pVCA + mDia1 system. (A) Representative images of mDia1-mediated GUV movement. Yellow boxes indicate the area of blue light stimulation. Dotted lines indicate the initial position of the GUV. (B, C) Time course of the distance the front (illuminated) side of GUV membrane moved forward. Each line represents each GUV. Insets show the initial response.

**Fig. S17.**
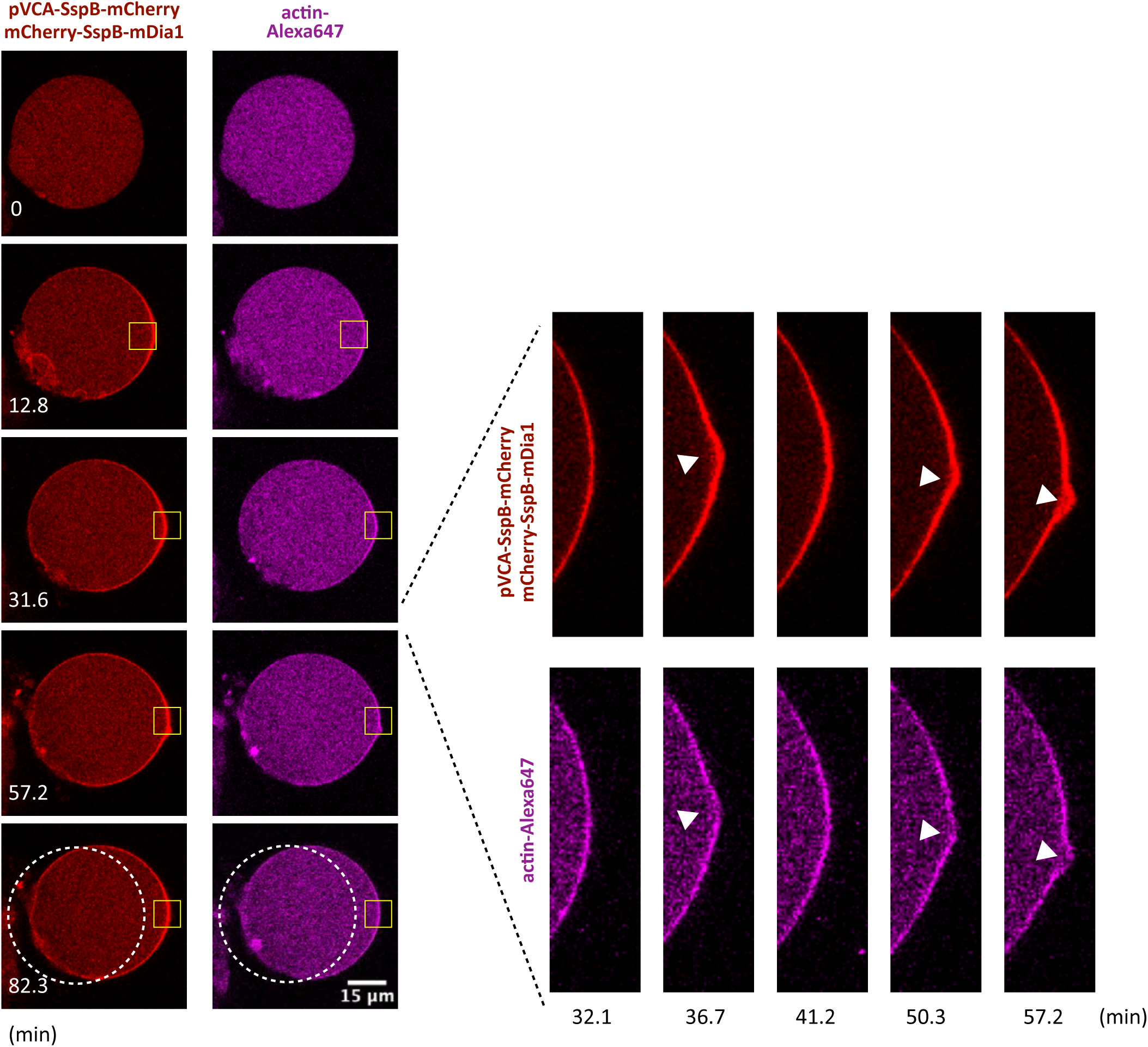
Local protrusions in pVCA + mDia1 GUVs. Representative images of local protrusions seen in pVCA + mDia1 GUVs. Yellow boxes indicate the area of blue light stimulation. Dotted lines indicate the initial position of the GUV. White arrows indicate where local protrusions arose.

**Fig. S18.**
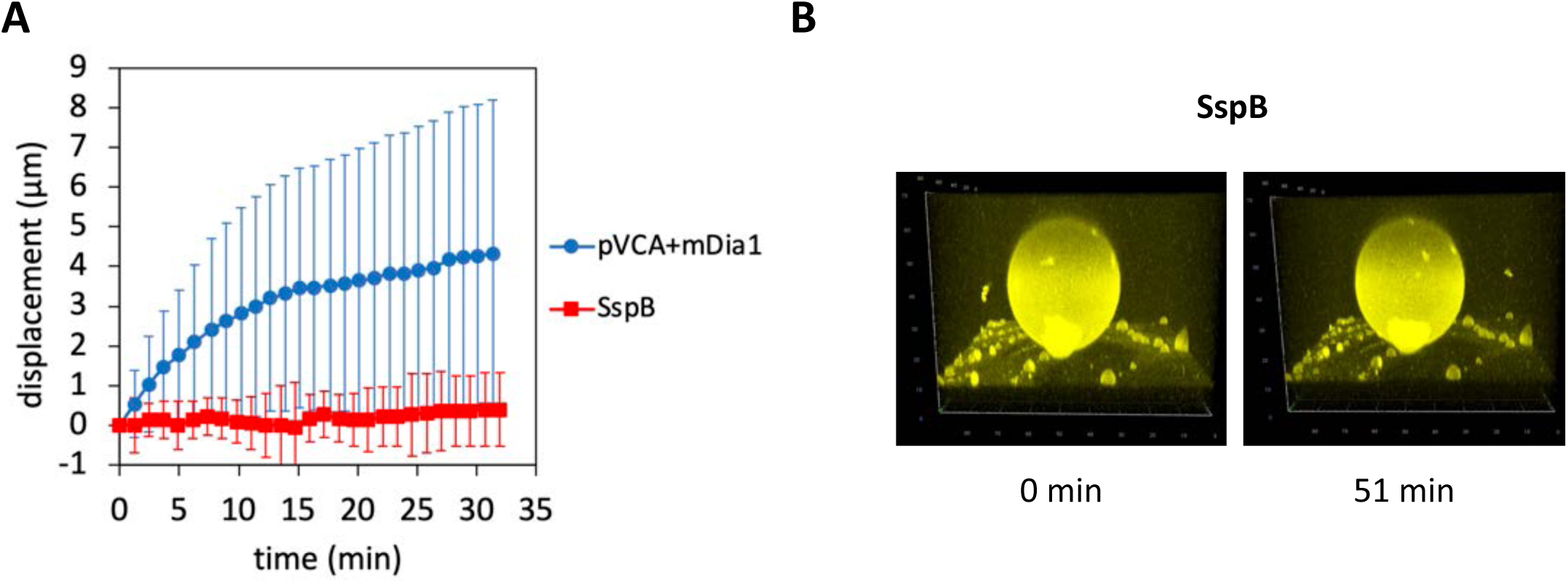
Lack of vesicle movement and shape change with mCherry-SspB. With pVCA-SspB-mCherry and mCherry-SspB-mDia1 being replaced with mCherry-SspB, GUVs neither moved nor changed their shape after blue light illumination. (A) Time course of the distance the front (illuminated) side of GUV membrane moved forward. pVCA + mDia1 GUVs: n=6 vesicles. SspB GUVs: n=4 vesicles. Error bars indicate 95% CI. (B) 3D reconstitution of iLID-YFP-SNAP images before and after blue light illumination.

**Fig. S19.**
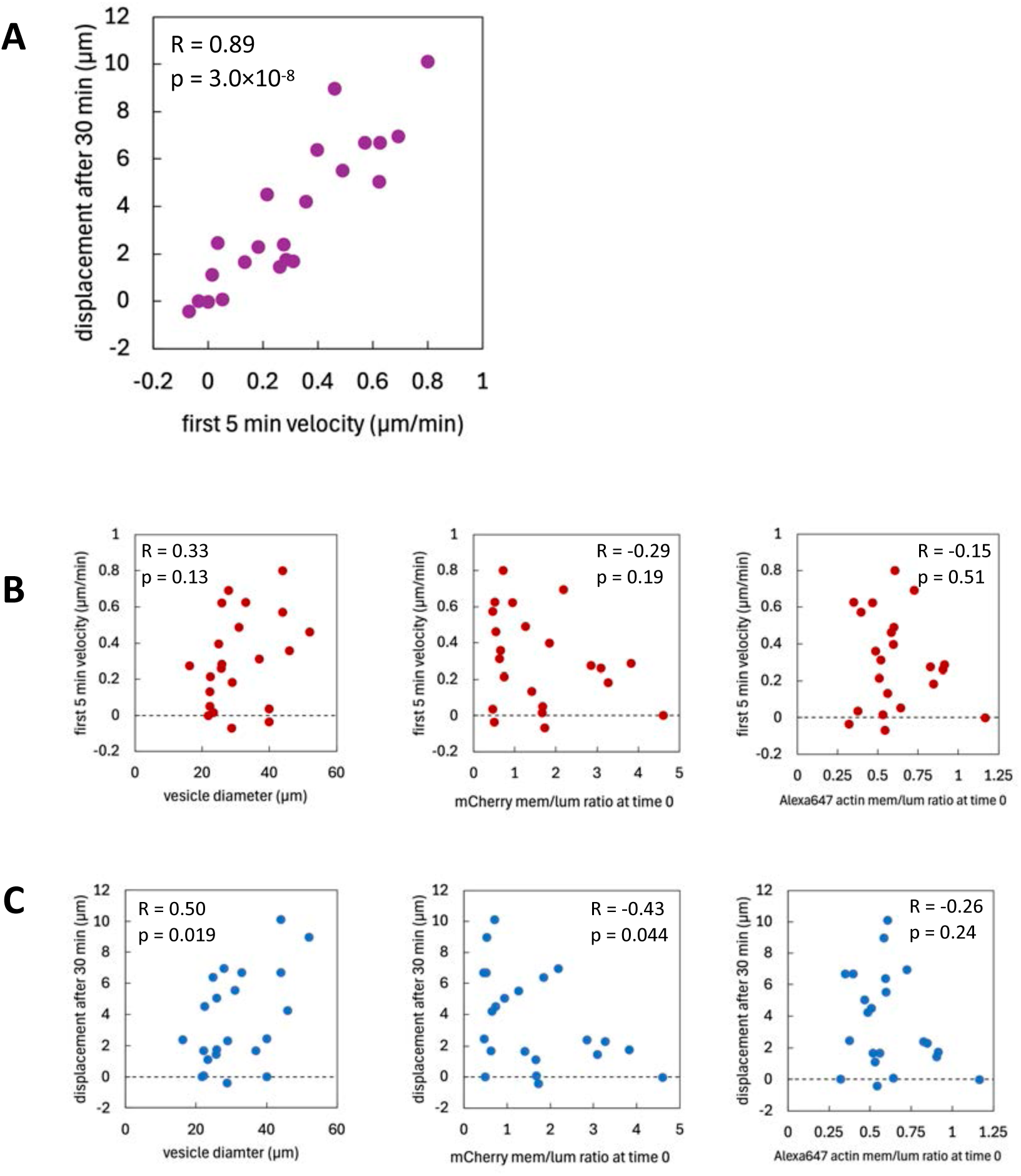
Relationship between vesicle motion and biophysical parameters. (A) Relationship between initial velocity (average velocity over the first 5 minutes) and net displacement after 30 minutes of GUV forward movement. (B, C) Biophysical parameters are plotted against initial velocity and displacement are plotted in (B) and (C), respectively. R and p indicate Pearson’s correlation coefficient and its p-value.

**Fig. S20.**
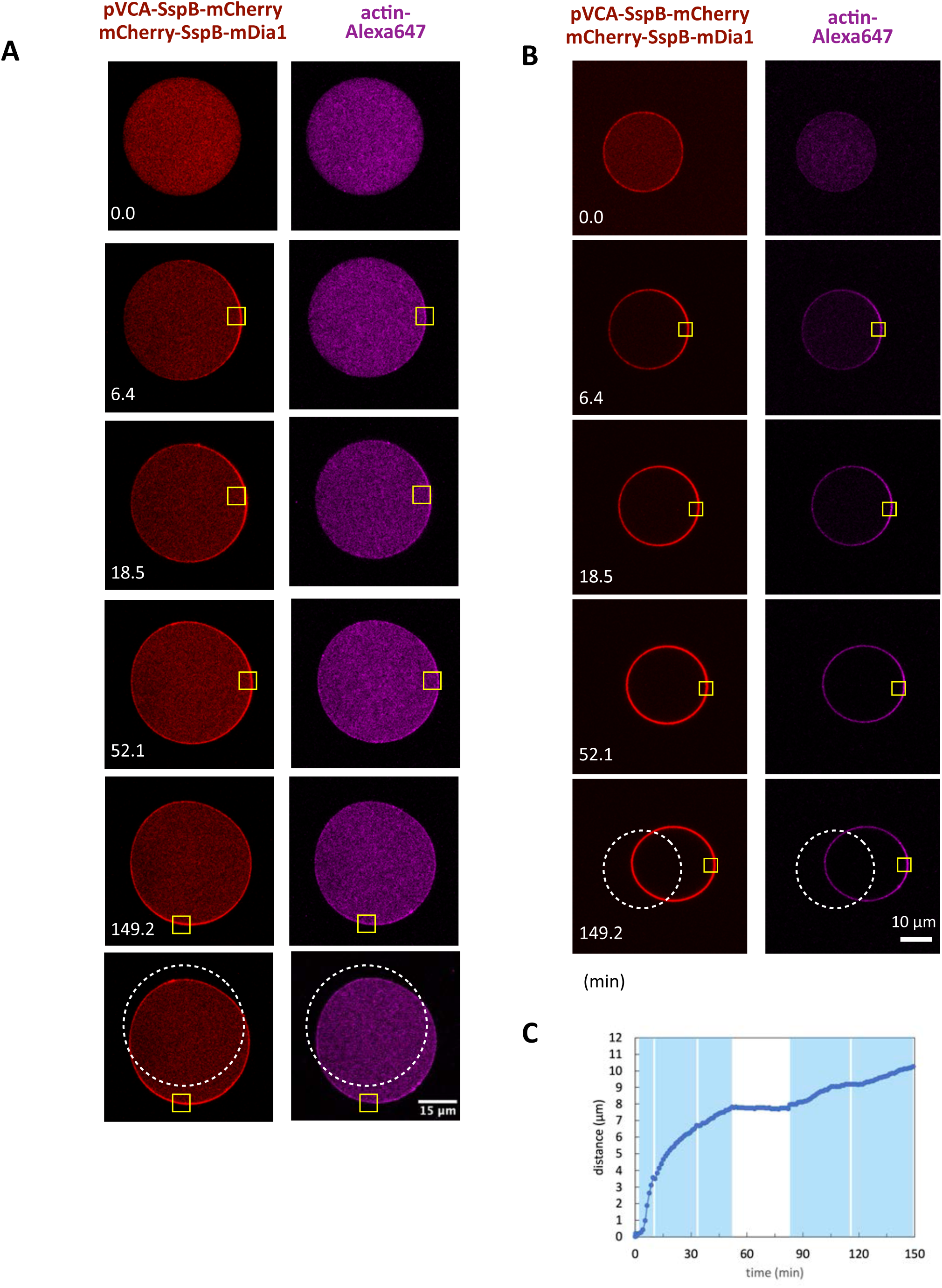
Representative images of pVCA-mDia1-mediated movement of GUVs. (A, B) Representative images of pVCA-mDia1-mediated movement of GUVs. Yellow boxes indicate the area of blue light stimulation. Dotted lines indicate the initial position of the GUV. White arrows indicate where local protrusions arose. (C) Distance the front side of GUV membrane moved forward. Data was quantified from the images of (B). Blue areas indicate the periods of blue light stimulation.

**Fig. S21.**
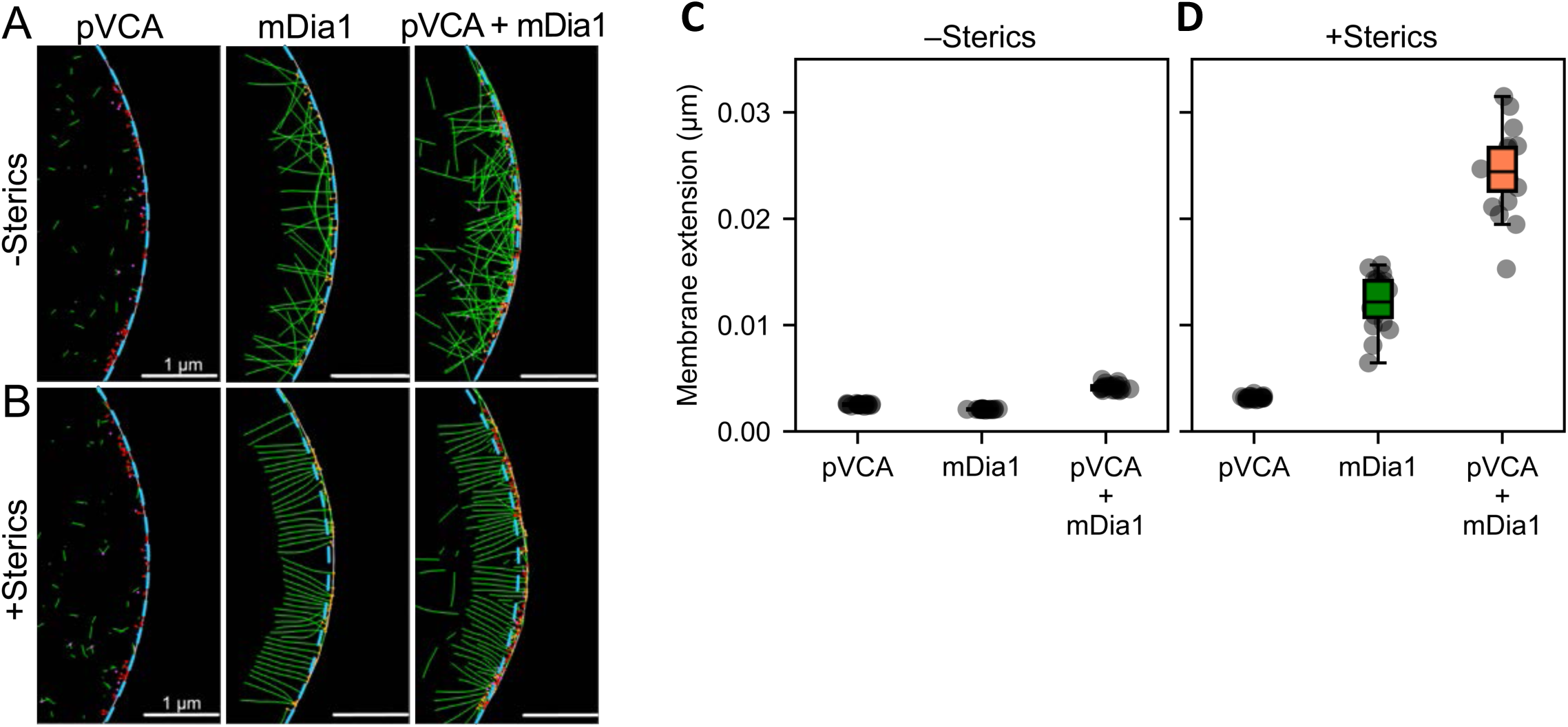
Computational modeling of actin network dynamics underlying membrane deformation. (A, B) Snapshots showing actin filaments at the leading edge of the membrane after 5 s of simulation. Actin filaments are shown in green, pVCA-Arp2/3 is shown in red, mDia1 is shown in orange, and Arp2/3 is shown in magenta. The membrane location at t = 0 is shown with the cyan dotted line to highlight the extent of membrane deformation. Simulations without and with inter-filament steric interactions are shown in A and B, respectively. (C, D) Box-whisker plots showing distributions of the membrane extension for the different actin-binding proteins. Results for networks without and with inter-filament steric interactions are shown in A and B, respectively. Each box plot is constructed using 20 independent simulation trajectories. Error bars represent standard deviation about the mean.

**Movie S1** Representative movie of repetitive and reversible SspB translocation. The data corresponds to Fig. 1G. Time in min:sec. Yellow regions indicate the area of blue light stimulations.

**Movie S2** A rare example where ActA induced actin polymerization in the direction of light stimulus. The data corresponds the leftmost images in Fig. S6. Red: ActA-SspB(nano)-mCherry. Magenta: Alexa 647-labeled actin. Time in hr:min:sec. Yellow regions indicate the area of blue light stimulation.

**Movie S3** Representative movie of global membrane recruitment of pVCA-SspB-mCherry. The data corresponds to Fig. 2C and “pVCA +Arp2/3 +Light” in Fig. S9. Red: pVCA-SspB-mCherry. Magenta: Alexa 647-labeled actin. Time in hr:min:sec. The label “Light” indicates when blue light stimulation is applied to the entire GUV.

**Movie S4** Representative movie of global and reversible membrane recruitment of pVCA-SspB-mCherry. The data corresponds to Fig. 2I and S13. Red: pVCA-SspB-mCherry. Magenta: Alexa 647-labeled actin. Time in hr:min:sec. The label “Light” indicates when blue light stimulation is applied to the entire GUV.

**Movie S5** Representative movie of asymmetric membrane recruitment of pVCA-SspB-mCherry. The data corresponds to Fig. 3B and C. Red: pVCA-SspB-mCherry. Magenta: Alexa 647-labeled actin. Time in hr:min:sec. Yellow regions indicate the area of blue light stimulation.

**Movie S6** Representative movie of asymmetric and reversible membrane recruitment of pVCA-SspB-mCherry. The data corresponds to Fig. 3D and E. Red: pVCA-SspB-mCherry. Magenta: Alexa 647-labeled actin. Time in hr:min:sec. Yellow regions indicate the area of blue light stimulation.

**Movie S7** Representative movie of pVCA-mDia1-mediated movement of GUV. The data corresponds to Fig. 4B and E. Red: pVCA-SspB-mCherry and mCherry-SspB-mDia1. Magenta: Alexa 647-labeled actin. Time in hr:min:sec. Yellow regions indicate the area of blue light stimulation.

**Movie S8** Representative movie of local protrusions seen in pVCA + mDia1 GUVs. The data corresponds to Fig. S17. Red: pVCA-SspB-mCherry and mCherry-SspB-mDia1. Magenta: Alexa 647-labeled actin. Time in hr:min:sec. Yellow regions indicate the area of blue light stimulation. In the zoomed-in view, the relative movement of the GUV was subtracted to better highlight the local protrusions.

**Movie S9** Representative movie of pVCA-mDia1-mediated movement of GUV. The data corresponds to Fig. 4F and S20A. Red: pVCA-SspB-mCherry and mCherry-SspB-mDia1. Magenta: Alexa 647-labeled actin. Time in hr:min:sec. Yellow regions indicate the area of blue light stimulation.

**Movie S10** Representative movie of pVCA-mDia1-mediated movement of GUV. The data corresponds to Fig. S20B. Red: pVCA-SspB-mCherry and mCherry-SspB-mDia1. Magenta: Alexa 647-labeled actin. Time in hr:min:sec. Yellow regions indicate the area of blue light stimulation.

**Movie S11** Representative simulation movies in the absence of inter-filament steric interactions. Actin filaments are shown in green, pVCA–Arp2/3 in red, mDia1 in orange, and Arp2/3 in magenta. The membrane position at *t* = 0 is marked by a cyan line to indicate the extent of membrane deformation.

**Movie S12** Representative simulation movies in the presence of inter-filament steric interactions. Actin filaments are shown in green, pVCA–Arp2/3 in red, mDia1 in orange, and Arp2/3 in magenta. The membrane position at *t* = 0 is marked by a cyan line to indicate the extent of membrane deformation.

