## Supplementary Texts for "Light-guided actin polymerization drives directed motility in protocells"

**Supplementary Text 1: Locality of iLID-SspB system**

The locality of SspB recruitment is determined by how fast and how far the iLID-SspB complex can diffuse in the unilluminated (dark) region before dissociating. Since this diffusion occurs on a two-dimensional lipid membrane, we assumed the locality would follow the equation:

$$\langle x^2 \rangle = 4Dt$$

where D is the diffusion coefficient and t is the half-time of the iLID recovery reaction in the dark. Using a typical diffusion coefficient of D = 6.0 $\mu m^2/s$ for a POPC membrane (*1*, *2*) and a half-time t = 42 s from our experiments (Fig. S3A), we calculated $\langle x^2 \rangle = 1.0\times10^3\ \mu m^2$. To ensure that this diffusion area is less than half the surface area of a sphere, the radius of the giant vesicle must be greater than 12.6 µm. In other words, the iLID-SspB system can establish sufficient polarity in a vesicle with a diameter larger than 25.2 µm.

**Supplementary Text 2: Optimization for iLID membrane anchoring**

To anchor iLID to the GUV membrane, we tested four protein-lipid interaction strategies, namely 6×His-tag/Ni-NTA-conjugated lipids, myristoylated alanine-rich C-kinase substrate-effector domain (MARCKS-ED)/phosphatidylserine (PS), 2×Strep-tag/StrepTactin/biotin-conjugated lipids, and SNAP-tag/benzylguanine (BG)-conjugated lipid. As commonly used in many reconstitution studies, we first investigated the 6×His-tag/Ni-NTA-conjugated system. However, the His-tag Ni-NTA pair caused non-specific membrane recruitment of SspB (Fig. S2A, B). While the exact reason remains unclear, this is possibly caused by the presence of 2×Strep-iLID-YFP-6×His because mCherry-SspB itself did not show non-specific binding to the membrane of 5% Ni-NTA DGS/95% POPC (Fig. S2C). We attempted to prevent this non-specific binding by adding PEGylated lipid up to 2%, but this resulted in reduced membrane anchoring of iLID rather than suppression of non-specific SspB binding (Fig. S2B). Next, we examined the MARCKS-ED/PS combination. Reflecting the weaker interaction (µM range Kd) of MARCKS-ED/PS compared to 6×His-tag/Ni-NTA (nM range Kd) (*3*, *4*), PS content had to be increased to 40% to effectively anchor iLID to the membrane (Fig. S2D, E). At 40% PS, we occasionally observed non-specific membrane binding of SspB, but this non-specific binding was mitigated by adding 0.5-2% PEGylated lipid (Fig. S2F). However, lipid compositions containing PS presented less reproducibility in GUV preparation. We also investigated the method of bridging biotin-modified lipids and Strep-Tag-II using StrepTactin, but determining the optimal stoichiometry of these three components proved challenging (Fig. S2G, H). The SNAP-tag/BG-conjugated lipid combination allowed sufficient membrane anchoring of iLID with just 2% BG-conjugated lipid and showed no non-specific SspB binding (Fig. S2I, J). Furthermore, BG-conjugated lipid provided high reproducibility in GUV preparation; therefore, we primarily used SNAP/BG for most of the experiments.

**Supplementary Text 3: Optimization of ActA–SspB and limitations of ActA-mediated asymmetric actin polymerization**

To optimize ActA-SspB for membrane recruitment and actin polymerization activity, we tested two SspB variants with different affinities and two fusion orders (resulting in four ActA-SspB constructs in total). Among three available SspB variants with different affinities for iLID under blue light—SspB(nano) (Kd = 0.13 μM), SspB(micro) (Kd = 0.80 μM), and SspB(milli) (Kd = 56 μM)—we used SspB(nano) and SspB(micro), whose Kd values fall within the concentration range used in this study (*5*, *6*). In bulk pyrene actin assays, which monitors actin polymerization through the florescence increase of pyrene-labeled actin, the fusion order ActA-SspB-mCherry exhibited relatively higher actin polymerization activity than the corresponding ActA-mCherry-SspB ones, suggesting possible steric hindrance between ActA and mCherry (fig. S4A, B). When tested for light-dependent membrane recruitment in GUV, the four constructs showed graded affinity differences. Although ActA-mCherry-SspB(nano) showed the highest membrane recruitment, its light-independent membrane binding was considerable (fig. S4C–E). Based on these results, we adopted ActA(1-183)-SspB(nano)-mCherry for combination with the actin system in GUVs.

In figure S6, light-induced membrane recruitment of ActA generated actin polymerization on the GUV membrane but failed to maintain stable directional asymmetry, as polymerized actin rapidly diffused away from the illuminated region. We reasoned that insufficient actin turnover at the membrane might limit sustained asymmetric actin organization. To facilitate the turnover, we attempted to induce membrane-associated actin polymerization in the presence of cofilin and capping proteins. However, under these conditions, ActA did not robustly induce actin polymerization on the GUV membrane (fig. S7). These results suggest that ActA alone is insufficient to support persistent, asymmetric actin remodeling in this optogenetic GUV system, motivating the exploration of more potent alternative NPFs.

**Supplementary Text 4: Computational modeling of actin network dynamics underlying membrane deformation**

An outstanding question is how growth of the actin network in the GUV system can deform the membrane—leading to motility? In cells, there are adhesion molecules such as adhesin and integrin which anchor the cell's membrane and cytoskeleton to the substrate (*7*, *8*). In such anchored systems, actin polymerization at the leading edge causes retrograde flow of filaments which are anchored at these adhesion sites enabling traction (*7*). In our reconstituted GUV system, there is no such backing mechanism which begs the question for why the growing actin network does not undergo unrestrained retrograde flow.

We hypothesized that there could be two main mechanisms by which bracing can be achieved considering the minimal components of the engineered GUV system. First, the network may be distributing load *through* the network configuration and to the membrane through the iLiD-SspB dimers which tether mDia1 and pVCA to the membrane. Second, frictional interactions between actin filaments may provide countering forces. It has been demonstrated experimentally (*9*, *10*) that interfilament frictional forces of sliding actin filaments can produce tens of piconewtons of force.

To test these hypotheses, we developed an extension which adds a general 2D deformable membrane model (*11*) to the popular open-source actin-network simulation tool, Cytosim (*12*). The physics of the coupling between our general membrane model and actin follows prior methods established in the literature (*13*). Altogether, our extension enables the modeling of stochastic actin networks interacting with a deformable membrane parameterized by a bending rigidity and surface tension. Utilizing the conventional features of Cytosim, which enable the definition and modeling of actin-binding proteins as agents in the simulation, we model the NPFs iLiD-SspB-pVCA and iLiD-SspB-mDia1 as membrane surface-bound beads that can nucleate actin. Arp2/3 enables the formation of branches on the actin network, while mDia1 increases the rate of barbed end elongation (*14*). The base and mDia1-enhanced polymerization rates were determined from the literature.

First, to test the hypothesis that the actin network may be forming some load-bearing configuration/scaffold, in the absence of interfilament steric interactions, we simulated the nucleation of networks in the presence of pVCA, mDia1, and both NPFs (Fig. S21A, C). We observed no deformation of the membrane in this experiment. Furthermore, visual inspection of the networks suggests that under the minimal conditions of the experiment, a highly connected branched actin network is not formed, and therefore we conclude that the bracing is not a feature endowed by force transduction through a branched actin network scaffold.

Next, to test our hypothesis that interfilament frictional forces may be the source of resistance against retrograde flow, we add to our model a repulsive potential between filaments which contributes to an effective inter-filament friction. The resulting configurations from models with the additional steric interactions are reported in Fig. S21B and D. In these simulations, we find that there is deformation of the membrane as a function of NPF (pVCA < mDia1 < pVCA-mDia1, Fig. S21D). This trend in deformation extent as a function of NPFs present is consistent with experimental observations (Fig. 4G–I, Fig. S16) which suggests that inter-filament friction can provide the bracing to support productive membrane deformation.

Taken together, our results suggest that interfilament friction, actin polymerization rate, and the specific actin network architecture influence actin-mediated membrane protrusions and explain the experimental observations.
