## Supplementary Tables for "Light-guided actin polymerization drives directed motility in protocells"

| | | Figure 2C, global actin | Figure 2F, global reversible actin | Figure 3B, local actin | Figure 3D, local reversible actin | Figure 4B, motility |
|---|---|---|---|---|---|---|
| Lipid Composition | inner leaflet | 40 mol% POPS, 60 mol% POPC | 2 mol% PE-PEG2000-benzylguanine, 98 mol% POPC | 2 mol% PE-PEG2000-benzylguanine, 98 mol% POPC | 2 mol% PE-PEG2000-benzylguanine, 98 mol% POPC | 2 mol% PE-PEG2000-benzylguanine, 98 mol% POPC |
| | outer leaflet | 40 mol% POPS, 60 mol% POPC | 100 mol% POPC | 100 mol% POPC | 100 mol% POPC | 100 mol% POPC |
| Membrane anchor | | 15 µM iLID-YFP-MARCKS | 9.8 µM iLID-YFP-SNAP | 9.8 µM iLID-YFP-SNAP | 11.75 µM iLID-YFP-SNAP | 9.8 µM iLID-YFP-SNAP |
| NPF | pVCA | 4.2 µM GST-pVCA-SspBnano-mCherry | 4.2 µM GST-pVCA-SspBnano-mCherry | 4 µM GST-pVCA-SspBnano-mCherry | 4 µM GST-pVCA-SspBnano-mCherry | 1 µM GST-pVCA-SspBmicro-mCherry |
| | mDia1 | | | | | 1 µM mCherry-SspBmicro-mDia1 |
| actin cytoskeleton | Actin | 7.5 µM (5.6% Alexa 647 labeled) | 7.5 µM (5.6% Alexa 647 labeled) | 7.5 µM (10.3% Alexa 647 labeled) | 7.5 µM (10.3% Alexa 647 labeled) | 7.5 µM (10.3% Alexa 647 labeled) |
| | Profilin | 3 µM | 3 µM | 3 µM | 3 µM | 3 µM |
| | Cofilin | 4 µM | 4 µM | 4 µM | 2 µM | 2 µM |
| | Arp2/3 | 150 nM | 150 nM | 150 nM | 150 nM | 150 nM |
| | Capping protein | 100 nM | 100 nM | 100 nM | 50 nM | 50 nM |
| Energy regeneration | ATP | 1.1 mM | 1.1 mM | 1.1 mM | 1.1 mM | 1.1 mM |
| | Creatine kinase | - | - | - | 1 µM | 1 µM |
| | Creatine Phosphate | - | - | - | 25 mM | 25 mM |
| Buffer, small molecules | Sucrose/Glucose (in/out) | 240 mM | 240 mM | 240 mM | 240 mM | 240 mM |
| | Tris-HCl (pH 7.5 at RT) | 2.2 mM | 2.2 mM | 2.2 mM | 2.0 mM | 2.2 mM |
| | HEPES-NaOH (pH 7.5 at RT) | 5 mM | 5 mM | 5 mM | 7.3 mM | 3.8 mM |
| | Imidazole (pH 8.0 at RT) | 7.2 mM | 7.2 mM | 7.2 mM | 7.3 mM | 7.0 mM |
| | KCl | 36 mM | 36 mM | 36 mM | 36 mM | 35 mM |
| | NaCl | 25 mM | 25 mM | 25 mM | 39 mM | 29 mM |
| | MgCl2 | 0.74 mM | 0.74 mM | 0.75 mM | 0.75 mM | 0.70 mM |
| | CaCl2 | 0.05 mM | 0.05 mM | 0.05 mM | 0.04 mM | 0.05 mM |
| | EGTA | 0.9 mM | 0.9 mM | 1.0 mM | 1.0 mM | 0.8 mM |
| | beta-mercaptoethanol | 1.0 mM | 1.0 mM | 1.0 mM | 1.5 mM | 1.0 mM |
| | DTT | 0.24 mM | 0.24 mM | 0.24 mM | 0.19 mM | 0.25 mM |
| | NaN3 | 0.5 mM | 0.5 mM | 0.5 mM | 0.4 mM | 0.5 mM |

| | |
|---|---|
| ATP | 1.1 mM |
| Creatin kinase | 1 µM |
| Creatin Phosphate | 25 mM |
| Sucrose/Glucose (in/out) | 240 mM |
| Tris-HCl (pH 7.5 at RT) | 2.2 mM |
| HEPES-NaOH (pH 7.5 at RT) | 3.8 mM |
| Imidazole (pH 8.0 at RT) | 7.0 mM |
| KCl | 35 mM |
| NaCl | 29 mM |
| MgCl2 | 0.70 mM |
| CaCl2 | 0.05 mM |
| EGTA | 0.8 mM |
| beta-mercaptoethanol | 1.0 mM |
| DTT | 0.25 mM |
| NaN3 | 0.5 mM |
